# On taming the effect of transcript level intra-condition count variation during differential expression analysis: a story of dogs, foxes and wolves

**DOI:** 10.1101/2022.01.24.477470

**Authors:** Diana Lobo, Raquel Godinho, John Patrick Archer

## Abstract

The evolution of RNA-seq technologies has yielded datasets of high scientific value that are often generated as condition associated biological replicates within differential expression studies. As the number of replicates increase, so to does confidence in identifying differentially expressed transcripts. With rapidly expanding RNA-seq data archives there is opportunity to augment replicate numbers when conditions of interest overlap at an inter-study level. Despite correction procedures for estimating transcript abundance, a remaining source of error is transcript level intra-condition count variation; as partially indicated by the disjointed results between differential expression analysis tools. Such variation is amplified at an inter-study level. Here, we present TVscript, a tool that removes reference-based transcripts associated with intra-condition variation above specified thresholds. With this tool we explore the effects of removing transcripts associated with varying degrees of intra-condition variation on differential expression analysis. This is done in relation to inter- and intra-study datasets representing brain samples of dogs, wolves and foxes (wolves *vs.* dogs and aggressive *vs.* tame foxes). We demonstrate that 20% of the transcripts identified as being differentially expressed are associated with high levels of intra-condition variation. This is an over-representation relative to the reference set. As transcripts harbouring such variation are removed from the reference prior to differential expression analysis a discordance from 26 to 40% in the lists of differentially expressed transcripts is observed when compared to those obtained using the non-filtered reference. For our data, the removal of transcripts possessing intra-condition variation values within (and above) the 97^th^ and 95^th^ percentiles, for wolves *vs.* dogs and aggressive *vs.* tame foxes, maximized the detection of differentially expressed transcripts as a result of alterations within gene-wise dispersion estimates. Through this analysis the support for seven genes with potential for being involved with selection for tameness is provided. TVscript is available at: https://sourceforge.net/projects/tvscript/.

## Introduction

Developments in RNA-seq technology have revolutionized transcriptomic studies by allowing for a rapid hi-resolution view of transcript expression [1]. In a typical RNA-seq experiment, transcript expression profiles are estimated for each sample using a metric based upon the number of sequenced reads associated with each transcript within a reference set [2–8]. Condition dependent expression profiles can then be used in order to identify which transcripts are differentially expressed [9, 10]. A challenge arises due to sources of variation within expression profiles that are independent of, or partially overlapping with, the condition of interest [2,11–13]. The inclusion of biological replicates reduces the effect of such noise [14, 15],, and it has been demonstrated that sufficient replicate numbers outweigh sequencing depth in terms of increasing the accuracy within differential expression experiments [14, 16]. In studies not involving highly controlled isolated environments, RNA-seq data from the rapidly growing repertoire of published works can be incorporated [17–19], if data from a matching condition to that being studied is available. This effectively increases the number replicates although variability can be amplified [20, 21].

Differential expression tools compute a statistical significance for each transcript, based upon the abundance estimates within a condition, that reflect the possibility of that transcript being differentially expressed [9,10,22]. To reduce the effect of intra-condition variation across biological replicates on the estimation of abundance several methods have been proposed including ALDEx2 [23], EDASeq [24] and PEER [25]. In addition to these, and more generally applied, are the abundance estimation techniques implemented within established differential expression tools such as DESeq2 [9] and EdgeR [10]. However, when methods are compared, relative to the final sets of transcripts identified as being differentially expressed, variable results are observed [15,23,26–29]. This is an indication that the problem of intra-condition variation relative to the detection of differentially expressed transcripts using RNA-seq data has not been completely resolved. Furthermore, there is no consensus on the best approach to use [30].

Here we explore the effects that individual transcripts associated with high levels of intra-condition abundance variation have on the end results of differential expression analysis using the tool DESeq2 [9]; a tool that is well established and that has consistently demonstrated reliability in identifying differentially expressed transcripts [27,30,31]. Our aim is to investigate the possibility of whether or not the removal of transcripts, harbouring the highest levels of intra-condition variation, from the reference set used during differential expression analysis can produce sets of differentially expressed transcripts that display an increased level of confidence. The latter being achieved through either: (a) the direct removal of transcripts previously identified as being differentially expressed, but whose expression patterns are ambiguous, or (b) the indirect addition, or removal, of transcripts to, or from, those previously identified as being differentially expressed as a consequence of alterations in p-adj values. The latter being associated with shifts in the distribution of intra-condition variation, following the removal of transcripts harbouring the highest levels of such variation. A by-product of this is the explicit quantification of the level of intra-condition abundance variation present within the final lists of differentially expressed transcripts.

To aid this exploration we present TVscript, a tool, for the identification of transcripts above user-specified levels of intra-condition normalized count variation, the latter being strongly associated with transcript abundance estimation [4–8]. As input TVscript requires one file per replicate that contains the per transcript read counts following the mapping of reads from the replicate to a common reference set. As output TVscript produces a set of corresponding count files that are absent of transcripts harbouring normalized intra-condition count variation higher than that associated with a user specified percentile. These updated count files can be subsequently used within the differential expression tool of choice, in our case Deseq2 [9]. Through multiple iterations of differential analysis following filtering at varying thresholds, the effects of transcripts associated with high intra-condition variation can be explored.

Using TVscript we explore the effects of intra-condition read count variation within two distinct case-studies, involving intra and inter-study datasets. In these case-studies differential expression patterns arising from data derived from brain samples of dogs, wolves, and foxes, generated in the scope of domestication experiments, are compared at varying thresholds of intra-condition normalized read count variation exclusion. Our aim here was twofold, as in addition to exploring the effects of transcripts harbouring high levels of intra-condition normalized read count variation on the outcome of differential expression analysis, we also had an interested in understanding whether or not there were genes commonly up or down regulated within the brain of both forms of domestic canids (dogs and tame foxes), but simultaneously not so within their “wild/aggressive” counter parts (wolves and wild foxes). Such genes are candidates for being associated with tameness. Domestic dogs present marked behaviour differences from wolves, their wild ancestors, due to the evolution of unique social cognitive capabilities [32–34]. Tame red foxes resulted from deliberated selection against fear and aggression over several generations of cross-breeding [35] and they present several behavioural and pheno-typical traits that resemble those found in dogs [36–38].

## Materials & Methods

### RNA-seq datasets

To validate the performance of TVscript, we used both intra and inter-study datasets. At an intra-study level, we combined publicly available RNA-seq data from brain tissues of 12 tame and 12 aggressive red foxes (Table S1) generated within the same study [37]. For the inter-study case, we combined multiple publicly available RNA-seq datasets from several dogs and six wolves [34,39–41], also derived from brain tissues (Table S1). All the samples were downloaded from the National Center for Biotechnology Information (NCBI) and the European Bioinformatics Institute (EMBL-EBI), covering a wide range of ages, both sex as well as multiple replicate and sequencing strategies (Table S1). We selected these specific case studies because, firstly we were interested in evaluating the effects of TVscript both at intra and inter-study levels and the domestic dog, being a model organism, has an available high-quality reference transcriptome as well as several high-quality RNA-seq datasets generated across different studies; while secondly, we sought to perform a brief exploratory inter-study scan to investigate if the domestication of both dogs and foxes has resulted in the co-expression of a set of common brain genes, relative to their “wild/aggressive” type, since behavioural modifications are considered to have been the first target in domestication [42].

Reads from all samples were mapped to the dog reference transcriptome [41], which contained 26,107 annotated transcripts (Ensembl CanFam3.1, release 92) using Bowtie v.2.3.4.1 [43]. Reads were not mapped to the dog genome as our aim was to explore the effects of intra-condition variation on a predefined set of reference transcripts, and not infer novel transcripts from these previously published datasets. Next, we used the *pileup.sh* script from the BBmap package [44] to obtain per transcript abundance estimates (measured by the number of mapped reads to the corresponding transcript) in each sample. Read counts from technical replicates of “Dog_8” and “Dog_9” were averaged and merged into one file (Table S1), while read counts from the two biological replicates of “Dog_7” were treated separately.

### Software

TVscript requires as input: (1) multiple files containing per transcript read counts (one per sample), (2) a file containing the lengths of the transcripts that the reads were mapped to, (3) a percentile threshold value for intra-condition varation and (4) a configuration file that indicates the locations all files as well as the condition allocations of the count files in (a). An example configuration file along with further details is available from the SourceForge project page. To explore the effects of intra-condition variation on differential expression analysis within our data, we ran TVscript multiple times within each of our case studies (wolves *vs.* dogs and aggressive *vs.* tame foxes) using a range of percentile thresholds for variance (described within the next section). The steps that TVscript implements to identify transcripts associated with high levels of disjointed read counts are (see Fig. 1 for a workflow): i) each input dataset, containing count values from a particular sample, is allocated to either condition A or B, as indicated within the configuration file; ii) counts are normalized by dividing them by the length of the corresponding reference transcript and by the sum of all counts for that sample; (iii) for each reference transcript (t), the absolute pairwise differences between normalized read counts across all samples within condition A are calculated; (iv) the corresponding variances are calculated; (v) steps (iii) and (iv) are repeated for condition B; (vi) variance scores from each condition are placed in ascending order and associated with corresponding percentiles; (vii) reference transcripts are removed (or filtered) if their variance score is above that associated with the user specified percentile threshold; (viii) raw read counts associated with the remaining transcripts are outputted into separate files that correspond to each input dataset. As to avoid overwriting the original count files, the names of updated count files are specified within the configuration file. These updated count files can be used as input for differential expression analysis software such as DESeq2. TVscript is implemented in Java programming language and runs on all operating systems with installed Java Runtime Environment v.8.0 or higher. It is available under the GNU General Public License v3.0, along with source code, usage instructions and sample data, from our SourceForge project page: https://sourceforge.net/projects/tvscript/.

**Figure 1:**
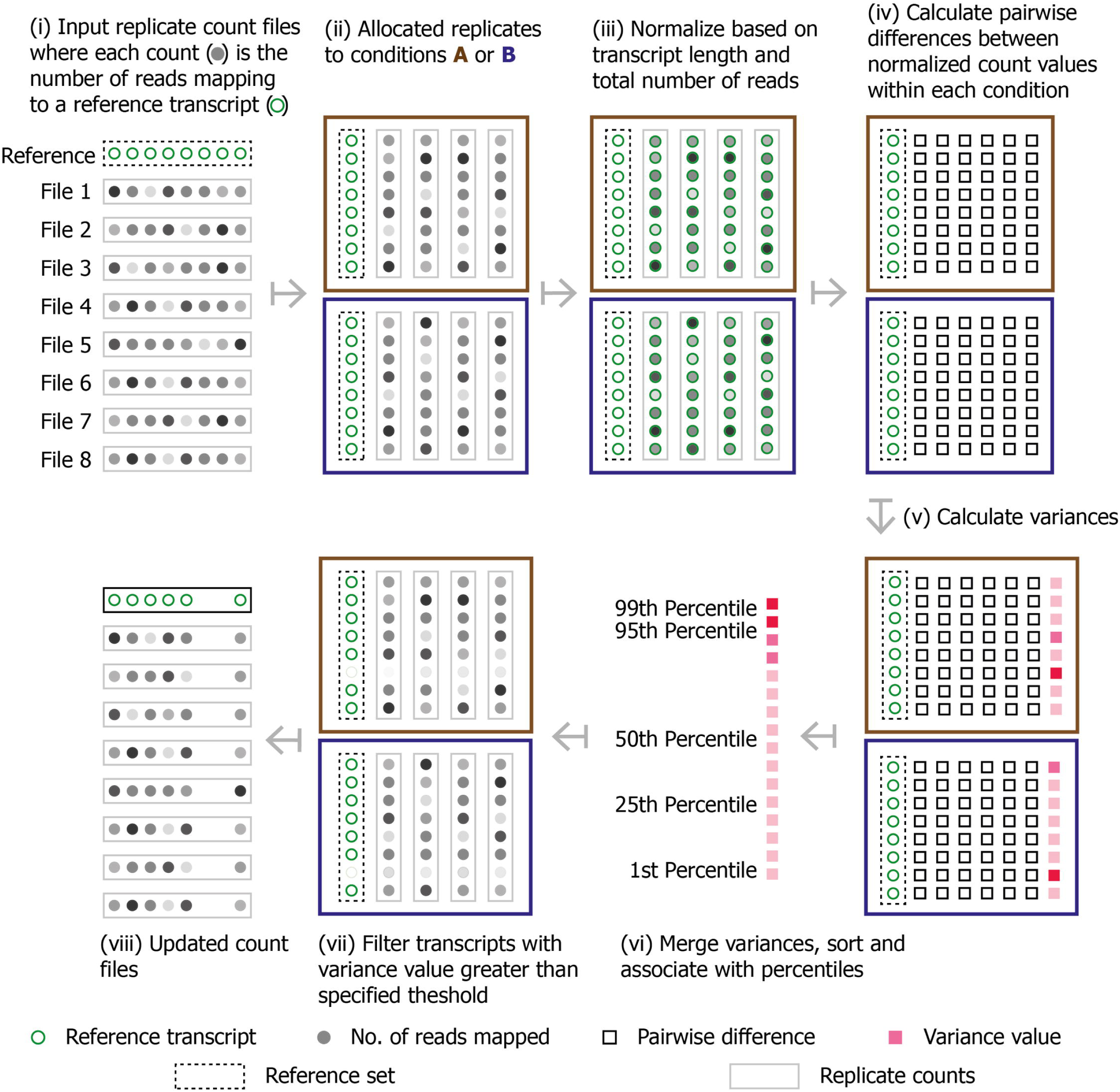
Diagramatic overview of how reference based transcripts are removed. Steps (i) to (viii) indicate actions taken. Reference transcripts (green circles) are showen for diagramtic purposes in order to highlight how read counts (grey circles) across replicates are treated for each transcript independently. Read counts are grouped into individual files (gray rectangles) in accordence to replicate. These files are grouped into one of two conditions (blue and brown boxes). Remaining keys are indicated at the bottom of the figure.

### Exploring the removal of transcripts associated with high intra-condition variation

For each case study, we ran TVscript using variance thresholds corresponding to the 70^th^ up to the 90^th^ percentiles (in steps of five), and to the 91^st^ up to the 99^th^ (in steps of one). Steps of one were used in the latter as to allow for transcripts associated with the highest levels of intra-condition variation to be explored in more detail. For each threshold value, only transcripts with variance below that value were maintained. During each run, we recorded the number and IDs of all transcripts that were removed so that they could be cross-compared.

Within each case study, following each run, we used the updated count files produced to perform differential expression analysis using DESeq2 v.1.22.2 [9]. Transcripts with p-adj < 0.05 (corrected by the Benjamini and Hochberg method) were considered to be differentially expressed. DESeq2 identifies differentially expressed transcripts by estimating gene-wise dispersions and applying shrinking methods to model counts and thus effectively normalize for individual outliers [31]. Distributions of gene-wise dispersions following normalization are conveniently accessible and provide a good metric to visualize the effects of removing transcripts associated differing filter levels. Differential expression analysis using the original non-filtered datasets was also performed. To visualize the overall effects of experimental covariates in each case study, we performed a principal component analysis (PCA) using the *plotPCA* function from DESeq2 with non-filtered normalized count data.

### Evaluation metrics

To evaluate TVscript we used three metrics that when combined quantify the overall impact of intra-condition variation on downstream differential expression analysis. The metrics were: i) number of ambiguous positives within transcripts identified as being differentially expressed in the non-filtered datasets; ii) distributions of dispersion estimates and outliers in differential expression analysis for non-filtered and all filtered datasets; and iii) discordance in the list of differentially expressed transcripts between non-filtered and filtered datasets (selected percentiles: 97^th^, 95^th^, 90^th^).

#### (i) Ambiguous positives

We identified transcripts appearing as being differentially expressed when using the non-filtered datasets as input to DESeq2 that were associated with the top 10 levels of intra-condition variation (above the 90^th^ percentile threshold value). These we designated as ambiguous positives. Small numbers of these, relative to the overall number of identified differentially expressed transcripts would indicate that TVScript is having little direct effect on lists of identified differentially expressed genes.

#### (ii) Distributions of dispersion estimates

For the non-filtered and all filtered input datasets (70^th^ up to the 90^th^ percentiles in steps of five and to the 91^st^ up to the 99^th^ in steps of one) we calculated correlation coefficients (r^2^) using a linear regression analysis in R, between dispersion estimates and the mean of normalized counts, both the latter calculated by DESeq2 during differential expression analysis. Dispersion is inversely related to the mean, as lower mean counts are affected by variation to a higher degree. If a stronger correlation is seen for the filtered input datasets then this would suggest that the distribution used to model differential expression could be more reliable in relation to identifying differentially expressed transcripts. In addition to this, we retrieved the number of outliers detected by DESeq2, expecting a decrease after each filtering step. Outliers are recognized by the DESeq2 as the points with extremely high dispersion values that cannot by shrunk towards the fit curve. This was performed independently for both case studies.

#### (iii) Discordance lists of differentially expressed transcripts between filtered levels

We calculated the proportion of discordance between lists of differentially expressed transcripts produced when using non-filtered and filtered datasets at the 97^th^, 95^th^ and 90^th^ percentile threshold values. Two types of observed discordances relative to the non-filtered list were considered: (a) transcripts that were lost directly due to filtering or indirectly due to p-adj values no longer being significant, and (b) transcripts that were added due to alterations in p-adj values. Quantifying the nature of these discordances provides insight into the general consistency of genes identified as being differentially expressed across varying filter thresholds. To visualize the overlap between the non-filtered and filtered lists we used the *VennDiagram* v.1.6.20 package in R [45].

### Gene annotation and gene family analysis

For each case study, differentially expressed transcripts obtained using the non-filtered and filtered datasets were matched to the correspondent gene ID. This was done with the R package BioMart [46] using the Ensembl Gene database (version 94). To begin to identify gene families that displayed similar regulation in both dogs and tame foxes, i.e. relative to the “wild/aggressive” type, we grouped up and down regulated genes into gene families. Genes within these families were then classified according to whether they were unique to dogs or tame foxes or shared between the two. Within each case study, a gene family was only considered if all the associated genes agreed in relation to their direction of expression (up or down regulation).

## Results

### Mapping success

Mapping of the 44 datasets corresponding to dogs and wolves against the dog reference transcriptome revealed an average success of 60% and 58% respectively, in terms of the number of mapped reads (Fig. S1). Similar values between wolves and dogs were expected, given their recent divergence of ∼23,000 years ago [47]. Comparable proportions of reads failing to map (∼40%) have been previously reported for dog brain samples [39] and are most likely associated with i) novel genes; ii) regions that are not translated despite being transcribed; iii) contamination with genomic DNA; and iv) uncharacterized chimeras and other artefacts within reference sets resulting from library preparation during sequencing [48] and various assembly errors [49]. For the fox datasets, an average of 50% of reads mapped to the dog reference transcriptome (Fig. S1). This lower percentage of mapped reads, relative to the dog and wolf datasets, could be expected due to an increased genetic divergence from dogs (∼10 mya) [50] together with the other aforementioned factors.

### Exploring the removal of transcripts associated with high intra-condition variation

No significant difference existed between the overall distributions of the per-transcript intra-condition variation values for wolf and dog samples (Wilcoxon-test, p-value < 0.198, Fig. 2a). The PCA based on the entire set of normalized non-filtered counts, revealed that the wolf samples were more aggregated than dog samples (Fig. 2c). For aggressive and tame fox samples, we observed a significant difference (Wilcoxon-test, p-value < 2.2e^-16^, Fig. 2b) between the distributions of the per-transcript intra-condition variation values, most likely resulting from an increased intra-condition variability within tame fox samples. In particular, we found five samples that were differentiated from the remaining seven in the PCA (Fig. 2d), with 80% variance being explained by this clustering in PC1.

**Figure 2:**
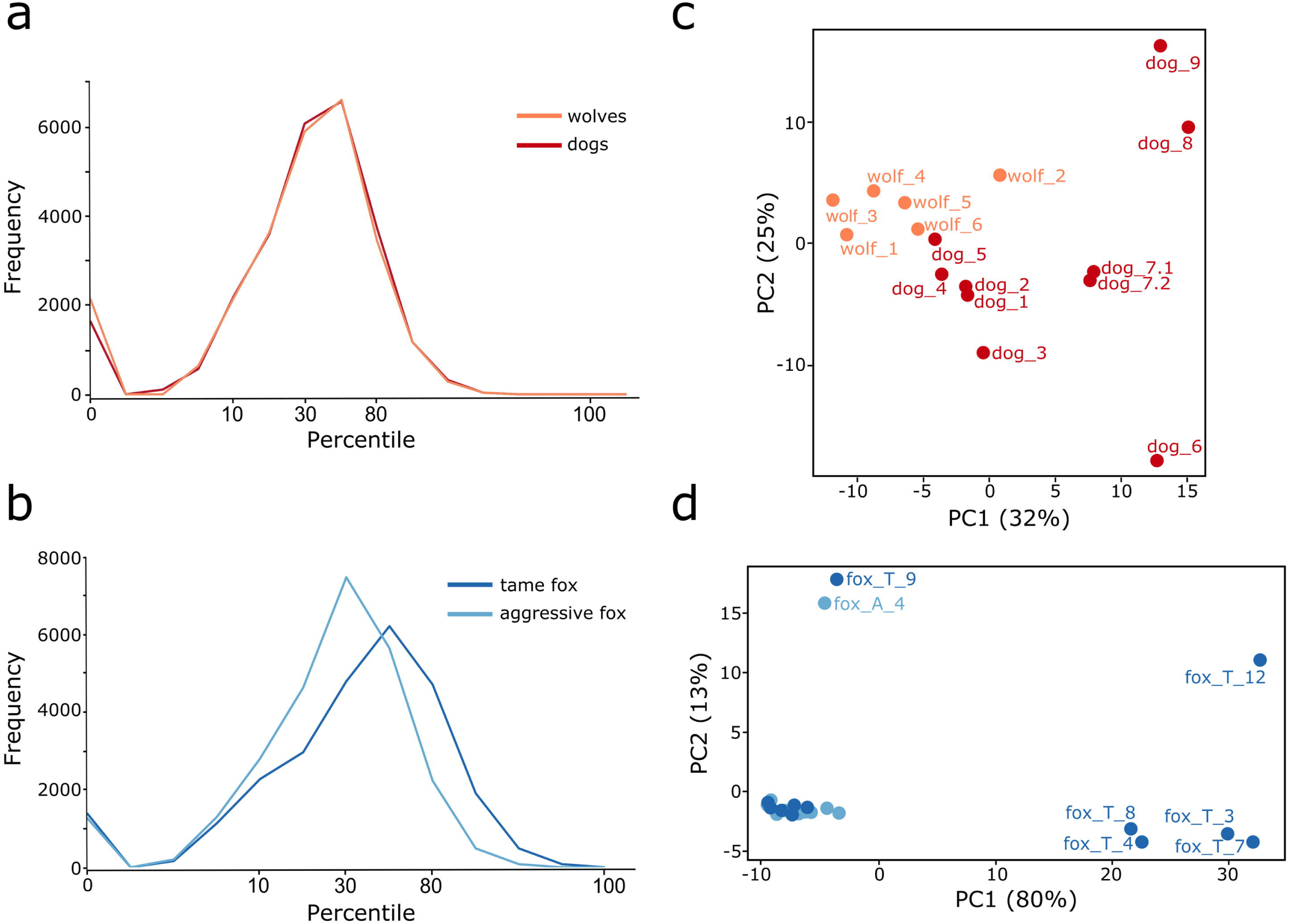
Characterization of intra-condition variation. Percentile range of intra-condition variation scores (x-axis) observed prior to filtering, across both case studies, a) wolves (orange) and dogs (red); b) tame (dark blue) and aggressive (light blue) foxes. PCA plots based on normalized and non-filtered count data of the individual datasets comparing c) wolf and dog, and d) tame and aggressive fox samples. In the latter only individual samples that were positioned within a distant cluster are labelled with the sample ID.

Prior to differential expression analysis, for each case study (wolves *vs.* dogs and aggressive *vs.* tame foxes), TVscript was used to remove transcripts in accordance with a series of intra-condition variance thresholds as previously described (Fig. 3a, 3b; Table S2). Initially, for wolves and dogs 184 transcripts (out of the 26,107) associated with high intra-condition variation (99^th^ percentile and above) were removed, while for the aggressive and tame fox samples, 235 transcripts were removed. The number of transcripts removed was higher for the fox samples than for those of wolf and dog, reflecting the higher intra-condition variability present. Combined across the top 10 levels of intra-condition variation 12% (n = 3134) and 14.89% (n = 3888) of the reference transcripts were removed in wolf/dog datasets, and aggressive/tame fox datasets, respectively (Table S2).

**Figure 3:**
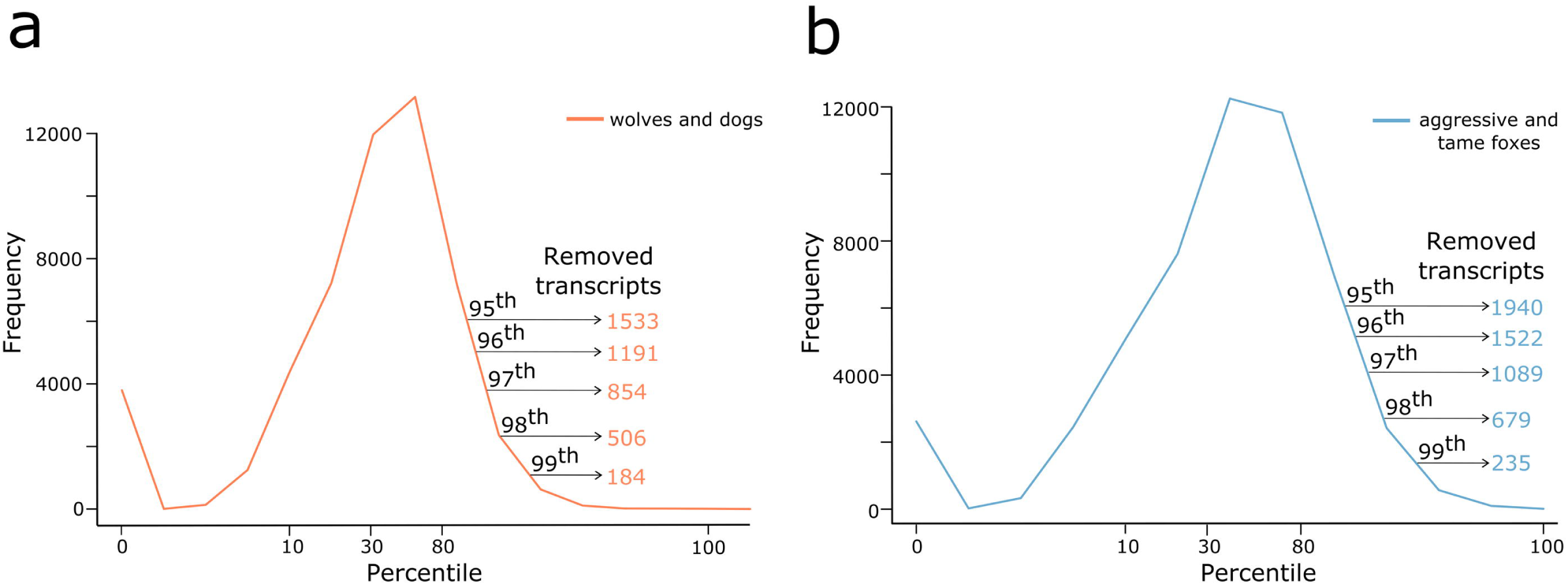
Removal of transcripts above specified levels of intra-condition variation. Percentile range of combined intra-condition variation scores (x-axis) present in each case study, a) wolves and dogs; b) tame and aggressive foxes. The number of transcripts removed in the top five percentiles (from the 95^th^ to the 99^th^) are presented in each panel.

### Differential gene expression analysis

Using non-filtered datasets as input to DESeq2, 430 differentially expressed transcripts were identified between wolves and dogs (Fig. 4a; Table S3). Of those, 259 were up regulated, while 171 were down regulated in dogs. Between aggressive and tame foxes, 651 differentially expressed transcripts were identified (Fig. 4a; Table S4), of which, 532 and 119 were up and down regulated, respectively, in tame foxes. Post filtering, within the first ten steps of size one from the 99^th^ down to the 90^th^ percentiles, the number of differentially expressed transcripts identified peaks at the 97^th^ (n = 430; up = 255, down =175) and the 95^th^ percentiles (n = 730; up = 607, down = 123) in dogs and tame foxes (Fig. 4a), respectively. This indicates that for these data the removal of the 3% (n = 854) and 5% (n = 1940) of transcripts associated with the highest levels of intra-condition variation maximized the detection of differentially expressed transcripts.

**Figure 4:**
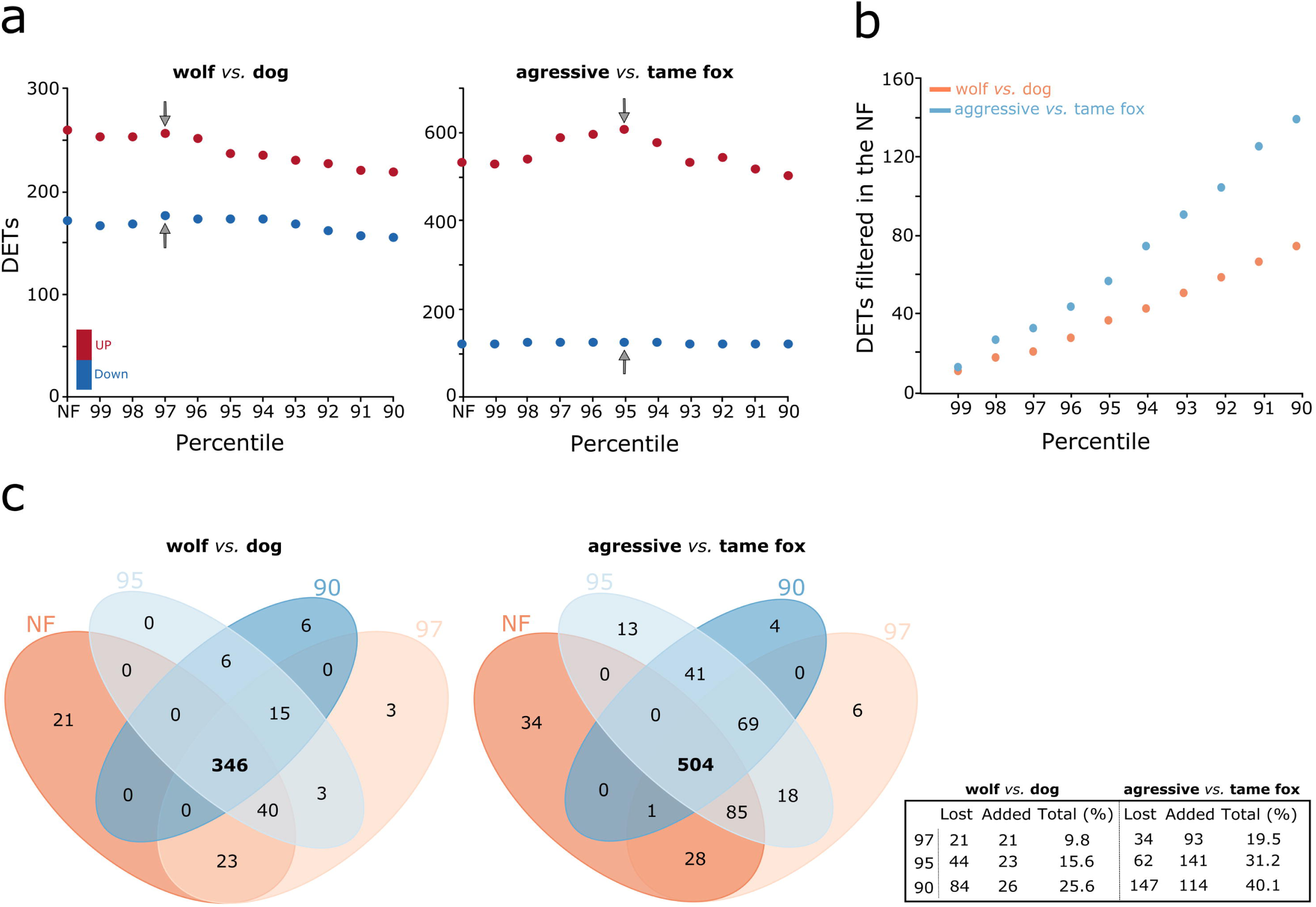
Effects of removing transcripts above specified levels of intra-condition variation on differential expression analysis. a) Number of differentially expressed transcripts (DETs) identified using non-filtered (NF) and filtered datasets, based on the top 10 percentiles (99^th^ to the 90^th^), for both case studies. Up and down regulated transcripts are represented by red and blue dots respectively. Gray arrows identify the selected thresholds for which the results of subsequent correspondng differential expression analysis were used for the identification of candidate transcript associated with tameness within each case study. b) Number of transcripts identified as differentially expressed within the non-filtered datasets that were associated the highest levels of intra-condtion variation (99^th^ to 90^th^) within both case studies, wolves and dogs (orange dots), and tame and aggressive foxes (blue dots); c) Venn diagrams representing the number of overlapping differential expressed transcripts found following differential expression analysis using non-filtered datasets and filtered datasets (97^th^, 95^th^ and 90^th^ percentiles), within each case studies. The associated table provides information about the number of differential expressed transcritps lost/added following each filter step in relation to the non-filtered dataset as well as the the percentage of total discordance.

### Evaluation Metrics

#### (i) Ambiguous positives

Of the transcripts that appeared as being differentially expressed, when using non-filtered datasets as input to DESeq2, 17.44 (n = 75) and 21.51% (n = 140) were associated with high intra-condition variation (above the 90^th^ percentile threshold) within the wolves *vs.* dogs and aggressive *vs.* tame foxes case studies respectively (Fig. 4b). These were transcripts that we considered as ambiguous positives and the average across both case studies was 19.45%. The number of was higher within the fox datasets where elevated variability among samples was observed, suggesting that differences within intra-condition read counts could of had an influencing the final outcome of identified differentially expressed transcripts.

#### (ii) Distributions of dispersion estimates

Within both case studies the regression analysis indicates that removing transcripts associated with high levels of intra-condition variation improved correlation coefficients in relation to those from the non-filtered datasets (Fig. 5a, 5b; Table S5). Associated with the elevated levels of variation observed within the fox datasets there was a better fit within the wolves *vs.* dogs (r^2^ > 0.7) comparison when compared to the aggressive *vs.* tame foxes (r^2^ > 0.4). For the latter three was visible elevation in the number of dispersed points around the line of best fit. With the removal of transcripts associated with the highest levels of intra-condition variation a reduction in the number of outliers within both case studies was also observed (Fig. 5a, 5b; Table S5).

**Figure 5:**
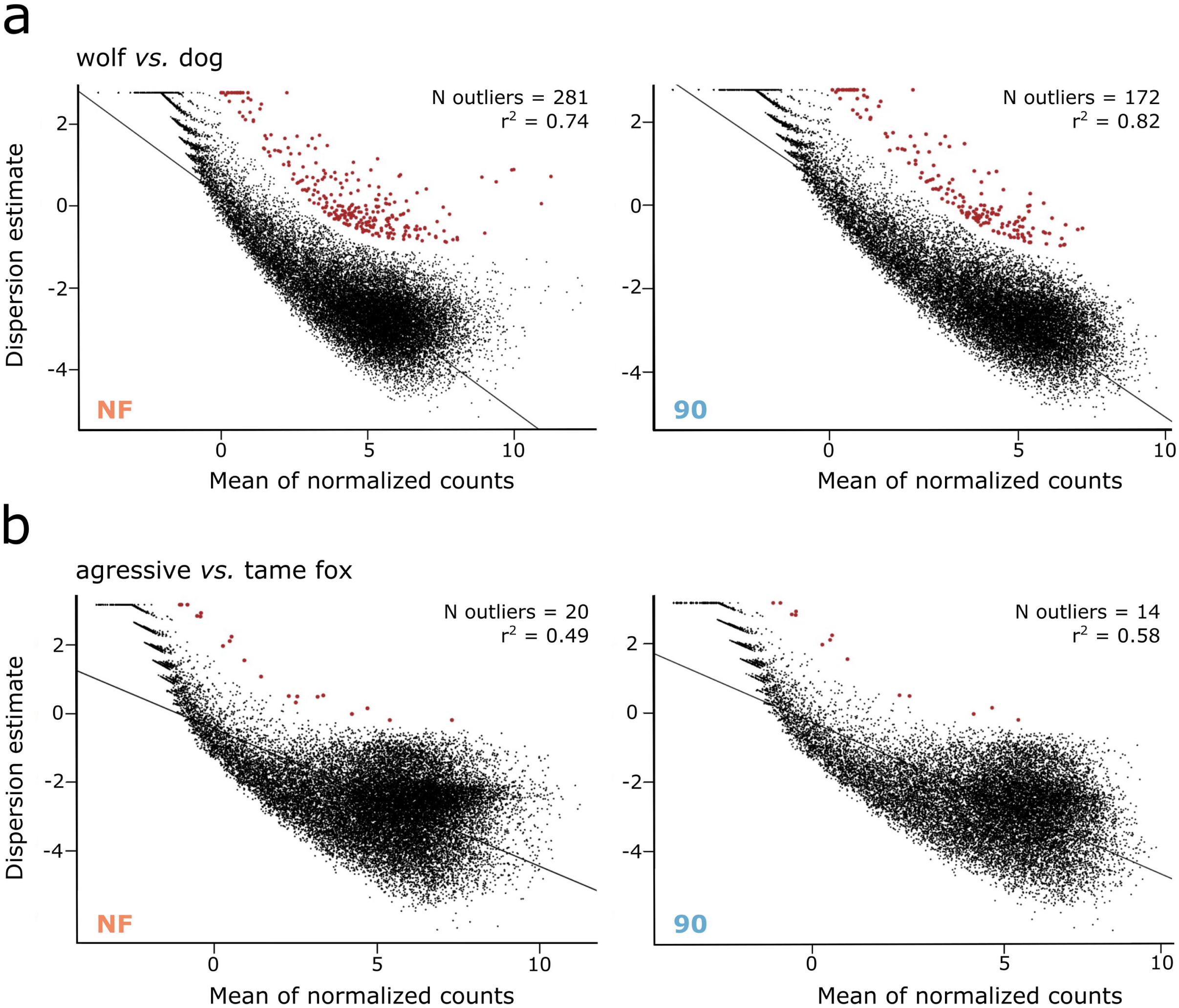
Distribution of dispersion estimates. Plots of final dispersion estimates for both case studies, a) wolves and dogs; b) tame and aggressive foxes, calculated using DESeq2 for the non-filtered (NF; orange) and 10% filtered datasets (90^th^; blue). Each black dot represents a single transcript, and red dots represent outliers. The number of outliers and correlation index (r^2^) are displayed in the top right corner of each pannel. Both x and y-axis are transformed into a logarithm scale. The line in each graph corresponds to the regression analysis between the mean of normalized counts and dispersion estimates.

#### (iii) Discordance lists of differentially expressed transcripts between filtered levels

Within the wolves *vs.* dogs case study, from the 430 differentially expressed transcripts identified when using non-filtered data as input to DESeq2, 346 were maintained when compared to differentially expressed genes obtained using input data filtered at the 97^th^, 95^th^, and 90^th^ percentile threshold values (Fig. 4c and Table S3). 26 transcripts were added as differentially expressed following filtering (Fig. 4c). Overall for this case study between the differentially expressed transcripts identified using filtered and non-filter input data there was a discordance of 25.58% (n = 110). For the second case study, aggressive *vs.* tame foxes, 504 out of the 651 differentially expressed transcripts identified using the non-filtered input were maintained when compared to those using input data filtered at the 97^th^, 95^th^, and 90^th^ percentile threshold values, with up to 114 being added following filtering (Fig. 4c and Table S4). This time the overall level of discordance was 40.09% (n = 261) between the differentially expressed transcripts identified using filtered and non-filter input data. Importantly, in both case studies, we found that the added transcripts were consistently maintained across the three filter levels.

### Candidate genes and gene families

By performing annotation using the filtered datasets where the number of differentially expressed transcripts was maximized (3% and 5%, in dogs and tame foxes, respectively), we found 21 gene families in common among the up-regulated genes in dogs and tame foxes. These 21 gene families contained 50 genes (Table 1), of which 19 were exclusive to dogs while 24 were exclusive to tame foxes. The remaining seven genes (RGR, CHRNA5, SQLE, ARHGAP25, ITGA7, MYO7A and TRIB2), were found to be commonly up regulated in both dogs and tame foxes.

**Table 1:** Shared genes and gene families (Up regulation). List of the gene families, and shared genes, that were commonly up regulated in dogs and tame foxes. The number, and name, of the genes within each gene family are provided, with the corresponding log2fold-change values in brackets for each species. Within each family, single genes were charecterized as shared between dogs and tame foxes, or as exclusively to each of the two groups. When more than one transcript for a specific gene was present, all the log2FC values are reported.

| Gene Family | Group | N of genes | Gene name and log2FC value |
| --- | --- | --- | --- |
| Retinal G protein-coupled receptor | Shared | 1 | **RGR** (2.10 in dogs, 0.78 in tame foxes) |
| Cholinergic receptor nicotinic alpha | Shared | 1 | **CHRNA5** (1.1 in dogs, 0.4 in tame foxes) |
| Squalene epoxidase | Shared | 1 | **SQLE** (0.54 in dogs, 0.31 in tame foxes) |
| Rho GTPase activating protein | Shared | 1 | **ARHGAP25** (0.86 in dogs, 0.72 in tame foxes) |
|  | Tame fox | 2 | ARHGAP4 (0.64); ARHGAP30 (0.57) |
| Integrin alpha subunits | Dog | 3 | ITGA6 (1.25, 1.24); ITGA8 (1.14, 0.90); ITGAX (0.97) |
|  | Tame fox | 1 | ITGAL (0.73) |
|  | Shared | 1 | **ITGA7** (0.76 in dogs, 0.46 and 0.49 in tame foxes) |
| Myosin | Dog | 1 | MYO3A (1.12) |
|  | Tame fox | 3 | MYOZ1 (1.53); MYO1F (0.93); MYO1C (0.47) |
|  | Shared | 1 | **MYO7A** (0.82 in dogs; 0.41 in tame foxes) |
| Tribbles pseudokinase | Tame fox | 2 | TRIB1 (0.94); TRIB3 (0.78) |
|  | Shared | 1 | **TRIB2** (0.61 in dogs; 0.2 in tame foxes) |
| EF hand calcium binding | Dog | 1 | EFCAB1 (2.59) |
|  | Tame fox | 1 | EFCAB2 (0.46) |
| Transcription factor | Dog | 1 | TCF23 (2.04) |
|  | Tame fox | 1 | TCF19 (0.63) |
| Adhesion G protein-coupled receptors | Dog | 1 | ADGRG6 (1.45) |
|  | Tame fox | 1 | ADGRG1 (0.57) |
| Patatin Like Phospholipase Domain | Dog | 1 | PNPLA4 (1.41) |
|  | Tame fox | 1 | PNPLA7 (0.59) |
| SRY-box | Dog | 1 | SOX6 (1.26) |
|  | Tame fox | 2 | SOX17(0.84); SOX10 (0.66) |
| Hyaluronan and proteoglycan link protein | Dog | 1 | HAPLN1 (1.15) |
|  | Tame fox | 1 | HAPLN3 (0.70) |
| Serine/threonine kinase | Dog | 2 | STK17A (1.15, 1.14); STK32A (1.10) |
|  | Tame fox | 1 | STK40 (0.57) |
| Potassium channels | Dog | 1 | KCTD16 (0.98) |
|  | Tame fox | 1 | KCTD15 (0.72) |
| Podocalyxin like | Dog | 1 | PODXL (0.95, 0.84) |
|  | Tame fox | 1 | PODXL2 (0.70, 0.69, 0.67) |
| ATP binding cassette subfamily B | Dog | 1 | ABCB1 (0.93) |
|  | Tame fox | 1 | ABCB9 (0.52) |
| Zinc finger DHHC-type | Dog | 1 | ZDHHC15 (0.75) |
|  | Tame fox | 1 | ZDHHC1 (0.70) |
| Sushi domain | Dog | 1 | SUSD1 (0.68) |
|  | Tame fox | 2 | SUSD3 (0.79); SUSD6 (0.47) |
| TBC1 domain family | Dog | 1 | TBC1D5 (0.54) |
|  | Tame fox | 1 | TBC1D7 (0.27) |
| Mitogen-activated protein kinase kinase kinases | Dog | 1 | MAP3K5 (0.51) |
|  | Tame fox | 1 | MAP3K11 (0.76) |

In addition, we also found three gene families, containing four genes, simultaneously down regulated in both groups (Table 2). Two of these genes (STMND1 and OASL) were shared between dogs and tame foxes, while the other two were unique to each group. The same analysis performed using the non-filtered datasets revealed similar results (Table S6), although the RGR gene family which included a shared gene between dogs and tame foxes, was lost. This gene was not differentially expressed in the non-filtered fox dataset, representing an example of genes added as differentially expressed after filtering.

**Table 2:** Shared genes and gene families (Down regulation). List of the gene families, and shared genes, that were commonly down regulated in dogs and tame foxes. The number, and name, of the genes within each gene family are provided, with the corresponding log2fold-change values in brackets for each species. Within each family, single genes were characterized as shared between dogs and tame foxes, or as exclusively to each of the two groups. When more than one transcript for a specific gene was present, all the log2FC values are reported.

| Gene Family | Group | Number of UE | Gene name and log2FC value |
| --- | --- | --- | --- |
| Stathmin domain | Shared | 1 | STMND1 (-1.18 in dogs, -0.53 in tame foxes) |
| Oligoadenylate synthetase like | Shared | 1 | OASL (-0.41 in dogs, -0.52 in tame foxes) |
| Heat shock protein family B | Dog | 1 | HSPB8 (-0.70) |
|  | Tame fox | 1 | HSPB11 (-0.32) |

## Discussion

Studies involving RNA-seq data often rely on the identification of one, few or many differentially expressed transcripts in order to draw conclusions about biological pathways or in relation to general transcriptome function and evolution. The explicit quantification of intra-condition read count variation associated with such transcripts is important for maintaining the context of ambiguity that may exist following differential expression analysis. This is especially true given the growing ability to base highly informative studies around archived transcriptomics datasets at an inter-study level. Here, we developed a method to quantify intra-condition variation for each individual transcript within the reference set and that can be used to identify and remove reference-based transcripts harbouring intra-condition variation a above specified thresholds. By applying the method across varying thresholds of intra-condition variation, we explored the effects of reducing such variation that the well-established differential expression analysis tool, deseq2, is required to accommodate. Our case study data was from brain samples of dogs, wolves and foxes and obtained from both inter- and intra-study levels.

By associating all transcripts with an intra-condition variation value we found that following differential expression analysis, using non-filtered input data, on average nearly 20% of the transcripts identified as being differentially expressed across both case studies contained levels intra-condition variation equal to or above the 90^th^ percentile value of the total distribution (Fig. 4b). This was higher than the relative proportion of such transcripts with the reference set in general, where 12% and 14.89% of transcripts possessed intra-condition variation above this level for the wolf/dog and the aggressive/tame fox case studies respectively (Fig. 3a, 3b; Table S2). This indicates that within these case studies, transcripts associated with higher intra-condition variation have a tendency to being identified as differentially expressed. When transcripts possess large amounts of such variation, some ambiguity in their identification as being differentially expressed is inevitable, since reliable expression patterns for at least one of the two conditions being compared have not been fully established; even if statistical correction is applied. This likely partially explains the level of discordance between various differential expression tools available [15,23,26–29], for which no consensus on the best approach to apply exists [30]. However, more importantly, when such transcripts are used for drawing biological conclusion, the context of this uncertainty must be maintained.

We then explored what effects the removal of transcripts associated with intra-condition variation, at varying threshold levels, would have on the gene-wise dispersion estimates, used by DESeq2. Within both case studies, as such transcripts were increasingly removed from input datasets prior to differential expression analysis, the correlation between the mean of normalized counts and dispersion estimates increased, and the number of outliers identified decreased (Fig. 5a, 5b; Table S5; Fig. S2). Relative to this, and to the differences within the lists of differentially expressed transcripts identified when using non-filtered and filtered datasets (at the 97^th^, 95^th^, and 90^th^ percentiles), transcripts were not simply removed because of physical exclusion from the input data, they were also removed, and added, as a result of the effects of removing intra-condition variation from the general gene-wise dispersions applied. We found high discordance rates, reaching 40% within the aggressive *vs.* tame fox case study (Fig. 4c and Table S4), between the list of differentially expressed transcripts within the non-filtered and filtered datasets, thus revealing how dependent the identification of differentially expressed transcripts is on the accuracy of gene-wise dispersion estimates used; these in turn being affected by transcripts associated with high intra-condition variation.

Such variation at an inter, and to a lesser extent intra, study level can arise from a range of sources including i) biological differences between samples such as age, sex, diet, and health; ii) *in silica* error involving assembly tools producing poorly understood chimeras within the reference transcriptome [51]; iii) ambiguities in read mapping to such references [52]; iv) normalization of count data derived from such mapped reads [53]; and v) including *in vitro* error during library preparation protocols [54, 55]. Although we have used DESeq2 within our study, the results are in no way indicative of the performance levels of Deseq2 as no benchmarking was involved. Indeed we used Deseq2 as such benchmarking studies, as discussed within the introduction section, indicated that it is a reliable tool for differential expression analysis. The results of our exploration apply to other software for differential expression analysis that rely on shared information across replicates and transcripts for the estimation of abundance and dispersion, for example, edgeR [10], BBSeq [56], DSS [57], baySeq [58] and ShrinkBayes [59].

Following the removal of the 3% and 5% of transcripts associated with the highest levels of variation between wolves and dogs, and aggressive and tame foxes, respectively, we observed an increase in the number of differentially expressed transcripts. This may be indicative that for our datasets the removal of variation at these levels optimized the detection of differentially expressed transcripts. Using these 3% and 5% cut-offs, amongst the 50 over expressed genes identified, across the 21 shared gene families, seven genes were shared between dogs and tame foxes (Table 1). Of these seven genes, three main functions related to brain development, neurotransmission, and immune response were identified. These functions have been repeatedly associated with behavior selection during domestication by different approaches, such as QTL analysis [36,60,61], whole-genome sequencing [62–64], and RNA data both using microarrays and RNA-seq [32,37,65–67].

Up until recently, almost no gene overlap had been observed between gene expression profiles involving pairs of domesticated and wild animals [34]. However, a recently published paper performing population genomic and brain transcriptional comparisons in seven bird and mammal domesticated species has revealed a strong convergent pattern in genes implicated in neurotransmission and neuroplasticity [68]. These functions are compatible with those found in our analysis. The shared gene ITGA7 belongs to a gene family that is known to play an essential role in the control of neuronal connectivity [69] and the inflammatory response [70]. Other genes from this family, for example, ITGA8, have been previously observed to be over expressed in tame foxes [66], and here we also observed its over expression in dogs providing further evidence of the family’s role in tameness. Similar functions are associated with the shared genes CHRNA5 [71, 72] and TRIB2 [73] from the cholinergic and tribbles family, respectively. Additionally, we found a shared gene involved in sensing local environmental stimuli, the MYO7A, whose mutation results in loss of hearing and vision [74]. Amongst the three gene families identified as under expressed (Table 2), we found the shared gene STMND1, which deficiency in the amygdala of mice was connected to a deficiency in innate and learned fear [75], a behavior that speculatively could also have an important role in domestication. Although we are aware that this overlap analysis between genes that show the same direction of expression in both dogs and tame foxes is not a formal test for gene convergence, we identified genes involved in several functions previously validated in the scope of domestication.

## Supporting Information Captions

**S1 Figure: Alignment rates**. Mapping success rates (%) resulting from the alignment of the 44 samples used in this study to the complete dog trascnriptome. For each sample, the percentage of aligned reads is presented by the blue bars, while the percentage of reads failing to map is represented in red (the number of raw reads is available in Suplementary Table S1).

**S2 Figure: Distribution of dispersion estimates.** Plots of dispersion estimates in relation to the mean of normalized counts for both case studies, wolves and dogs (left panels), and tame and aggressive foxes (right panels). Estimates were calculated using DESeq2 for the non-filtered (NF) and all filtered datasets (99^th^, 95^th^ and 90^th^ are shown as an example). Gray dots represent the gene-wise maximum likelihood estimates (MLE), the red curve shows the fit to the MLEs, and blue dots identify the final maximum *a posteriori* (MAP) estimates of dispersion. Red dots represent the outliers detected by DESeq2. Both x and y-axis are transformed into a logarithm scale.

**S1 Table**: **Dataset description.** Full details of all datasets, including the location of the relative tissue, age, and sex of each individual, replicate information and sequencing details (FC – frontal cortex; CC – cerebral cortex; PFC – prefrontal cortex; FL – frontal lobe; NS – not specified; F – female; M – male; AD – adult; ya – years old; PE – paired-end; SE – single end).

**S2 Table**: **Removal of intra-condition variation.** Number of transcripts kept and removed from the reference in each case study, wolves and dogs, and aggressive and tame foxes, across the filtered levels used (from the 99^th^ to the 70^th^ percentile). The first ten percentiles were explored in greater detail in steps of one, while the remaining were performed in steps of 5.

**S3 Table**: **Differentially expressed transcripts in wolf *vs.* dog.** Complete list of differentially expressed transcripts in dogs when compared to wolves, identified using non-filtered datasets, and those that got removed within the highest 10% of intra-condition variation, as also those added as differentially expressed across selected filtered datasets (97^th^, 95^th^, and 90^th^ percentiles). The correspondent annotated gene ID, log2FC values and p-values are provided.

**S4 Table**: **Differentially expressed transcripts in aggressive *vs.* tame fox.** Complete list of differentially expressed transcripts in tame foxes when compared to aggressive foxes, identified using non-filtered datasets, and those that got removed within the highest 10% of intra-condition variation, as also those added as differentially expressed across selected filtered datasets (97^th^, 95^th^, and 90^th^ percentiles). The correspondent annotated gene ID, log2FC values and p-values are provided.

**S5 Table**: **Correlation and outliers.** Correlation values (r^2^) and the root mean square error (RMSE) from the regression analysis between the final dispersion estimates and the mean of normalized counts for both case studies, wolves and dogs, and aggressive and tame foxes. The number of outliers identified by DESeq2 are also presented. Values are shown for the non-filtered (NF) and all the filtered datasets used in differential expression analysis.

**S6 Table**: **Shared genes and gene families between non-filtered datasets.** List of the gene families, and shared genes, that were commonly regulated in dogs and tame foxes, using the non-filtered datasets. The number, and name, of the genes within each gene family are provided, with the corresponding log2fold-change values in brackets for each species. Within each family, single genes were charecterized as shared between dogs and tame foxes, or as exclusive to each of the two groups. When more than one transcript for a specific gene was present, all the log2FC values are reported.

## References

1. Wang Z, Gerstein M, Snyder M. RNA-Seq: A revolutionary tool for transcriptomics. Nat Rev Genet. 2009;10: 57–63. doi:10.1038/nrg2484

2. Conesa A, Madrigal P, Tarazona S, Gomez-Cabrero D, Cervera A, McPherson A, et al. A survey of best practices for RNA-seq data analysis. Genome Biol. 2016;17. doi:10.1186/s13059-016-0881-8

3. Kukurba KR, Montgomery SB. RNA Sequencing and Analysis. Cold Spring Harb Protoc. 2015;2015: 951. doi:10.1101/PDB.TOP084970

4. Mortazavi A, Williams BA, McCue K, Schaeffer L, Wold B. Mapping and quantifying mammalian transcriptomes by RNA-Seq. Nat Methods 2008 57. 2008;5: 621–628. doi:10.1038/nmeth.1226

5. Lee S, Li S, Seo CH, Lim B, Yang JO, Oh J, et al. Accurate quantification of transcriptome from RNA-Seq data by effective length normalization. Nucleic Acids Res. 2011;39. doi:10.1093/nar/gkq1015

6. Trapnell C, Williams BA, Pertea G, Mortazavi A, Kwan G, Van Baren MJ, et al. Transcript assembly and quantification by RNA-Seq reveals unannotated transcripts and isoform switching during cell differentiation. Nat Biotechnol. 2010;28: 511–515. doi:10.1038/nbt.1621

7. Jiang H, Wong WH. Statistical inferences for isoform expression in RNA-Seq. Bioinformatics. 2009;25: 1026–1032. doi:10.1093/bioinformatics/btp113

8. Nagalakshmi U, Wang Z, Waern K, Shou C, Raha D, Gerstein M, et al. The transcriptional landscape of the yeast genome defined by RNA sequencing. Science (80- ). 2008;320: 1344–1349. doi:10.1126/science.1158441

9. Love MI, Huber W, Anders S. Moderated estimation of fold change and dispersion for RNA-seq data with DESeq2. Genome Biol. 2014;15:550. doi:10.1186/s13059-014-0550-8

10. Robinson MD, McCarthy DJ, Smyth GK. edgeR: a Bioconductor package for differential expression analysis of digital gene expression data. Bioinformatics. 2010;26: 139–140. doi:10.1093/bioinformatics/btp616

11. Hansen KD, Wu Z, Irizarry RA, Leek JT. Sequencing technology does not eliminate biological variability. Nat Biotechnol. 2011;29: 572–573. doi:10.1038/nbt.1910

12. Xu Z, Asakawa S. Physiological RNA dynamics in RNA-Seq analysis. Brief Bioinform. 2019;20: 1725–1733. doi:10.1093/bib/bby045

13. McIntyre LM, Lopiano KK, Morse AM, Amin V, Oberg AL, Young LJ, et al. RNA-seq: Technical variability and sampling. BMC Genomics. 2011;12:293. doi:10.1186/1471-2164-12-293

14. Liu Y, Zhou J, White KP. RNA-seq differential expression studies: More sequence or more replication? Bioinformatics. 2014;30: 301–304. doi:10.1093/bioinformatics/btt688

15. Schurch NJ, Schofield P, Gierliński M, Cole C, Sherstnev A, Singh V, et al. How many biological replicates are needed in an RNA-seq experiment and which differential expression tool should you use? RNA. 2016;22: 839–851. doi:10.1261/rna.053959.115

16. Robles JA, Qureshi SE, Stephen SJ, Wilson SR, Burden CJ, Taylor JM. Efficient experimental design and analysis strategies for the detection of differential expression using RNA-Sequencing. BMC Genomics. 2012;13: 484. doi:10.1186/1471-2164-13-484

17. Lachmann A, Torre D, Keenan AB, Jagodnik KM, Lee HJ, Wang L, et al. Massive mining of publicly available RNA-seq data from human and mouse. Nat Commun 2018 91. 2018;9: 1–10. doi:10.1038/s41467-018-03751-6

18. Zoabi Y, Shomron N. Processing and Analysis of RNA-seq Data from Public Resources. Methods in Molecular Biology. Methods Mol Biol; 2021. pp. 81–94. doi:10.1007/978-1-0716-1103-6_4

19. Sudmant PH, Alexis MS, Burge CB. Meta-analysis of RNA-seq expression data across species, tissues and studies. Genome Biol 2015 161. 2015;16: 1–11. doi:10.1186/S13059-015-0853-4

20. Rau A, Marot G, Jaffrézic F. Differential meta-analysis of RNA-seq data from multiple studies. BMC Bioinformatics. 2014;15:91. doi:10.1186/1471-2105-15-91

21. Jeng SL, Chi YC, Ma MC, Chan SH, Sun HS. Gene expression analysis of combined RNA-seq experiments using a receiver operating characteristic calibrated procedure. Comput Biol Chem. 2021;93: 107515. doi:10.1016/J.COMPBIOLCHEM.2021.107515

22. McDermaid A, Monier B, Zhao J, Liu B, Ma Q. Interpretation of differential gene expression results of RNA-seq data: review and integration. Brief Bioinform. 2019;20: 2044. doi:10.1093/BIB/BBY067

23. Quinn TP, Crowley TM, Richardson MF. Benchmarking differential expression analysis tools for RNA-Seq: normalization-based vs. log-ratio transformation-based methods. BMC Bioinformatics. 2018;19: 274. doi:10.1186/s12859-018-2261-8

24. Risso D, Schwartz K, Sherlock G, Dudoit S. GC-Content Normalization for RNA-Seq Data. BMC Bioinformatics. 2011. doi:10.1186/1471-2105-12-480

25. Stegle O, Parts L, Durbin R, Winn J. A bayesian framework to account for complex non-genetic factors in gene expression levels greatly increases power in eQTL studies. PLoS Comput Biol. 2010;6(5): e100. doi:10.1371/journal.pcbi.1000770

26. Li S, Labaj PP, Zumbo P, Sykacek P, Shi W, Shi L, et al. Detecting and correcting systematic variation in large-scale RNA sequencing data. Nat Biotechnol. 2014;32: 888–895. doi:10.1038/nbt.3000

27. Wang T, Li B, Nelson CE, Nabavi S. Comparative analysis of differential gene expression analysis tools for single-cell RNA sequencing data. BMC Bioinformatics. 2019;20: 40. doi:10.1186/s12859-019-2599-6

28. Mou T, Deng W, Gu F, Pawitan Y, Vu TN. Reproducibility of Methods to Detect Differentially Expressed Genes from Single-Cell RNA Sequencing. Front Genet. 2020;10. doi:10.3389/fgene.2019.01331

29. Arora S, Pattwell SS, Holland EC, Bolouri H. Variability in estimated gene expression among commonly used RNA-seq pipelines. Sci Rep. 2020;10: 2734. doi:10.1038/s41598-020-59516-z

30. Stupnikov A, McInerney CE, Savage KI, McIntosh SA, Emmert-Streib F, Kennedy R, et al. Robustness of differential gene expression analysis of RNA-seq. Comput Struct Biotechnol J. 2021;19: 3470–3481. doi:10.1016/j.csbj.2021.05.040

31. Costa-Silva J, Domingues D, Lopes FM. RNA-Seq differential expression analysis: An extended review and a software tool. PLoS One. 2017;12:e019015. doi:10.1371/journal.pone.0190152

32. Li Y, Wang GD, Wang MS, Irwin DM, Wu DD, Zhang YP. Domestication of the dog from the Wolf was promoted by enhanced excitatory synaptic plasticity: A hypothesis. Genome Biol Evol. 2014;6: 3115–3121. doi:10.1093/gbe/evu245

33. Li Y, VonHoldt BM, Reynolds A, Boyko AR, Wayne RK, Wu DD, et al. Artificial selection on brain-expressed genes during the domestication of dog. Mol Biol Evol. 2013;30: 1867–1876. doi:10.1093/molbev/mst088

34. Albert FW, Somel M, Carneiro M, Aximu-Petri A, Halbwax M, Thalmann O, et al. A Comparison of Brain Gene Expression Levels in Domesticated and Wild Animals. Akey JM, editor. PLoS Genet. 2012;8:e1002962. doi:10.1371/journal.pgen.1002962

35. Lord KA, Larson G, Coppinger RP, Karlsson EK. The History of Farm Foxes Undermines the Animal Domestication Syndrome. Trends Ecol Evol. 2019. doi:10.1016/j.tree.2019.10.011

36. Kukekova A, Trut L, Chase K, Kharlamova A, Johnson J, Temnykh S, et al. Mapping loci for fox domestication: Deconstruction/Reconstruction of a behavioral phenotype. Behav Genet. 2011;41: 593–606. doi:10.1007/s10519-010-9418-1

37. Wang X, Pipes L, Trut L, Herbeck Y, Vladimirova A, Gulevich R, et al. Genomic responses to selection for tame/aggressive behaviors in the silver fox (Vulpes vulpes). Proc Natl Acad Sci. 2018;115: 10398–10403. doi:10.1073/pnas.1800889115

38. Hekman J, Johnson J, Edwards W, Vladimirova A, Gulevich R, Ford A, et al. Anterior Pituitary Transcriptome Suggests Differences in ACTH Release in Tame and Aggressive Foxes. G3; Genes|Genomes|Genetics. 2018;8: 859–873. doi:10.1534/g3.117.300508

39. Roy M, Kim N, Kim K, Chung WH, Achawanantakun R, Sun Y, et al. Analysis of the canine brain transcriptome with an emphasis on the hypothalamus and cerebral cortex. Mamm Genome. 2013;24: 484–499. doi:10.1007/s00335-013-9480-0

40. Fushan AA, Turanov AA, Lee SG, Kim EB, Lobanov A V, Yim SH, et al. Gene expression defines natural changes in mammalian lifespan. Aging Cell. 2015;14: 352–365. doi:10.1111/acel.12283

41. Hoeppner MP, Lundquist A, Pirun M, Meadows JRS, Zamani N, Johnson J, et al. An improved canine genome and a comprehensive catalogue of coding genes and non-coding transcripts. PLoS One. 2014;9(3):91172. doi:10.1371/journal.pone.0091172

42. Hou Y, Qi F, Bai X, Ren T, Shen X, Chu Q, et al. Genome-wide analysis reveals molecular convergence underlying domestication in 7 bird and mammals. BMC Genomics. 2020;21: 1–20. doi:10.1186/s12864-020-6613-1

43. Langmead B, Salzberg SL. Fast gapped-read alignment with Bowtie 2. Nat Methods. 2012. doi:10.1038/nmeth.1923

44. Bushnell, Brian. BBMap: A Fast, Accurate, Splice-Aware Aligner. Conference: 9th Annual Genomics of Energy {\&} Environment Meeting. 2014. doi:10.1186/1471-2105-13-238

45. R Development Core Team. R: A language and environment for statistical computing. Vienna, Austria. 2017. doi:R Foundation for Statistical Computing, Vienna, Austria. ISBN 3-900051-07-0, URL http://www.R-project.org.

46. Durinck S, Moreau Y, Kasprzyk A, Davis S, De Moor B, Brazma A, et al. BioMart and Bioconductor: A powerful link between biological databases and microarray data analysis. Bioinformatics. 2005;21: 3439–3440. doi:10.1093/bioinformatics/bti525

47. Perri AR, Feuerborn TR, Frantz LAF, Larson G, Malhi RS, Meltzer DJ, et al. Dog domestication and the dual dispersal of people and dogs into the Americas. Proc Natl Acad Sci U S A. 2021;118: 1–8. doi:10.1073/pnas.2010083118

48. Tu J, Guo J, Li J, Gao S, Yao B, Lu Z. Systematic Characteristic Exploration of the Chimeras Generated in Multiple Displacement Amplification through Next Generation Sequencing Data Reanalysis. PLoS One. 2015;6: e0139857. doi:10.1371/journal.pone.0139857

49. Rizzi R, Beretta S, Patterson M, Pirola Y, Previtali M, Della Vedova G, et al. Overlap graphs and de Bruijn graphs: data structures for de novo genome assembly in the big data era. Quant Biol. 2019;7: 278–292. doi:10.1007/s40484-019-0181-x

50. Wayne RK, Geffen E, Girman DJ, Koepfli K-P, Lau LM, Marshall CR. Molecular Systematics of the Canidae. Syst Biol. 1997;46: 622–653. doi:doi:10.1093/sysbio/46.4.622

51. Hsieh PH, Oyang YJ, Chen CY. Effect of de novo transcriptome assembly on transcript quantification. Sci Rep. 2019;9. doi:10.1038/s41598-019-44499-3

52. Reinert K, Langmead B, Weese D, Evers DJ. Alignment of Next-Generation Sequencing Reads. Annu Rev Genomics Hum Genet. 2015;16: 133–151. doi:10.1146/annurev-genom-090413-025358

53. Evans C, Hardin J, Stoebel DM. Selecting between-sample RNA-Seq normalization methods from the perspective of their assumptions. Brief Bioinform. 2018;19: 776–792. doi:10.1093/bib/bbx008

54. Brodin J, Mild M, Hedskog C, Sherwood E, Leitner T, Andersson B, et al. PCR-Induced Transitions Are the Major Source of Error in Cleaned Ultra-Deep Pyrosequencing Data. PLoS One. 2013;8. doi:10.1371/journal.pone.0070388

55. Ma X, Shao Y, Tian L, Flasch DA, Mulder HL, Edmonson MN, et al. Analysis of error profiles in deep next-generation sequencing data. Genome Biol. 2019;20. doi:10.1186/s13059-019-1659-6

56. Zhou YH, Xia K, Wright FA. A powerful and flexible approach to the analysis of RNA sequence count data. Bioinformatics. 2011. doi:10.1093/bioinformatics/btr449

57. Wu H, Wang C, Wu Z. A new shrinkage estimator for dispersion improves differential expression detection in RNA-seq data. Biostatistics. 2013;14: 232–243. doi:10.1093/biostatistics/kxs033

58. Hardcastle TJ, Kelly KA. BaySeq: Empirical Bayesian methods for identifying differential expression in sequence count data. BMC Bioinformatics. 2010;11:422. doi:10.1186/1471-2105-11-422

59. Van De Wiel MA, Leday GGR, Pardo L, Rue H, Van Der Vaart AW, Van Wieringen WN. Bayesian analysis of RNA sequencing data by estimating multiple shrinkage priors. Biostatistics. 2013;14: 113–128. doi:10.1093/biostatistics/kxs031

60. Wirén A, Wright D, Jensen P. Domestication-related variation in social preferences in chickens is affected by genotype on a growth QTL. Genes, Brain Behav. 2013;12: 330–337. doi:10.1111/gbb.12017

61. Albert FW, Carlborg Ö, Plyusnina I, Besnier F, Hedwig D, Lautenschläger S, et al. Genetic architecture of tameness in a rat model of animal domestication. Genetics. 2009;182: 541–554. doi:10.1534/genetics.109.102186

62. Carneiro M, Rubin CJ, Palma F Di, Albert FW, Alföldi J, Barrio AM, et al. Rabbit genome analysis reveals a polygenic basis for phenotypic change during domestication. Science (80- ). 2014;345: 1074–1079. doi:10.1126/science.1253714

63. Freedman AH, Schweizer RM, Ortega-Del Vecchyo D, Han E, Davis BW, Gronau I, et al. Demographically-Based Evaluation of Genomic Regions under Selection in Domestic Dogs. PLoS Genet. 2016;12:e100585. doi:10.1371/journal.pgen.1005851

64. Kukekova A, Johnson J, Xiang X-Y, Feng S-H, Liu S, Rando H, et al. The red fox genome assembly identifies genomic regions associated with tame and aggressive behaviors. Nat Ecol Evol. 2018;2: 1479–1491. doi:10.1038/s41559-018-0611-6

65. Saetre P, Lindberg J, Ellegren H, Vila C, Jazin E, Leonard JA, et al. From wild wolf to domestic dog: Gene expression changes in the brain. Mol Brain Res. 2004;126: 198–206. doi:10.1016/j.molbrainres.2004.05.003

66. Kukekova A, Johnson J, Teiling C, Li L, Oskina I, Kharlamova A, et al. Sequence comparison of prefrontal cortical brain transcriptome from a tame and an aggressive silver fox (Vulpes vulpes). BMC Genomics. 2011;12:482. doi:10.1186/1471-2164-12-482

67. Heyne HO, Lautenschläger S, Nelson R, Besnier F, Rotival M, Cagan A, et al. Genetic influences on brain gene expression in rats selected for tameness and aggression. Genetics. 2014;198: 1277–1290. doi:10.1534/genetics.114.168948

68. Hou Y, Qi F, Bai X, Ren T, Shen X, Chu Q, et al. Genome-wide analysis reveals molecular convergence underlying domestication in 7 bird and mammals. BMC Genomics. 2020;21. doi:10.1186/s12864-020-6613-1

69. Lilja J, Ivaska J. Integrin activity in neuronal connectivity. J Cell Sci. 2018;131:jcs212. doi:10.1242/jcs.212803

70. González-Amaro R, Sánchez-Madrid F. Cell adhesion molecules: selectins and integrins. Crit Rev Immunol. 1999;19: 389–429.

71. Winterer G, Mittelstrass K, Giegling I, Lamina C, Fehr C, Brenner H, et al. Risk gene variants for nicotine dependence in the CHRNA5-CHRNA3-CHRNB4 cluster are associated with cognitive performance. Am J Med Genet Part B Neuropsychiatr Genet. 2010;153: 1448–1458. doi:10.1002/ajmg.b.31126

72. Zhang H, Kranzler HR, Poling J, Gruen JR, Gelernter J. Cognitive flexibility is associated with KIBRA variant and modulated by recent tobacco use. Neuropsychopharmacology. 2009;34: 2508– 2516. doi:10.1038/npp.2009.80

73. Eyers PA, Keeshan K, Kannan N. Tribbles in the 21st Century: The Evolving Roles of Tribbles Pseudokinases in Biology and Disease. Trends Cell Biol. 2017;27: 284–298. doi:10.1016/j.tcb.2016.11.002

74. Miller KA, Williams LH, Rose E, Kuiper M, Dahl HHM, Manji SSM. Inner Ear Morphology Is Perturbed in Two Novel Mouse Models of Recessive Deafness. PLoS One. 2012;7(12):e512. doi:10.1371/journal.pone.0051284

75. Martel G, Nishi A, Shumyatsky GP. Stathmin reveals dissociable roles of the basolateral amygdala in parental and social behaviors. Proc Natl Acad Sci U S A. 2008;105: 14620–14625. doi:10.1073/pnas.0807507105

