## Supplementary figures and images for "On taming the effect of transcript level intra-condition count variation during differential expression analysis: a story of dogs, foxes and wolves"

### Supplemental Figure 1

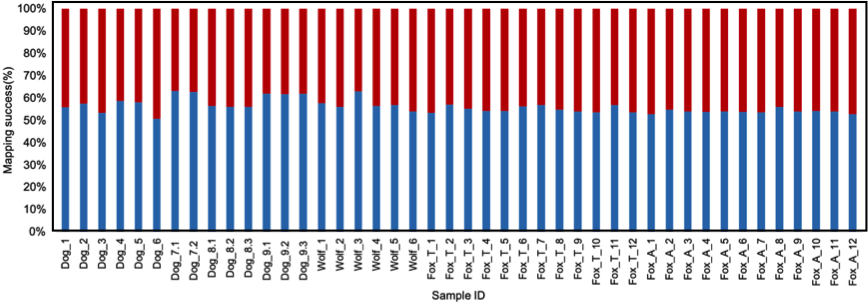

### Supplemental Figure 2

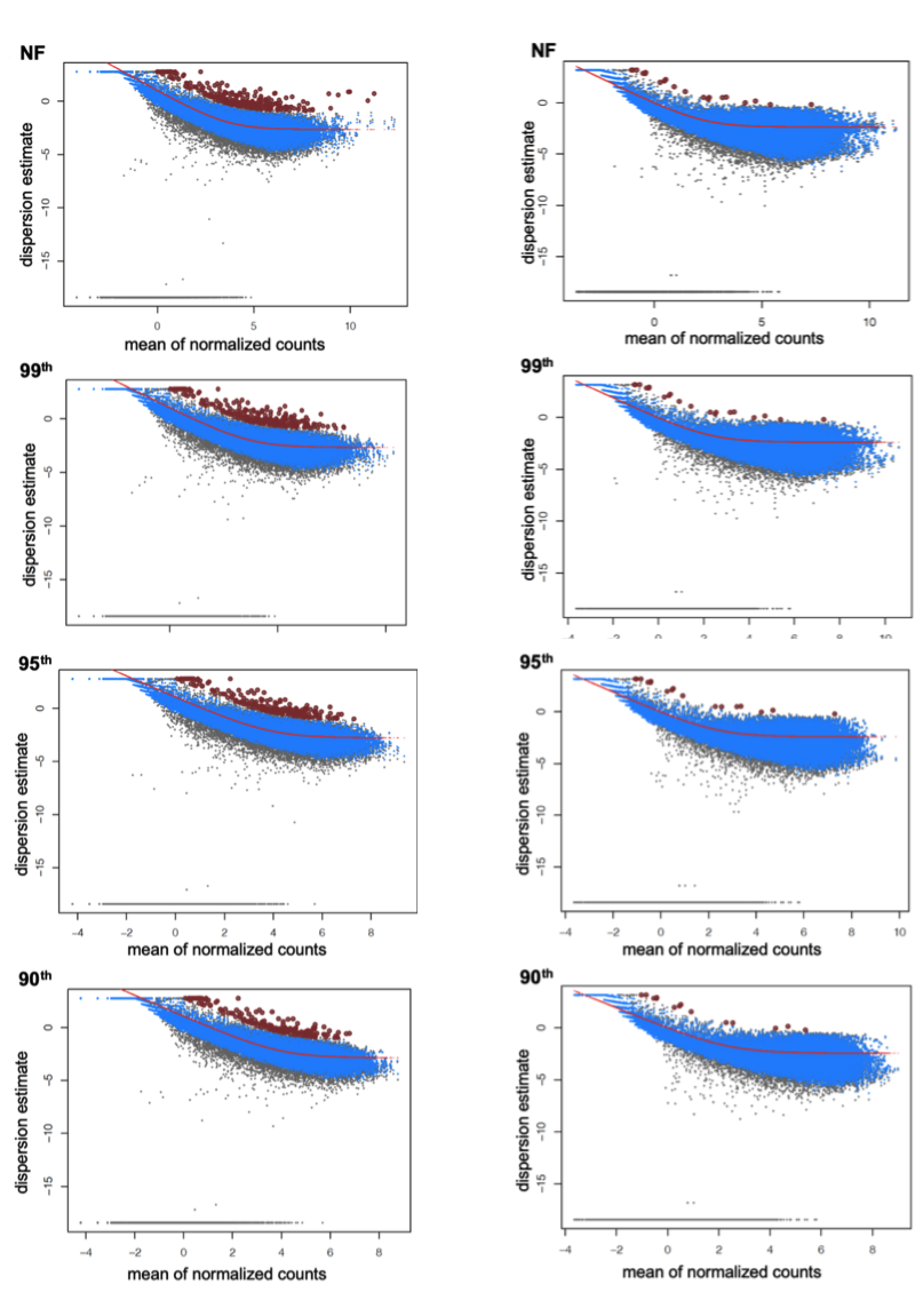
