## Supplemental Table 1 for "On taming the effect of transcript level intra-condition count variation during differential expression analysis: a story of dogs, foxes and wolves"

| **Species** | **Study** | **Sample ID** | **No. individuals** | **Tissue** | **Compartment** | **Sex** | **Age** | **Condition** | **Library** | **Sequencing platform** | **Insert size (bp)** | **N of raw reads** | **Project accession** | **Run accession** |
| --- | --- | --- | --- | --- | --- | --- | --- | --- | --- | --- | --- | --- | --- | --- |
| **Canis *familiaris*** | Albert *et* *al* 2012 | Dog_1 | single | brain | FC | F | old AD | domesticated | PE | Illumina GA II | 51 | 43 363 388 | PRJEB3197 | ERR266355 |
|  |  | Dog_2 |  |  |  | M |  |  |  |  |  | 34 735 876 |  | ERR266386 |
|  |  | Dog_3 |  |  |  | M |  |  |  |  |  | 31 956 420 |  | ERR266395 |
|  |  | Dog_4 |  |  |  | F |  |  |  |  |  | 41 264 506 |  | ERR266403 |
|  |  | Dog_5 |  |  |  | M |  |  |  |  |  | 42 044 332 |  | ERR266382 |
|  | Roy *et* *al* 2013 | Dog_6 | several | brain | CC | M | 1-2 ya | domesticated | PE | Illumina HiSeq 2000 | 100 | 178 091 798 | PRJEB4668 | ERR351173 |
|  | Fushan *et a*l 2015 | Dog_7.1 | single | brain | PFC / FL | M | young AD | domesticated | PE | Illumina HiSeq 2000 | 51 | 29 915 644 | PRJNA185055 | SRR636937 |
|  |  | Dog_7.2 |  |  |  |  |  |  |  |  |  | 28 668 648 |  | SRR636938 |
|  | Hoeppner *et al* 2014 | Dog_8.1 | several | brain | NS | NS | NS | domesticated | PE | Illumina HiSeq 2000 | 580 | 18 561 120 | PRJNA78827 | SRR388737 |
|  |  | Dog_8.2 |  |  |  |  |  |  |  |  |  | 18 534 772 |  | SRR388740 |
|  |  | Dog_8.3 |  |  |  |  |  |  |  |  |  | 18 488 968 |  | SRR388766 |
|  |  | Dog_9.1 | several | brain |  |  |  |  |  |  | 450 | 40 195 676 |  | SRR543733 |
|  |  | Dog_9.2 |  |  |  |  |  |  |  |  |  | 41 554 898 |  | SRR536881 |
|  |  | Dog_9.3 |  |  |  |  |  |  |  |  |  | 39 271 576 |  | SRR536883 |
| **Canis *lupus*** | Albert *et al* 2012 | Wolf_1 | single | brain | FC | M | old AD | wild | PE | Illumina GA II | 51 | 42 800 148 | PRJEB3197 | ERR266407 |
|  |  | Wolf_2 |  |  |  | F |  |  |  |  |  | 43 721 960 |  | ERR266371 |
|  |  | Wolf_3 |  |  |  | F |  |  |  |  |  | 37 195 758 |  | ERR266359 |
|  |  | Wolf_4 |  |  |  | M |  |  |  |  |  | 40 138 236 |  | ERR266374 |
|  |  | Wolf_5 |  |  |  | M |  |  |  |  |  | 32 500 494 |  | ERR266366 |
|  |  | Wolf_6 |  |  |  | M |  |  |  |  |  | 43 088 096 |  | ERR266400 |
| **Vulpes *vulpes*** | Wang *et* *al* 2018 | Fox_T_1 | single | brain | PFC | M | 1.5 ya | tame | SE | Illumina HiSeq 2000 | 51 | 30 760 396 | PRJNA307604 | SRR3084300 |
|  |  | Fox_T_2 |  |  |  |  |  |  |  |  |  | 23 943 747 |  | SRR3084299 |
|  |  | Fox_T_3 |  |  |  |  |  |  |  |  |  | 38 957 530 |  | SRR3084298 |
|  |  | Fox_T_4 |  |  |  |  |  |  |  |  |  | 43 980 944 |  | SRR3084297 |
|  |  | Fox_T_5 |  |  |  |  |  |  |  |  |  | 50 603 352 |  | SRR3084296 |
|  |  | Fox_T_6 |  |  |  |  |  |  |  |  |  | 35 614 595 |  | SRR3084295 |
|  |  | Fox_T_7 |  |  |  |  |  |  |  |  |  | 23 317 339 |  | SRR3084294 |
|  |  | Fox_T_8 |  |  |  |  |  |  |  |  |  | 37 020 077 |  | SRR3084293 |
|  |  | Fox_T_9 |  |  |  |  |  |  |  |  |  | 20 659 559 |  | SRR3084292 |
|  |  | Fox_T_10 |  |  |  |  |  |  |  |  |  | 31 077 357 |  | SRR3084291 |
|  |  | Fox_T_11 |  |  |  |  |  |  |  |  |  | 42 038 051 |  | SRR3084290 |
|  |  | Fox_T_12 |  |  |  |  |  |  |  |  |  | 25 492 942 |  | SRR3084289 |
|  |  | Fox_A_1 |  |  |  |  |  | aggressive |  |  |  | 33 541 449 |  | SRR3084312 |
|  |  | Fox_A_2 |  |  |  |  |  |  |  |  |  | 40 582 881 |  | SRR3084311 |
|  |  | Fox_A_3 |  |  |  |  |  |  |  |  |  | 43 558 186 |  | SRR3084310 |
|  |  | Fox_A_4 |  |  |  |  |  |  |  |  |  | 32 082 055 |  | SRR3084309 |
|  |  | Fox_A_5 |  |  |  |  |  |  |  |  |  | 47 274 581 |  | SRR3084308 |
|  |  | Fox_A_6 |  |  |  |  |  |  |  |  |  | 28 570 574 |  | SRR3084307 |
|  |  | Fox_A_7 |  |  |  |  |  |  |  |  |  | 35 859 502 |  | SRR3084306 |
|  |  | Fox_A_8 |  |  |  |  |  |  |  |  |  | 25 896 719 |  | SRR3084305 |
|  |  | Fox_A_9 |  |  |  |  |  |  |  |  |  | 45 247 750 |  | SRR3084304 |
|  |  | Fox_A_10 |  |  |  |  |  |  |  |  |  | 45 763 357 |  | SRR3084303 |
|  |  | Fox_A_11 |  |  |  |  |  |  |  |  |  | 27 253 195 |  | SRR3084302 |
|  |  | Fox_A_12 |  |  |  |  |  |  |  |  |  | 18 588 067 |  | SRR3084301 |
