## Supplemental Table 2 for "On taming the effect of transcript level intra-condition count variation during differential expression analysis: a story of dogs, foxes and wolves"

|  | **Wolves and dogs** | | **Aggressive and tame foxes** | |
| --- | --- | --- | --- | --- |
| Percentile | Transcripts kept | Transcripts removed | Transcripts kept | Transcripts removed |
| 99 | 25923 | 184 | 25872 | 235 |
| 98 | 25601 | 506 | 25428 | 679 |
| 97 | 25253 | 854 | 25018 | 1089 |
| 96 | 24916 | 1191 | 24585 | 1522 |
| 95 | 24574 | 1533 | 24167 | 1940 |
| 94 | 24237 | 1870 | 23767 | 2340 |
| 93 | 23921 | 2186 | 23372 | 2735 |
| 92 | 23611 | 2496 | 22978 | 3129 |
| 91 | 23293 | 2814 | 22578 | 3529 |
| 90 | 22973 | 3134 | 22219 | 3888 |
| 85 | 21399 | 4708 | 20407 | 5700 |
| 80 | 19867 | 6240 | 18723 | 7384 |
| 75 | 18361 | 7746 | 17128 | 8979 |
| 70 | 16958 | 9149 | 15661 | 10446 |
