## Supplemental Table 3 for "On taming the effect of transcript level intra-condition count variation during differential expression analysis: a story of dogs, foxes and wolves"

| **Ranking** | **Transcript ID** | **Gene ID** | **log2 FC** | **p-adj value** | **Filter level** |
| --- | --- | --- | --- | --- | --- |
| 1 | ENSCAFT00000049574 | BCO1 | 3,68 | 1,51E-16 | Not filtered |
| 2 | ENSCAFT00000030934 | TKTL1 | 5,12 | 4,77E-10 | Not filtered |
| 3 | ENSCAFT00000049123 | CCDC190 | 2,23 | 4,90E-10 | Not filtered |
| 4 | ENSCAFT00000029783 | PIEZO2 | 2,12 | 7,21E-10 | Not filtered |
| 5 | ENSCAFT00000010686 | EFCAB1 | 2,60 | 2,47E-09 | Not filtered |
| 6 | ENSCAFT00000047954 | SCN11A | 1,61 | 3,72E-08 | Not filtered |
| 7 | ENSCAFT00000044099 | PIEZO2 | 1,90 | 1,58E-07 | Not filtered |
| 8 | ENSCAFT00000043213 | SCN11A | 1,46 | 4,19E-07 | Not filtered |
| 9 | ENSCAFT00000022535 | ERBB4 | 1,36 | 6,12E-07 | Not filtered |
| 10 | ENSCAFT00000038874 |  | -3,39 | 1,17E-06 | Not filtered |
| 11 | ENSCAFT00000044516 |  | -2,98 | 1,74E-06 | Not filtered |
| 12 | ENSCAFT00000048021 |  | -0,96 | 2,03E-06 | Not filtered |
| **13** | **ENSCAFT00000036702** | **CRYZ** | **1,49** | **6,14E-06** | **95^th^** |
| **14** | **ENSCAFT00000035561** |  | **-8,10** | **6,14E-06** | **98^th^** |
| 15 | ENSCAFT00000036942 |  | -3,76 | 8,95E-06 | Not filtered |
| 16 | ENSCAFT00000048139 | LYPD6 | 1,36 | 1,17E-05 | Not filtered |
| 17 | ENSCAFT00000001452 | COL14A1 | 1,79 | 1,19E-05 | Not filtered |
| 18 | ENSCAFT00000025553 | AP5Z1 | -1,17 | 1,51E-05 | Not filtered |
| 19 | ENSCAFT00000005798 | BVES | 1,28 | 2,34E-05 | Not filtered |
| 20 | ENSCAFT00000027675 | NSMCE1 | -0,62 | 2,34E-05 | Not filtered |
| 21 | ENSCAFT00000012020 | TDRD12 | 1,38 | 2,95E-05 | Not filtered |
| 22 | ENSCAFT00000021631 |  | -3,27 | 3,40E-05 | Not filtered |
| **23** | **ENSCAFT00000013788** | **PTK2B** | **-0,78** | **4,58E-05** | **97^th^** |
| 24 | ENSCAFT00000019357 | CTSZ | -1,29 | 4,62E-05 | Not filtered |
| 25 | ENSCAFT00000000233 | CCDC68 | 1,98 | 5,43E-05 | Not filtered |
| 26 | ENSCAFT00000010803 | NLRP14 | 1,68 | 8,97E-05 | Not filtered |
| **27** | **ENSCAFT00000001241** |  | **-0,98** | **1,04E-04** | **94^th^** |
| 28 | ENSCAFT00000048789 | ACCS | 0,69 | 1,05E-04 | Not filtered |
| **29** | **ENSCAFT00000028679** | **TTR** | **5,73** | **1,16E-04** | **99^th^** |
| 30 | ENSCAFT00000010637 | STK32A | 1,11 | 1,39E-04 | Not filtered |
| 31 | ENSCAFT00000049816 | PODXL | 0,96 | 1,39E-04 | Not filtered |
| 32 | ENSCAFT00000048591 | HAPLN1 | 1,16 | 1,57E-04 | Not filtered |
| 33 | ENSCAFT00000022447 |  | -2,96 | 1,77E-04 | Not filtered |
| 34 | ENSCAFT00000012985 | PLSCR4 | 1,27 | 2,00E-04 | Not filtered |
| 35 | ENSCAFT00000032487 | SLC44A5 | 1,45 | 2,33E-04 | Not filtered |
| 36 | ENSCAFT00000045631 |  | -2,92 | 2,33E-04 | Not filtered |
| 37 | ENSCAFT00000007509 | ITGA8 | 1,14 | 2,49E-04 | Not filtered |
| 38 | ENSCAFT00000015081 | SEMA3G | 1,18 | 2,56E-04 | Not filtered |
| 39 | ENSCAFT00000012064 | ADM | -1,50 | 3,15E-04 | Not filtered |
| 40 | ENSCAFT00000004810 | NT5E | 1,45 | 3,45E-04 | Not filtered |
| 41 | ENSCAFT00000017267 | SYNM | -0,41 | 3,68E-04 | Not filtered |
| 42 | ENSCAFT00000043196 |  | -0,83 | 3,68E-04 | Not filtered |
| 43 | ENSCAFT00000004066 | PLPPR1 | 1,01 | 3,79E-04 | Not filtered |
| 44 | ENSCAFT00000006882 | MYO3A | 1,12 | 3,81E-04 | Not filtered |
| 45 | ENSCAFT00000005420 | FIGNL1 | 0,92 | 4,24E-04 | Not filtered |
| 46 | ENSCAFT00000022445 |  | -4,61 | 4,69E-04 | Not filtered |
| 47 | ENSCAFT00000013736 | SOX6 | 1,27 | 5,12E-04 | Not filtered |
| 48 | ENSCAFT00000001407 | COL11A2 | 1,23 | 5,12E-04 | Not filtered |
| 49 | ENSCAFT00000031837 | VCAM1 | 1,37 | 6,01E-04 | Not filtered |
| 50 | ENSCAFT00000039500 | POSTN | 1,83 | 6,36E-04 | Not filtered |
| 51 | ENSCAFT00000026335 | MRI1 | -1,33 | 7,30E-04 | Not filtered |
| 52 | ENSCAFT00000022067 | SERPINE1 | -2,03 | 7,84E-04 | Not filtered |
| 53 | ENSCAFT00000027824 | MYH3 | 1,92 | 9,48E-04 | Not filtered |
| 54 | ENSCAFT00000047995 | PITPNC1 | 1,15 | 9,79E-04 | Not filtered |
| 55 | ENSCAFT00000045799 | TDRD12 | 1,12 | 9,79E-04 | Not filtered |
| 56 | ENSCAFT00000020699 | ITGA6 | 1,25 | 1,02E-03 | Not filtered |
| **57** | **ENSCAFT00000015150** | **SPARCL1** | **0,81** | **1,08E-03** | **99^th^** |
| **58** | **ENSCAFT00000039007** | **CLU** | **1,00** | **1,21E-03** | **99^th^** |
| 59 | ENSCAFT00000046959 |  | 4,49 | 1,33E-03 | Not filtered |
| 60 | ENSCAFT00000044997 | SLC25A48 | 1,29 | 1,46E-03 | Not filtered |
| 61 | ENSCAFT00000022158 | SYTL5 | 1,11 | 1,46E-03 | Not filtered |
| 62 | ENSCAFT00000006393 | SPATA5 | 0,88 | 1,46E-03 | Not filtered |
| 63 | ENSCAFT00000026672 |  | -1,43 | 1,46E-03 | Not filtered |
| **64** | **ENSCAFT00000011873** |  | **-1,66** | **1,46E-03** | **92^nd^** |
| 65 | ENSCAFT00000021042 |  | -3,49 | 1,46E-03 | Not filtered |
| **66** | **ENSCAFT00000038737** | **SPARCL1** | **0,94** | **1,48E-03** | **99^th^** |
| **67** | **ENSCAFT00000010134** | **CHL1** | **0,99** | **1,56E-03** | **91^st^** |
| 68 | ENSCAFT00000035851 |  | -0,98 | 1,64E-03 | Not filtered |
| 69 | ENSCAFT00000026736 | ITGAX | 0,97 | 1,76E-03 | Not filtered |
| 70 | ENSCAFT00000022797 | VSIR | 1,29 | 1,85E-03 | Not filtered |
| **71** | **ENSCAFT00000013347** | **CLU** | **0,86** | **1,85E-03** | **99^th^** |
| **72** | **ENSCAFT00000049861** | **HSPB8** | **-0,69** | **1,87E-03** | **90^th^** |
| 73 | ENSCAFT00000014739 | CRHBP | 1,13 | 2,23E-03 | Not filtered |
| **74** | **ENSCAFT00000047619** | **CHL1** | **1,02** | **2,28E-03** | **92^nd^** |
| 75 | ENSCAFT00000003015 | ATP10D | 0,63 | 2,52E-03 | Not filtered |
| 76 | ENSCAFT00000003270 | PDGFRA | 1,46 | 2,59E-03 | Not filtered |
| 77 | ENSCAFT00000011166 | RGS20 | 0,98 | 2,61E-03 | Not filtered |
| 78 | ENSCAFT00000010726 | SUCLG2 | 0,84 | 2,80E-03 | Not filtered |
| 79 | ENSCAFT00000021625 |  | -5,08 | 2,85E-03 | Not filtered |
| 80 | ENSCAFT00000010945 | FZD10 | 1,82 | 2,86E-03 | Not filtered |
| 81 | ENSCAFT00000043504 |  | -1,37 | 2,89E-03 | Not filtered |
| 82 | ENSCAFT00000013975 | GRIK1 | 0,87 | 2,97E-03 | Not filtered |
| 83 | ENSCAFT00000046707 |  | -1,10 | 3,19E-03 | Not filtered |
| **84** | **ENSCAFT00000044854** | **CHL1** | **1,01** | **3,21E-03** | **92^nd^** |
| 85 | ENSCAFT00000037196 | TLR1 | 1,23 | 3,23E-03 | Not filtered |
| 86 | ENSCAFT00000015510 | ACAD8 | -0,68 | 3,33E-03 | Not filtered |
| 87 | ENSCAFT00000049563 | THEM4 | 1,10 | 3,34E-03 | Not filtered |
| **88** | **ENSCAFT00000037353** | **SMIM3** | **-1,36** | **3,34E-03** | **92^nd^** |
| 89 | ENSCAFT00000002896 | ABCB1 | 0,93 | 3,55E-03 | Not filtered |
| 90 | ENSCAFT00000029252 | RHBDL3 | -1,33 | 3,55E-03 | Not filtered |
| 91 | ENSCAFT00000019970 | CD84 | 1,56 | 3,73E-03 | Not filtered |
| 92 | ENSCAFT00000002169 | PODXL | 0,85 | 3,73E-03 | Not filtered |
| 93 | ENSCAFT00000025758 | FUT8 | 0,71 | 3,73E-03 | Not filtered |
| 94 | ENSCAFT00000014585 | ARSB | 0,59 | 3,73E-03 | Not filtered |
| 95 | ENSCAFT00000000458 | ADGRG6 | 1,46 | 3,82E-03 | Not filtered |
| 96 | ENSCAFT00000043004 | GBA2 | -0,53 | 3,85E-03 | Not filtered |
| 97 | ENSCAFT00000049725 |  | 1,23 | 3,90E-03 | Not filtered |
| 98 | ENSCAFT00000028525 | DTNA | 0,53 | 3,94E-03 | Not filtered |
| 99 | ENSCAFT00000035852 |  | -1,00 | 4,27E-03 | Not filtered |
| 100 | ENSCAFT00000032396 | ADGRL4 | 1,21 | 4,62E-03 | Not filtered |
| 101 | ENSCAFT00000046656 | CX3CR1 | 1,07 | 4,62E-03 | Not filtered |
| 102 | ENSCAFT00000001409 | COL11A2 | 1,12 | 4,64E-03 | Not filtered |
| 103 | ENSCAFT00000036013 | MST1R | -0,87 | 4,74E-03 | Not filtered |
| 104 | ENSCAFT00000029594 | TENM1 | 0,90 | 4,74E-03 | Not filtered |
| 105 | ENSCAFT00000006096 | RAB3C | 0,90 | 4,78E-03 | Not filtered |
| **106** | **ENSCAFT00000027126** | **NUPR1** | **-1,35** | **4,94E-03** | **93^rd^** |
| **107** | **ENSCAFT00000009052** | **P2RY2** | **-1,67** | **5,05E-03** | **92^nd^** |
| 108 | ENSCAFT00000048687 | GRIK1 | 0,88 | 5,56E-03 | Not filtered |
| 109 | ENSCAFT00000050122 |  | -1,02 | 5,56E-03 | Not filtered |
| **110** | **ENSCAFT00000019347** |  | **-1,03** | **5,56E-03** | **91^st^** |
| **111** | **ENSCAFT00000039881** |  | **-1,39** | **5,56E-03** | **98^th^** |
| 112 | ENSCAFT00000029596 | TENM1 | 0,98 | 5,61E-03 | Not filtered |
| 113 | ENSCAFT00000047013 |  | 1,54 | 5,64E-03 | Not filtered |
| 114 | ENSCAFT00000020574 | STPG1 | 1,33 | 5,64E-03 | Not filtered |
| 115 | ENSCAFT00000005594 | STK17A | 1,16 | 5,64E-03 | Not filtered |
| 116 | ENSCAFT00000021648 |  | 4,82 | 5,69E-03 | Not filtered |
| 117 | ENSCAFT00000008732 |  | -3,07 | 5,69E-03 | Not filtered |
| 118 | ENSCAFT00000005257 | CDCA8 | -0,81 | 5,79E-03 | Not filtered |
| 119 | ENSCAFT00000023962 | KCNN1 | -1,05 | 5,79E-03 | Not filtered |
| 120 | ENSCAFT00000021514 |  | -3,48 | 5,79E-03 | Not filtered |
| 121 | ENSCAFT00000020206 | EPS8 | 0,96 | 6,06E-03 | Not filtered |
| 122 | ENSCAFT00000043005 | CCDC3 | 0,83 | 6,06E-03 | Not filtered |
| 123 | ENSCAFT00000031740 | TLDC1 | -0,83 | 6,06E-03 | Not filtered |
| 124 | ENSCAFT00000048547 | MRPL34 | -1,02 | 6,06E-03 | Not filtered |
| 125 | ENSCAFT00000012229 | MPEG1 | 1,43 | 6,49E-03 | Not filtered |
| 126 | ENSCAFT00000043964 | JADE2 | -0,56 | 6,62E-03 | Not filtered |
| 127 | ENSCAFT00000026974 | EME1 | -0,95 | 6,62E-03 | Not filtered |
| 128 | ENSCAFT00000015805 | STMND1 | -1,18 | 6,62E-03 | Not filtered |
| 129 | ENSCAFT00000038122 | OOEP | 4,44 | 6,84E-03 | Not filtered |
| 130 | ENSCAFT00000019796 | CASQ1 | 2,04 | 7,05E-03 | Not filtered |
| 131 | ENSCAFT00000021851 |  | -0,68 | 7,05E-03 | Not filtered |
| **132** | **ENSCAFT00000024572** | **SCG3** | **0,92** | **7,47E-03** | **98^th^** |
| 133 | ENSCAFT00000027202 | SHE | 0,69 | 7,53E-03 | Not filtered |
| 134 | ENSCAFT00000049656 | ABCG2 | 1,01 | 7,58E-03 | Not filtered |
| 135 | ENSCAFT00000005878 | KEL | -1,05 | 7,66E-03 | Not filtered |
| 136 | ENSCAFT00000031659 | CA5A | 1,38 | 7,89E-03 | Not filtered |
| 137 | ENSCAFT00000016783 | MTTP | 1,20 | 7,89E-03 | Not filtered |
| 138 | ENSCAFT00000022236 | AIFM2 | -0,80 | 7,89E-03 | Not filtered |
| 139 | ENSCAFT00000018222 | PITPNC1 | 1,04 | 8,07E-03 | Not filtered |
| **140** | **ENSCAFT00000018127** |  | **-0,70** | **8,07E-03** | **96^th^** |
| 141 | ENSCAFT00000000397 | MAP3K5 | 0,51 | 8,08E-03 | Not filtered |
| 142 | ENSCAFT00000013222 | TM4SF18 | 1,08 | 8,34E-03 | Not filtered |
| 143 | ENSCAFT00000000298 | SLC18B1 | 0,83 | 8,38E-03 | Not filtered |
| 144 | ENSCAFT00000049200 | TRIB2 | 0,62 | 8,40E-03 | Not filtered |
| 145 | ENSCAFT00000012150 | CD180 | 1,26 | 9,10E-03 | Not filtered |
| 146 | ENSCAFT00000049322 | FEN1 | -0,70 | 9,10E-03 | Not filtered |
| 147 | ENSCAFT00000015494 | ELOVL2 | 1,11 | 9,37E-03 | Not filtered |
| 148 | ENSCAFT00000039489 | SFRP4 | -1,14 | 9,45E-03 | Not filtered |
| 149 | ENSCAFT00000046208 | SNAI3 | -1,72 | 9,47E-03 | Not filtered |
| 150 | ENSCAFT00000046049 | ITGA6 | 1,26 | 9,59E-03 | Not filtered |
| 151 | ENSCAFT00000001083 | ANGPT1 | 1,07 | 9,64E-03 | Not filtered |
| 152 | ENSCAFT00000008784 | CLDN10 | 0,73 | 9,64E-03 | Not filtered |
| 153 | ENSCAFT00000032339 | CDH3 | -0,90 | 9,88E-03 | Not filtered |
| 154 | ENSCAFT00000008856 | SUPT5H | -0,45 | 1,02E-02 | Not filtered |
| 155 | ENSCAFT00000006034 | SLC7A11 | 1,32 | 1,04E-02 | Not filtered |
| 156 | ENSCAFT00000011237 | RAB3C | 0,80 | 1,08E-02 | Not filtered |
| 157 | ENSCAFT00000014333 | CALB1 | 2,48 | 1,10E-02 | Not filtered |
| 158 | ENSCAFT00000010417 | ELK3 | 0,68 | 1,10E-02 | Not filtered |
| 159 | ENSCAFT00000008294 | VHL | 0,56 | 1,10E-02 | Not filtered |
| 160 | ENSCAFT00000043101 | EMP1 | 0,74 | 1,11E-02 | Not filtered |
| 161 | ENSCAFT00000001042 | ZBTB12 | -0,70 | 1,11E-02 | Not filtered |
| 162 | ENSCAFT00000006826 | ATP6V0A4 | 1,48 | 1,12E-02 | Not filtered |
| 163 | ENSCAFT00000046164 |  | -3,07 | 1,14E-02 | Not filtered |
| **164** | **ENSCAFT00000014467** | **BHMT** | **1,10** | **1,17E-02** | **95^th^** |
| 165 | ENSCAFT00000022939 | MAOA | 0,77 | 1,18E-02 | Not filtered |
| 166 | ENSCAFT00000012133 | DPY19L3 | 0,57 | 1,21E-02 | Not filtered |
| 167 | ENSCAFT00000030449 | PNMA6A | -0,99 | 1,21E-02 | Not filtered |
| 168 | ENSCAFT00000010017 | CCNA1 | -0,67 | 1,21E-02 | Not filtered |
| 169 | ENSCAFT00000045654 | PPP2R1B | 0,62 | 1,24E-02 | Not filtered |
| 170 | ENSCAFT00000026017 | SEL1L3 | 0,54 | 1,24E-02 | Not filtered |
| 171 | ENSCAFT00000047154 | CYYR1 | 1,02 | 1,25E-02 | Not filtered |
| **172** | **ENSCAFT00000008624** | **EHD3** | **-0,52** | **1,27E-02** | **93^rd^** |
| 173 | ENSCAFT00000000346 | TRIM36 | 0,69 | 1,27E-02 | Not filtered |
| 174 | ENSCAFT00000032167 | LRRC8C | 0,68 | 1,34E-02 | Not filtered |
| 175 | ENSCAFT00000016073 | BMPR1B | 1,31 | 1,35E-02 | Not filtered |
| 176 | ENSCAFT00000028869 | LAMA3 | -1,60 | 1,38E-02 | Not filtered |
| 177 | ENSCAFT00000024392 | TNFSF10 | 1,07 | 1,39E-02 | Not filtered |
| 178 | ENSCAFT00000027898 | GDPD1 | 0,60 | 1,39E-02 | Not filtered |
| **179** | **ENSCAFT00000013168** | **GPD1** | **-0,89** | **1,43E-02** | **94^th^** |
| 180 | ENSCAFT00000014475 | BHMT2 | 1,11 | 1,43E-02 | Not filtered |
| 181 | ENSCAFT00000043020 | FNDC1 | 1,03 | 1,43E-02 | Not filtered |
| 182 | ENSCAFT00000044792 | DIO2 | 1,74 | 1,46E-02 | Not filtered |
| **183** | **ENSCAFT00000031879** | **VAT1L** | **0,86** | **1,46E-02** | **95^th^** |
| 184 | ENSCAFT00000030320 |  | -1,00 | 1,46E-02 | Not filtered |
| 185 | ENSCAFT00000017406 | NPNT | 1,16 | 1,46E-02 | Not filtered |
| 186 | ENSCAFT00000044761 | BDKRB2 | -2,62 | 1,46E-02 | Not filtered |
| 187 | ENSCAFT00000039527 | C3AR1 | 1,13 | 1,51E-02 | Not filtered |
| 188 | ENSCAFT00000044109 |  | -0,75 | 1,51E-02 | Not filtered |
| 189 | ENSCAFT00000027222 | GSTZ1 | -0,85 | 1,51E-02 | Not filtered |
| **190** | **ENSCAFT00000023993** | **QDPR** | **-1,16** | **1,51E-02** | **99^th^** |
| 191 | ENSCAFT00000049832 | STK17A | 1,14 | 1,51E-02 | Not filtered |
| 192 | ENSCAFT00000010805 | NLRP14 | 1,49 | 1,53E-02 | Not filtered |
| **193** | **ENSCAFT00000021914** | **TMEM47** | **1,07** | **1,58E-02** | **95^th^** |
| 194 | ENSCAFT00000014470 | ADAM28 | 0,95 | 1,58E-02 | Not filtered |
| **195** | **ENSCAFT00000017013** | **ADD3** | **0,78** | **1,58E-02** | **94^th^** |
| 196 | ENSCAFT00000026818 | SLC35B1 | -0,43 | 1,58E-02 | Not filtered |
| 197 | ENSCAFT00000032475 | CES2 | -1,15 | 1,58E-02 | Not filtered |
| 198 | ENSCAFT00000045856 | VSX2 | -1,25 | 1,58E-02 | Not filtered |
| 199 | ENSCAFT00000021879 | CCR5 | 1,48 | 1,59E-02 | Not filtered |
| **200** | **ENSCAFT00000008101** | **SLC6A1** | **0,58** | **1,61E-02** | **95^th^** |
| 201 | ENSCAFT00000046233 | LTBP2 | -2,26 | 1,67E-02 | Not filtered |
| 202 | ENSCAFT00000006384 | PIK3CG | 0,99 | 1,68E-02 | Not filtered |
| **203** | **ENSCAFT00000027435** |  | **-0,53** | **1,70E-02** | **91^st^** |
| 204 | ENSCAFT00000015321 | ABCG2 | 0,86 | 1,70E-02 | Not filtered |
| **205** | **ENSCAFT00000017616** |  | **-0,87** | **1,72E-02** | **93^rd^** |
| **206** | **ENSCAFT00000018912** | **PCBP3** | **-1,50** | **1,74E-02** | **98^th^** |
| 207 | ENSCAFT00000049390 | MOV10L1 | 1,12 | 1,86E-02 | Not filtered |
| 208 | ENSCAFT00000048622 | SUN2 | -0,48 | 1,87E-02 | Not filtered |
| 209 | ENSCAFT00000012270 | SMAD1 | 0,58 | 1,87E-02 | Not filtered |
| 210 | ENSCAFT00000002736 | CHRNA5 | 1,11 | 1,88E-02 | Not filtered |
| 211 | ENSCAFT00000004046 | COL19A1 | 0,90 | 1,88E-02 | Not filtered |
| 212 | ENSCAFT00000008744 | SLCO2B1 | 0,72 | 1,88E-02 | Not filtered |
| 213 | ENSCAFT00000016720 | PKNOX1 | -0,69 | 1,88E-02 | Not filtered |
| 214 | ENSCAFT00000048536 | POLN | -0,86 | 1,88E-02 | Not filtered |
| 215 | ENSCAFT00000036890 |  | -3,58 | 1,88E-02 | Not filtered |
| 216 | ENSCAFT00000048492 | CENPW | 1,21 | 1,90E-02 | Not filtered |
| **217** | **ENSCAFT00000000986** |  | **-1,86** | **1,90E-02** | **99^th^** |
| 218 | ENSCAFT00000026830 | LTBP2 | -2,12 | 1,90E-02 | Not filtered |
| 219 | ENSCAFT00000017178 | SLC9B2 | 0,98 | 1,95E-02 | Not filtered |
| 220 | ENSCAFT00000032225 | DHX38 | -0,35 | 1,95E-02 | Not filtered |
| **221** | **ENSCAFT00000043488** | **CSPG5** | **0,72** | **1,97E-02** | **92^nd^** |
| 222 | ENSCAFT00000009440 | TBC1D5 | 0,55 | 1,97E-02 | Not filtered |
| 223 | ENSCAFT00000023694 | PDE4A | -0,62 | 1,97E-02 | Not filtered |
| 224 | ENSCAFT00000007481 | FADS6 | -0,74 | 2,01E-02 | Not filtered |
| 225 | ENSCAFT00000001085 | RSPO2 | 0,90 | 2,04E-02 | Not filtered |
| **226** | **ENSCAFT00000000984** |  | **-1,80** | **2,12E-02** | **99^th^** |
| 227 | ENSCAFT00000043813 | PGAP1 | 1,18 | 2,13E-02 | Not filtered |
| 228 | ENSCAFT00000029073 | PPP1R13B | -0,51 | 2,23E-02 | Not filtered |
| 229 | ENSCAFT00000036549 |  | 0,71 | 2,24E-02 | Not filtered |
| 230 | ENSCAFT00000045214 | NHSL1 | 0,79 | 2,28E-02 | Not filtered |
| 231 | ENSCAFT00000005411 | CSF3R | -1,93 | 2,35E-02 | Not filtered |
| 232 | ENSCAFT00000038735 |  | 0,67 | 2,36E-02 | Not filtered |
| 233 | ENSCAFT00000012461 | OCLN | 1,32 | 2,37E-02 | Not filtered |
| **234** | **ENSCAFT00000043311** |  | **-1,01** | **2,37E-02** | **99^th^** |
| **235** | **ENSCAFT00000000831** | **VIP** | **0,76** | **2,37E-02** | **91^st^** |
| 236 | ENSCAFT00000045921 |  | 1,02 | 2,38E-02 | Not filtered |
| 237 | ENSCAFT00000038891 | JAM2 | 0,74 | 2,38E-02 | Not filtered |
| 238 | ENSCAFT00000000665 | CPM | -1,15 | 2,38E-02 | Not filtered |
| **239** | **ENSCAFT00000002268** | **MEST** | **0,96** | **2,39E-02** | **93^rd^** |
| 240 | ENSCAFT00000008331 | MYO7A | 0,82 | 2,40E-02 | Not filtered |
| 241 | ENSCAFT00000015657 | PPM1M | -0,62 | 2,46E-02 | Not filtered |
| 242 | ENSCAFT00000048836 | ZNF471 | 1,22 | 2,48E-02 | Not filtered |
| 243 | ENSCAFT00000003615 | ECM2 | 1,09 | 2,48E-02 | Not filtered |
| 244 | ENSCAFT00000031573 | ACSF3 | -0,49 | 2,48E-02 | Not filtered |
| 245 | ENSCAFT00000045753 | PDZRN4 | 0,85 | 2,50E-02 | Not filtered |
| 246 | ENSCAFT00000029091 | ALKBH5 | -0,83 | 2,51E-02 | Not filtered |
| 247 | ENSCAFT00000020583 | PCYT1A | 0,48 | 2,52E-02 | Not filtered |
| 248 | ENSCAFT00000012231 | PREX2 | 1,11 | 2,56E-02 | Not filtered |
| 249 | ENSCAFT00000045600 | ZDBF2 | 0,87 | 2,56E-02 | Not filtered |
| 250 | ENSCAFT00000047906 |  | -1,56 | 2,56E-02 | Not filtered |
| 251 | ENSCAFT00000023165 | PYGL | -0,59 | 2,57E-02 | Not filtered |
| 252 | ENSCAFT00000044729 | GPR179 | 0,67 | 2,57E-02 | Not filtered |
| 253 | ENSCAFT00000049677 | PNPLA4 | 1,42 | 2,58E-02 | Not filtered |
| 254 | ENSCAFT00000044927 |  | 0,73 | 2,58E-02 | Not filtered |
| 255 | ENSCAFT00000013389 | PAG1 | 0,65 | 2,58E-02 | Not filtered |
| 256 | ENSCAFT00000024467 | KIAA0753 | -0,55 | 2,58E-02 | Not filtered |
| 257 | ENSCAFT00000036074 |  | -0,67 | 2,64E-02 | Not filtered |
| 258 | ENSCAFT00000016681 | SLC19A3 | 0,98 | 2,67E-02 | Not filtered |
| 259 | ENSCAFT00000049634 |  | 3,88 | 2,67E-02 | Not filtered |
| 260 | ENSCAFT00000021268 |  | 1,64 | 2,67E-02 | Not filtered |
| 261 | ENSCAFT00000024816 | GNB5 | -0,51 | 2,68E-02 | Not filtered |
| 262 | ENSCAFT00000031839 | CDC14A | 1,02 | 2,70E-02 | Not filtered |
| 263 | ENSCAFT00000002695 | C11H9orf72 | 0,71 | 2,72E-02 | Not filtered |
| 264 | ENSCAFT00000015985 | LZTS1 | -0,46 | 2,72E-02 | Not filtered |
| 265 | ENSCAFT00000002677 | TEK | 0,86 | 2,75E-02 | Not filtered |
| 266 | ENSCAFT00000008545 | SOCS3 | -1,50 | 2,76E-02 | Not filtered |
| 267 | ENSCAFT00000047656 | RGR | 2,11 | 2,79E-02 | Not filtered |
| 268 | ENSCAFT00000049541 | GRAMD2B | 0,65 | 2,79E-02 | Not filtered |
| 269 | ENSCAFT00000009396 | KCNH8 | -0,90 | 2,82E-02 | Not filtered |
| 270 | ENSCAFT00000046228 | COL25A1 | 1,14 | 2,84E-02 | Not filtered |
| **271** | **ENSCAFT00000013466** |  | **1,10** | **2,84E-02** | **93^rd^** |
| 272 | ENSCAFT00000000481 | LRIG3 | -0,76 | 2,84E-02 | Not filtered |
| 273 | ENSCAFT00000031177 | RPUSD1 | -0,78 | 2,84E-02 | Not filtered |
| 274 | ENSCAFT00000050051 |  | 4,64 | 2,86E-02 | Not filtered |
| **275** | **ENSCAFT00000001481** | **GJA1** | **1,01** | **2,88E-02** | **98^th^** |
| 276 | ENSCAFT00000031814 | COL11A2 | 0,80 | 2,88E-02 | Not filtered |
| 277 | ENSCAFT00000018339 | ABLIM1 | 0,62 | 2,88E-02 | Not filtered |
| 278 | ENSCAFT00000048682 |  | -0,58 | 2,88E-02 | Not filtered |
| 279 | ENSCAFT00000031391 | KCNT1 | -0,64 | 2,88E-02 | Not filtered |
| 280 | ENSCAFT00000001158 | CERK | -0,68 | 2,88E-02 | Not filtered |
| 281 | ENSCAFT00000044626 | MYH13 | -1,31 | 2,89E-02 | Not filtered |
| 282 | ENSCAFT00000016709 |  | -3,71 | 2,90E-02 | Not filtered |
| **283** | **ENSCAFT00000022271** | **LANCL1** | **0,58** | **2,92E-02** | **96^th^** |
| 284 | ENSCAFT00000011905 | SLC7A10 | 1,22 | 2,95E-02 | Not filtered |
| 285 | ENSCAFT00000047021 | NIPAL1 | 0,96 | 2,95E-02 | Not filtered |
| 286 | ENSCAFT00000027279 | ZDHHC15 | 0,76 | 2,95E-02 | Not filtered |
| 287 | ENSCAFT00000010231 | KCTD16 | 0,98 | 2,98E-02 | Not filtered |
| 288 | ENSCAFT00000007517 | ITGA8 | 0,91 | 2,98E-02 | Not filtered |
| **289** | **ENSCAFT00000011978** | **RILPL1** | **-0,71** | **2,98E-02** | **98^th^** |
| 290 | ENSCAFT00000043178 | PTPDC1 | -0,99 | 3,02E-02 | Not filtered |
| 291 | ENSCAFT00000012618 | RHOBTB3 | 0,93 | 3,03E-02 | Not filtered |
| 292 | ENSCAFT00000020084 | DERA | 0,62 | 3,03E-02 | Not filtered |
| 293 | ENSCAFT00000023491 | GULP1 | 0,92 | 3,11E-02 | Not filtered |
| 294 | ENSCAFT00000011507 | PTPRG | 0,66 | 3,11E-02 | Not filtered |
| **295** | **ENSCAFT00000018895** | **BZW1** | **0,66** | **3,11E-02** | **96^th^** |
| 296 | ENSCAFT00000044975 | JADE2 | -0,65 | 3,11E-02 | Not filtered |
| 297 | ENSCAFT00000012478 | RETSAT | 0,67 | 3,14E-02 | Not filtered |
| 298 | ENSCAFT00000009653 | MGAT4C | 0,82 | 3,18E-02 | Not filtered |
| 299 | ENSCAFT00000043408 | GTF3C3 | 0,72 | 3,20E-02 | Not filtered |
| 300 | ENSCAFT00000004517 | NFYC | -0,65 | 3,30E-02 | Not filtered |
| 301 | ENSCAFT00000050102 |  | 1,52 | 3,39E-02 | Not filtered |
| **302** | **ENSCAFT00000046855** | **CERK** | **-0,59** | **3,39E-02** | **90^th^** |
| 303 | ENSCAFT00000049206 |  | -1,37 | 3,41E-02 | Not filtered |
| 304 | ENSCAFT00000003755 | ANKRD39 | -0,73 | 3,41E-02 | Not filtered |
| 305 | ENSCAFT00000049378 |  | -0,86 | 3,53E-02 | Not filtered |
| 306 | ENSCAFT00000027762 | PAQR5 | 1,00 | 3,53E-02 | Not filtered |
| 307 | ENSCAFT00000002737 | CHRNA5 | 0,89 | 3,53E-02 | Not filtered |
| 308 | ENSCAFT00000026321 | MSN | 0,52 | 3,53E-02 | Not filtered |
| 309 | ENSCAFT00000003267 | ADGRF5 | 0,87 | 3,56E-02 | Not filtered |
| 310 | ENSCAFT00000007981 | PAK3 | 0,60 | 3,56E-02 | Not filtered |
| 311 | ENSCAFT00000037089 | DPY19L3 | 0,60 | 3,59E-02 | Not filtered |
| 312 | ENSCAFT00000003998 | P3H1 | -0,72 | 3,62E-02 | Not filtered |
| 313 | ENSCAFT00000049967 | SERPINB9 | 0,69 | 3,63E-02 | Not filtered |
| 314 | ENSCAFT00000000818 | SNCAIP | 0,63 | 3,63E-02 | Not filtered |
| 315 | ENSCAFT00000042869 | TFAP4 | -0,82 | 3,63E-02 | Not filtered |
| 316 | ENSCAFT00000015365 | WNT8B | -2,98 | 3,63E-02 | Not filtered |
| **317** | **ENSCAFT00000046391** | **RANBP3L** | **1,37** | **3,63E-02** | **91^st^** |
| **318** | **ENSCAFT00000029168** | **TMEM98** | **-0,79** | **3,66E-02** | **95^th^** |
| 319 | ENSCAFT00000028199 |  | -4,63 | 3,70E-02 | Not filtered |
| 320 | ENSCAFT00000000638 | PLEKHG1 | 0,59 | 3,71E-02 | Not filtered |
| 321 | ENSCAFT00000037219 |  | 0,75 | 3,72E-02 | Not filtered |
| 322 | ENSCAFT00000024144 | SNAP29 | 0,50 | 3,72E-02 | Not filtered |
| 323 | ENSCAFT00000019616 | HMBS | -0,62 | 3,75E-02 | Not filtered |
| 324 | ENSCAFT00000010343 | AMDHD1 | -0,71 | 3,79E-02 | Not filtered |
| 325 | ENSCAFT00000046491 | RHBDL3 | -1,08 | 3,79E-02 | Not filtered |
| 326 | ENSCAFT00000035378 |  | -2,02 | 3,79E-02 | Not filtered |
| 327 | ENSCAFT00000025789 | PGM2 | 0,52 | 3,80E-02 | Not filtered |
| 328 | ENSCAFT00000045572 | GHR | 0,72 | 3,81E-02 | Not filtered |
| 329 | ENSCAFT00000046022 | TCF23 | 2,05 | 3,82E-02 | Not filtered |
| **330** | **ENSCAFT00000024665** | **TMOD2** | **0,55** | **3,83E-02** | **95^th^** |
| 331 | ENSCAFT00000002700 | TOM1 | -0,67 | 3,88E-02 | Not filtered |
| 332 | ENSCAFT00000046313 | CCR6 | 2,57 | 3,88E-02 | Not filtered |
| **333** | **ENSCAFT00000009471** | **ANAPC11** | **-0,98** | **3,89E-02** | **97^th^** |
| **334** | **ENSCAFT00000023016** | **IGFBP2** | **-1,12** | **3,89E-02** | **92^nd^** |
| **335** | **ENSCAFT00000045335** | **PTPRZ1** | **1,32** | **3,91E-02** | **93^rd^** |
| 336 | ENSCAFT00000049097 | PPA2 | 0,73 | 3,98E-02 | Not filtered |
| 337 | ENSCAFT00000043409 |  | 1,64 | 3,98E-02 | Not filtered |
| **338** | **ENSCAFT00000045219** | **ABLIM1** | **0,49** | **3,98E-02** | **90^th^** |
| 339 | ENSCAFT00000044179 | NHS | 0,95 | 4,00E-02 | Not filtered |
| **340** | **ENSCAFT00000030037** | **DEXI** | **-1,27** | **4,01E-02** | **96^th^** |
| 341 | ENSCAFT00000023864 | KIF17 | -0,68 | 4,01E-02 | Not filtered |
| 342 | ENSCAFT00000045914 |  | -1,60 | 4,01E-02 | Not filtered |
| **343** | **ENSCAFT00000025184** | **MED11** | **-0,88** | **4,04E-02** | **90^th^** |
| **344** | **ENSCAFT00000019956** | **SH3BGRL3** | **-1,12** | **4,04E-02** | **99^th^** |
| 345 | ENSCAFT00000019633 | SLCO1A2 | 0,95 | 4,06E-02 | Not filtered |
| 346 | ENSCAFT00000010087 | DHX57 | 0,63 | 4,06E-02 | Not filtered |
| **347** | **ENSCAFT00000003721** | **GRHPR** | **-0,76** | **4,06E-02** | **91^st^** |
| 348 | ENSCAFT00000009843 | POSTN | 1,42 | 4,07E-02 | Not filtered |
| **349** | **ENSCAFT00000038326** | **SQLE** | **0,55** | **4,07E-02** | **90^th^** |
| 350 | ENSCAFT00000026389 | ADAM21 | -0,95 | 4,12E-02 | Not filtered |
| **351** | **ENSCAFT00000023712** | **CYP27A1** | **-1,21** | **4,12E-02** | **94^th^** |
| 352 | ENSCAFT00000032397 | ADGRL4 | 1,07 | 4,13E-02 | Not filtered |
| 353 | ENSCAFT00000000847 | RGS17 | 0,83 | 4,13E-02 | Not filtered |
| 354 | ENSCAFT00000029649 | SMARCA1 | 0,78 | 4,14E-02 | Not filtered |
| 355 | ENSCAFT00000017004 | SLC39A8 | 0,66 | 4,16E-02 | Not filtered |
| 356 | ENSCAFT00000032451 | E2F4 | -0,52 | 4,22E-02 | Not filtered |
| 357 | ENSCAFT00000003269 | ADGRF5 | 0,72 | 4,23E-02 | Not filtered |
| 358 | ENSCAFT00000045054 | EPS8 | 0,94 | 4,24E-02 | Not filtered |
| **359** | **ENSCAFT00000047366** | **SEC23A** | **0,73** | **4,24E-02** | **90^th^** |
| 360 | ENSCAFT00000049670 |  | -0,78 | 4,24E-02 | Not filtered |
| 361 | ENSCAFT00000043666 |  | -1,85 | 4,25E-02 | Not filtered |
| 362 | ENSCAFT00000044299 | KDM1A | -0,31 | 4,26E-02 | Not filtered |
| 363 | ENSCAFT00000031183 | KLHL21 | -0,72 | 4,26E-02 | Not filtered |
| 364 | ENSCAFT00000004714 |  | 0,87 | 4,26E-02 | Not filtered |
| 365 | ENSCAFT00000008283 | FNDC4 | -0,63 | 4,26E-02 | Not filtered |
| **366** | **ENSCAFT00000006737** |  | **-0,62** | **4,26E-02** | **96th** |
| 367 | ENSCAFT00000043067 | KIF17 | -0,74 | 4,26E-02 | Not filtered |
| **368** | **ENSCAFT00000011343** |  | **-1,39** | **4,26E-02** | **95^th^** |
| 369 | ENSCAFT00000022161 | SYTL5 | 1,36 | 4,27E-02 | Not filtered |
| 370 | ENSCAFT00000000089 | ITGA7 | 0,76 | 4,27E-02 | Not filtered |
| **371** | **ENSCAFT00000007160** | **FOSB** | **-1,08** | **4,27E-02** | **93^rd^** |
| 372 | ENSCAFT00000024296 | CD38 | 1,00 | 4,28E-02 | Not filtered |
| 373 | ENSCAFT00000023745 | ZSCAN25 | -0,61 | 4,28E-02 | Not filtered |
| **374** | **ENSCAFT00000018282** | **AIP** | **-0,72** | **4,28E-02** | **94^th^** |
| 375 | ENSCAFT00000017419 | ACSL5 | 1,05 | 4,29E-02 | Not filtered |
| 376 | ENSCAFT00000025658 | FAM114A1 | 0,76 | 4,29E-02 | Not filtered |
| 377 | ENSCAFT00000016750 | OASL | -0,40 | 4,30E-02 | Not filtered |
| **378** | **ENSCAFT00000008137** | **RNF157** | **-0,39** | **4,32E-02** | **94^th^** |
| 379 | ENSCAFT00000029520 | LAMA1 | 0,95 | 4,34E-02 | Not filtered |
| 380 | ENSCAFT00000047858 | PCSK5 | 0,66 | 4,35E-02 | Not filtered |
| **381** | **ENSCAFT00000029484** |  | **0,64** | **4,35E-02** | **96^th^** |
| 382 | ENSCAFT00000020806 | SART1 | -0,46 | 4,35E-02 | Not filtered |
| 383 | ENSCAFT00000004933 | NOD1 | -0,70 | 4,35E-02 | Not filtered |
| 384 | ENSCAFT00000003500 | PTPDC1 | -1,01 | 4,35E-02 | Not filtered |
| **385** | **ENSCAFT00000029354** |  | **-2,51** | **4,37E-02** | **92^nd^** |
| 386 | ENSCAFT00000038139 | SYCE1 | 3,29 | 4,42E-02 | Not filtered |
| 387 | ENSCAFT00000046875 | PIK3C2A | 0,78 | 4,42E-02 | Not filtered |
| **388** | **ENSCAFT00000009006** | **NCCRP1** | **-3,50** | **4,42E-02** | **90^th^** |
| 389 | ENSCAFT00000014595 | GPR15 | -2,61 | 4,43E-02 | Not filtered |
| 390 | ENSCAFT00000044965 | EGLN2 | -0,77 | 4,43E-02 | Not filtered |
| **391** | **ENSCAFT00000045554** |  | **-2,60** | **4,46E-02** | **91^st^** |
| **392** | **ENSCAFT00000022594** | **NCAN** | **0,87** | **4,47E-02** | **93^rd^** |
| 393 | ENSCAFT00000007000 | GPR17 | 1,35 | 4,49E-02 | Not filtered |
| **394** | **ENSCAFT00000020636** | **ACE** | **1,31** | **4,49E-02** | **97^th^** |
| **395** | **ENSCAFT00000025424** | **AKAP5** | **0,76** | **4,49E-02** | **95^th^** |
| 396 | ENSCAFT00000027662 | MYH13 | -0,98 | 4,49E-02 | Not filtered |
| 397 | ENSCAFT00000023000 |  | -1,26 | 4,50E-02 | Not filtered |
| 398 | ENSCAFT00000022292 | CDCP1 | 0,83 | 4,56E-02 | Not filtered |
| 399 | ENSCAFT00000023561 | DNM3 | 0,49 | 4,56E-02 | Not filtered |
| 400 | ENSCAFT00000001623 | TXNDC15 | 0,44 | 4,56E-02 | Not filtered |
| 401 | ENSCAFT00000028088 |  | -0,36 | 4,56E-02 | Not filtered |
| 402 | ENSCAFT00000019444 | MLPH | -2,01 | 4,56E-02 | Not filtered |
| 403 | ENSCAFT00000002939 | TRPM3 | 0,94 | 4,64E-02 | Not filtered |
| 404 | ENSCAFT00000015989 | ANO5 | 0,74 | 4,64E-02 | Not filtered |
| 405 | ENSCAFT00000012249 | C29H8orf34 | 0,90 | 4,66E-02 | Not filtered |
| 406 | ENSCAFT00000008063 | CCDC97 | -0,56 | 4,66E-02 | Not filtered |
| **407** | **ENSCAFT00000010234** | **LAMP1** | **-0,32** | **4,68E-02** | **96^th^** |
| 408 | ENSCAFT00000023805 | MAPK1IP1L | 0,51 | 4,72E-02 | Not filtered |
| 409 | ENSCAFT00000013457 | BLNK | 1,08 | 4,72E-02 | Not filtered |
| 410 | ENSCAFT00000039684 |  | 0,83 | 4,72E-02 | Not filtered |
| 411 | ENSCAFT00000022836 | ARHGEF10 | -0,75 | 4,72E-02 | Not filtered |
| 412 | ENSCAFT00000029569 | CACHD1 | 0,54 | 4,75E-02 | Not filtered |
| 413 | ENSCAFT00000012751 | ZNF366 | 0,93 | 4,75E-02 | Not filtered |
| 414 | ENSCAFT00000018076 | ACTL6A | 0,73 | 4,76E-02 | Not filtered |
| 415 | ENSCAFT00000004830 | SUSD1 | 0,69 | 4,76E-02 | Not filtered |
| 416 | ENSCAFT00000011640 |  | 0,73 | 4,78E-02 | Not filtered |
| 417 | ENSCAFT00000045071 | TIPARP | 0,96 | 4,80E-02 | Not filtered |
| **418** | **ENSCAFT00000024554** | **DES** | **-1,26** | **4,81E-02** | **91^st^** |
| 419 | ENSCAFT00000021954 | SEC23A | 0,53 | 4,83E-02 | Not filtered |
| 420 | ENSCAFT00000035362 | RCCD1 | -0,78 | 4,83E-02 | Not filtered |
| 421 | ENSCAFT00000005238 | ARHGAP25 | 0,86 | 4,83E-02 | Not filtered |
| 422 | ENSCAFT00000049648 |  | -1,05 | 4,85E-02 | Not filtered |
| 423 | ENSCAFT00000013538 | P2RY12 | 0,93 | 4,87E-02 | Not filtered |
| 424 | ENSCAFT00000043524 | COL19A1 | 0,75 | 4,87E-02 | Not filtered |
| 425 | ENSCAFT00000000563 | GNS | 0,51 | 4,87E-02 | Not filtered |
| 426 | ENSCAFT00000020997 | PISD | -0,55 | 4,87E-02 | Not filtered |
| **427** | **ENSCAFT00000024280** | **KCNA1** | **-0,66** | **4,88E-02** | **90^th^** |
| **428** | **ENSCAFT00000017551** | **SLC25A6** | **-0,65** | **4,89E-02** | **98^th^** |
| **429** | **ENSCAFT00000046388** |  | **-1,37** | **4,91E-02** | **99^th^** |
| 430 | ENSCAFT00000044007 | ELN | 1,19 | 4,95E-02 | Not filtered |
|  | **ENSCAFT00000023394** | **FBN1** |  |  | **Added 97^th^,95^th^,90^th^** |
|  | **ENSCAFT00000009676** | **LHFPL6** |  |  | **Added 97^th^,95^th^,90^th^** |
|  | **ENSCAFT00000008870** | **PFKP** |  |  | **Added 97^th^,95^th^** |
|  | **ENSCAFT00000047460** | **ANKRD31** |  |  | **Added 97^th^,95^th^,90^th^** |
|  | **ENSCAFT00000012552** |  |  |  | **Added 97^th^,95^th^,90^th^** |
|  | **ENSCAFT00000048320** | **ZNHIT1** |  |  | **Added 97^th^,95^th^,90^th^** |
|  | **ENSCAFT00000029069** | **PCYOX1L** |  |  | **Added 97^th^,95^th^,90^th^** |
|  | **ENSCAFT00000007457** |  |  |  | **Added 97^th^,95^th^** |
|  | **ENSCAFT00000030988** | **HAGH** |  |  | **Added 97^th^** |
|  | **ENSCAFT00000031390** | **KCNT1** |  |  | **Added 97^th^,95^th^,90^th^** |
|  | **ENSCAFT00000039399** | **SLC25A28** |  |  | **Added 97^th^,95^th^,90^th^** |
|  | **ENSCAFT00000029752** | **RANBP3** |  |  | **Added 97^th^,95^th^,90^th^** |
|  | **ENSCAFT00000045978** | **TIMP3** |  |  | **Added 97^th^,95^th^** |
|  | **ENSCAFT00000026940** | **AURKB** |  |  | **Added 97^th^,95^th^,90^th^** |
|  | **ENSCAFT00000029071** | **XAB2** |  |  | **Added 97^th^,95^th^,90^th^** |
|  | **ENSCAFT00000049561** | **MRI1** |  |  | **Added 97^th^,95^th^,90^th^** |
|  | **ENSCAFT00000007204** | **XPC** |  |  | **Added 97^th^,95^th^,90^th^** |
|  | **ENSCAFT00000043301** | **COL11A1** |  |  | **Added 97^th^** |
|  | **ENSCAFT00000022098** | **FAM20B** |  |  | **Added 97^th^** |
|  | **ENSCAFT00000025496** | **LDB3** |  |  | **Added 97^th^,95^th^,90^th^** |
|  | **ENSCAFT00000007637** | **PHYH** |  |  | **Added 97^th^,95^th^,90^th^** |
|  | **ENSCAFT00000002119** | **SUN2** |  |  | **Added 95^th^,90^th^** |
|  | **ENSCAFT00000025645** | **EPHX1** |  |  | **Added 95^th^,90^th^** |
|  | **ENSCAFT00000026577** | **PDE6B** |  |  | **Added 95^th^,90^th^** |
|  | **ENSCAFT00000014889** | **C18H11orf94** |  |  | **Added 95^th^,90^th^** |
|  | **ENSCAFT00000036022** | **PHYH** |  |  | **Added 95^th^,90^th^** |
|  | **ENSCAFT00000020738** | **TPD52L2** |  |  | **Added 90^th^** |
|  | **ENSCAFT00000018896** | **POLG** |  |  | **Added 90^th^** |
|  | **ENSCAFT00000010783** | **SLCO2A1** |  |  | **Added 90^th^** |
|  | **ENSCAFT00000018755** | **CRYBA4** |  |  | **Added 90^th^** |
|  | **ENSCAFT00000017387** | **HFE** |  |  | **Added 90^th^** |
|  | **ENSCAFT00000028420** | **CLK3** |  |  | **Added 90^th^** |
