## Supplemental Table 4 for "On taming the effect of transcript level intra-condition count variation during differential expression analysis: a story of dogs, foxes and wolves"

| **Ranking** | **Transcript ID** | **Gene ID** | **log2 FC** | **p adj value** | **Filter level** |
| --- | --- | --- | --- | --- | --- |
| 1 | ENSCAFT00000001643 | TRIB1 | 0,91 | 2,82E-07 | Not-filtered |
| 2 | ENSCAFT00000012381 | PRIMPOL | -0,86 | 2,82E-07 | Not-filtered |
| 3 | ENSCAFT00000002697 | FSCN3 | -0,79 | 2,82E-07 | Not-filtered |
| 4 | ENSCAFT00000003134 | CCDC138 | -0,98 | 9,06E-06 | Not-filtered |
| 5 | ENSCAFT00000015069 | FARS2 | 0,58 | 2,20E-05 | Not-filtered |
| 6 | ENSCAFT00000032259 | NQO1 | 0,62 | 2,66E-05 | Not-filtered |
| 7 | ENSCAFT00000046785 |  | 1,10 | 5,56E-05 | Not-filtered |
| 8 | ENSCAFT00000009874 | DUSP6 | 0,67 | 5,56E-05 | Not-filtered |
| 9 | ENSCAFT00000047474 |  | 1,07 | 9,37E-05 | Not-filtered |
| **10** | **ENSCAFT00000001962** | **EGR1** | **0,86** | **1,72E-04** | **95^th^** |
| 11 | ENSCAFT00000015649 |  | 0,92 | 1,77E-04 | Not-filtered |
| 12 | ENSCAFT00000048992 |  | 1,11 | 1,85E-04 | Not-filtered |
| 13 | ENSCAFT00000035279 |  | 1,04 | 1,85E-04 | Not-filtered |
| 14 | ENSCAFT00000044393 | KHK | 1,03 | 1,85E-04 | Not-filtered |
| 15 | ENSCAFT00000045717 |  | 0,99 | 1,85E-04 | Not-filtered |
| 16 | ENSCAFT00000015525 |  | 0,98 | 1,85E-04 | Not-filtered |
| **17** | **ENSCAFT00000048012** | **NRSN1** | **0,77** | **1,85E-04** | **97^th^** |
| 18 | ENSCAFT00000029200 | ESM1 | -1,45 | 2,94E-04 | Not-filtered |
| 19 | ENSCAFT00000039205 |  | 0,99 | 3,11E-04 | Not-filtered |
| 20 | ENSCAFT00000043849 | EGR2 | 0,72 | 4,02E-04 | Not-filtered |
| **21** | **ENSCAFT00000031607** | **APRT** | **0,99** | **4,20E-04** | **96^th^** |
| 22 | ENSCAFT00000031690 | IRF8 | 0,40 | 4,20E-04 | Not-filtered |
| 23 | ENSCAFT00000038072 | SAPCD1 | -1,35 | 4,32E-04 | Not-filtered |
| 24 | ENSCAFT00000017242 |  | 1,12 | 4,96E-04 | Not-filtered |
| 25 | ENSCAFT00000015258 |  | -1,39 | 5,47E-04 | Not-filtered |
| 26 | ENSCAFT00000026984 | FOS | 1,03 | 5,65E-04 | Not-filtered |
| 27 | ENSCAFT00000031697 | C5H16orf74 | 1,09 | 5,67E-04 | Not-filtered |
| 28 | ENSCAFT00000008522 | SLITRK6 | -1,50 | 6,59E-04 | Not-filtered |
| 29 | ENSCAFT00000020697 | EGR2 | 0,67 | 8,43E-04 | Not-filtered |
| **30** | **ENSCAFT00000009043** | **BHLHE40** | **0,54** | **9,46E-04** | **91^st^** |
| 31 | ENSCAFT00000007514 | KHK | 0,84 | 1,00E-03 | Not-filtered |
| **32** | **ENSCAFT00000001517** | **NDUFA6** | **0,35** | **1,02E-03** | **93^rd^** |
| 33 | ENSCAFT00000011062 | DDX4 | -2,02 | 1,02E-03 | Not-filtered |
| 34 | ENSCAFT00000005919 | DKKL1 | 0,63 | 1,05E-03 | Not-filtered |
| 35 | ENSCAFT00000003135 | CCDC138 | -1,03 | 1,07E-03 | Not-filtered |
| **36** | **ENSCAFT00000043415** | **JUNB** | **1,11** | **1,23E-03** | **96^th^** |
| 37 | ENSCAFT00000018592 | INPP5D | 0,68 | 1,23E-03 | Not-filtered |
| 38 | ENSCAFT00000004310 |  | -1,91 | 1,23E-03 | Not-filtered |
| 39 | ENSCAFT00000011853 | TMEM232 | -1,08 | 1,47E-03 | Not-filtered |
| **40** | **ENSCAFT00000014839** | **BPHL** | **0,41** | **1,48E-03** | **91^st^** |
| **41** | **ENSCAFT00000027182** |  | **1,07** | **1,56E-03** | **94^th^** |
| 42 | ENSCAFT00000007767 | ATRAID | 0,62 | 1,67E-03 | Not-filtered |
| 43 | ENSCAFT00000009509 | RYR1 | -0,41 | 1,67E-03 | Not-filtered |
| 44 | ENSCAFT00000046725 |  | -1,91 | 1,69E-03 | Not-filtered |
| 45 | ENSCAFT00000016864 |  | 0,97 | 1,73E-03 | Not-filtered |
| 46 | ENSCAFT00000005238 | ARHGAP25 | 0,70 | 1,73E-03 | Not-filtered |
| 47 | ENSCAFT00000013583 | WDR54 | 0,55 | 1,79E-03 | Not-filtered |
| 48 | ENSCAFT00000007829 | ECHDC3 | 0,80 | 2,09E-03 | Not-filtered |
| 49 | ENSCAFT00000009853 | PCDHGA1 | 0,57 | 2,16E-03 | Not-filtered |
| **50** | **ENSCAFT00000028075** | **ARMCX3** | **0,32** | **2,19E-03** | **92^nd^** |
| 51 | ENSCAFT00000042883 | DUSP4 | 0,49 | 2,34E-03 | Not-filtered |
| **52** | **ENSCAFT00000001053** | **SKIV2L** | **0,87** | **2,62E-03** | **91^st^** |
| 53 | ENSCAFT00000037859 | YIPF1 | 0,39 | 2,84E-03 | Not-filtered |
| 54 | ENSCAFT00000028559 | PRPS1 | 0,47 | 3,05E-03 | Not-filtered |
| 55 | ENSCAFT00000011460 | FBLN7 | 0,54 | 3,25E-03 | Not-filtered |
| 56 | ENSCAFT00000006262 | HSD17B14 | 0,70 | 3,58E-03 | Not-filtered |
| 57 | ENSCAFT00000036730 | MAP2K3 | 0,69 | 3,58E-03 | Not-filtered |
| 58 | ENSCAFT00000007384 |  | 0,65 | 3,58E-03 | Not-filtered |
| 59 | ENSCAFT00000018894 | PTAFR | 0,61 | 3,58E-03 | Not-filtered |
| 60 | ENSCAFT00000031437 |  | 0,42 | 4,04E-03 | Not-filtered |
| **61** | **ENSCAFT00000046412** | **GNG4** | **0,79** | **4,35E-03** | **95^th^** |
| 62 | ENSCAFT00000015473 | GLB1L3 | -0,59 | 4,62E-03 | Not-filtered |
| 63 | ENSCAFT00000027070 | MYCBPAP | 0,81 | 4,64E-03 | Not-filtered |
| 64 | ENSCAFT00000005724 | CDK5RAP2 | 0,52 | 4,64E-03 | Not-filtered |
| 65 | ENSCAFT00000037497 | C31H21orf62 | -1,79 | 4,69E-03 | Not-filtered |
| 66 | ENSCAFT00000045655 | TDRD15 | -1,72 | 4,93E-03 | Not-filtered |
| 67 | ENSCAFT00000019265 | GPR3 | 0,77 | 4,96E-03 | Not-filtered |
| 68 | ENSCAFT00000048546 | PDE4C | -0,53 | 5,16E-03 | Not-filtered |
| 69 | ENSCAFT00000030960 | TTC34 | 0,68 | 5,81E-03 | Not-filtered |
| 70 | ENSCAFT00000029439 | MYO1F | 0,90 | 5,84E-03 | Not-filtered |
| 71 | ENSCAFT00000045052 | IGFBP6 | 1,00 | 6,34E-03 | Not-filtered |
| 72 | ENSCAFT00000048748 | CLCF1 | 0,83 | 6,73E-03 | Not-filtered |
| 73 | ENSCAFT00000026360 | EIF4E1B | -1,23 | 6,73E-03 | Not-filtered |
| 74 | ENSCAFT00000007670 | STAC | -0,53 | 6,86E-03 | Not-filtered |
| 75 | ENSCAFT00000046113 | LDLR | 0,52 | 6,94E-03 | Not-filtered |
| 76 | ENSCAFT00000011875 | THNSL2 | -0,41 | 6,94E-03 | Not-filtered |
| 77 | ENSCAFT00000013860 | SPI1 | 0,80 | 7,78E-03 | Not-filtered |
| 78 | ENSCAFT00000018652 |  | 0,48 | 7,78E-03 | Not-filtered |
| 79 | ENSCAFT00000038333 | DNAJB5 | 0,41 | 7,78E-03 | Not-filtered |
| 80 | ENSCAFT00000000961 | CCDC192 | -3,77 | 7,78E-03 | Not-filtered |
| 81 | ENSCAFT00000037518 |  | -1,73 | 7,78E-03 | Not-filtered |
| 82 | ENSCAFT00000000149 | CCBE1 | -0,77 | 7,78E-03 | Not-filtered |
| 83 | ENSCAFT00000048904 | RYR1 | -0,39 | 7,78E-03 | Not-filtered |
| 84 | ENSCAFT00000014735 | DCLRE1B | 0,29 | 8,22E-03 | Not-filtered |
| **85** | **ENSCAFT00000018984** |  | **0,28** | **8,32E-03** | **90^th^** |
| 86 | ENSCAFT00000011990 |  | -3,51 | 8,32E-03 | Not-filtered |
| 87 | ENSCAFT00000018504 | SCN7A | -1,23 | 8,32E-03 | Not-filtered |
| 88 | ENSCAFT00000031681 |  | 0,83 | 8,37E-03 | Not-filtered |
| 89 | ENSCAFT00000005356 | WHRN | 0,88 | 8,49E-03 | Not-filtered |
| 90 | ENSCAFT00000027791 | LDLR | 0,63 | 8,49E-03 | Not-filtered |
| 91 | ENSCAFT00000036732 | MAP2K3 | 0,58 | 8,79E-03 | Not-filtered |
| 92 | ENSCAFT00000020898 | NPL | -0,74 | 8,99E-03 | Not-filtered |
| 93 | ENSCAFT00000005230 | PLEK | 0,81 | 9,06E-03 | Not-filtered |
| 94 | ENSCAFT00000042917 | TMEM119 | 0,90 | 9,15E-03 | Not-filtered |
| 95 | ENSCAFT00000045786 |  | 0,75 | 9,25E-03 | Not-filtered |
| 96 | ENSCAFT00000035869 |  | -1,76 | 9,26E-03 | Not-filtered |
| 97 | ENSCAFT00000042894 | JHY | -1,03 | 9,26E-03 | Not-filtered |
| 98 | ENSCAFT00000025337 | RASAL3 | 0,80 | 1,02E-02 | Not-filtered |
| 99 | ENSCAFT00000046077 | SCN7A | -0,95 | 1,05E-02 | Not-filtered |
| 100 | ENSCAFT00000012335 | GRK7 | -0,96 | 1,09E-02 | Not-filtered |
| 101 | ENSCAFT00000007621 | PLK3 | 0,66 | 1,10E-02 | Not-filtered |
| **102** | **ENSCAFT00000013864** |  | **0,35** | **1,11E-02** | **97^th^** |
| 103 | ENSCAFT00000020626 | SDHC | 0,31 | 1,11E-02 | Not-filtered |
| 104 | ENSCAFT00000046956 | PTGFRN | 0,38 | 1,12E-02 | Not-filtered |
| 105 | ENSCAFT00000014439 |  | 0,35 | 1,12E-02 | Not-filtered |
| 106 | ENSCAFT00000027073 | MYCBPAP | 0,77 | 1,13E-02 | Not-filtered |
| 107 | ENSCAFT00000004532 |  | -2,00 | 1,16E-02 | Not-filtered |
| 108 | ENSCAFT00000043505 |  | 0,71 | 1,18E-02 | Not-filtered |
| 109 | ENSCAFT00000025850 | DNAH14 | -1,31 | 1,23E-02 | Not-filtered |
| 110 | ENSCAFT00000046505 | ETV4 | 0,75 | 1,23E-02 | Not-filtered |
| 111 | ENSCAFT00000035664 | RARA | 0,59 | 1,27E-02 | Not-filtered |
| 112 | ENSCAFT00000032475 | CES2 | 0,67 | 1,27E-02 | Not-filtered |
| **113** | **ENSCAFT00000043933** | **APOA1** | **0,79** | **1,32E-02** | **99^th^** |
| **114** | **ENSCAFT00000027074** | **CALR** | **0,35** | **1,38E-02** | **98^th^** |
| 115 | ENSCAFT00000045964 | COL18A1 | 0,95 | 1,38E-02 | Not-filtered |
| 116 | ENSCAFT00000013671 | STBD1 | -0,33 | 1,38E-02 | Not-filtered |
| 117 | ENSCAFT00000020769 | EPHB3 | 0,71 | 1,40E-02 | Not-filtered |
| 118 | ENSCAFT00000015123 | COX15 | -0,36 | 1,42E-02 | Not-filtered |
| 119 | ENSCAFT00000017152 | RBM20 | -0,58 | 1,43E-02 | Not-filtered |
| **120** | **ENSCAFT00000013297** |  | **0,63** | **1,44E-02** | **91^st^** |
| 121 | ENSCAFT00000049637 | SERPINB6 | 0,72 | 1,46E-02 | Not-filtered |
| 122 | ENSCAFT00000031982 | SLC25A25 | 0,48 | 1,47E-02 | Not-filtered |
| 123 | ENSCAFT00000005713 | CADPS2 | 0,48 | 1,48E-02 | Not-filtered |
| 124 | ENSCAFT00000006366 | TP53I3 | 0,68 | 1,53E-02 | Not-filtered |
| 125 | ENSCAFT00000047739 | DUSP1 | 0,65 | 1,57E-02 | Not-filtered |
| 126 | ENSCAFT00000045561 | FSCN3 | -0,85 | 1,57E-02 | Not-filtered |
| 127 | ENSCAFT00000007951 | DLEC1 | -0,61 | 1,57E-02 | Not-filtered |
| 128 | ENSCAFT00000030998 | CEP104 | -0,38 | 1,57E-02 | Not-filtered |
| 129 | ENSCAFT00000026053 | CAPN8 | 0,97 | 1,58E-02 | Not-filtered |
| 130 | ENSCAFT00000028872 | CAPN6 | -0,73 | 1,60E-02 | Not-filtered |
| **131** | **ENSCAFT00000011743** | **NR4A1** | **0,94** | **1,62E-02** | **93^rd^** |
| **132** | **ENSCAFT00000021280** | **ECHS1** | **0,71** | **1,63E-02** | **95^th^** |
| 133 | ENSCAFT00000031608 | CDT1 | 0,91 | 1,64E-02 | Not-filtered |
| **134** | **ENSCAFT00000026906** | **PER1** | **0,77** | **1,66E-02** | **94^th^** |
| 135 | ENSCAFT00000050055 | SHCBP1L | -1,44 | 1,67E-02 | Not-filtered |
| **136** | **ENSCAFT00000042866** | **RPRML** | **1,18** | **1,69E-02** | **95^th^** |
| 137 | ENSCAFT00000036880 | PTGFRN | 0,45 | 1,69E-02 | Not-filtered |
| 138 | ENSCAFT00000020415 | LINGO4 | 0,91 | 1,70E-02 | Not-filtered |
| 139 | ENSCAFT00000003238 | SULT1C4 | -0,87 | 1,75E-02 | Not-filtered |
| 140 | ENSCAFT00000023918 |  | -0,70 | 1,79E-02 | Not-filtered |
| 141 | ENSCAFT00000014831 | CCDC148 | -0,92 | 1,85E-02 | Not-filtered |
| 142 | ENSCAFT00000013794 | LMBR1L | 0,47 | 1,88E-02 | Not-filtered |
| 143 | ENSCAFT00000030761 |  | 0,38 | 1,88E-02 | Not-filtered |
| 144 | ENSCAFT00000049378 |  | -0,49 | 1,90E-02 | Not-filtered |
| 145 | ENSCAFT00000032461 | B3GNT9 | 1,07 | 1,98E-02 | Not-filtered |
| 146 | ENSCAFT00000037411 | PROB1 | 0,90 | 1,98E-02 | Not-filtered |
| 147 | ENSCAFT00000045154 | CD82 | 0,86 | 1,98E-02 | Not-filtered |
| 148 | ENSCAFT00000043092 | DCXR | 0,78 | 1,98E-02 | Not-filtered |
| 149 | ENSCAFT00000022898 |  | 0,75 | 1,98E-02 | Not-filtered |
| 150 | ENSCAFT00000032088 | IL34 | 0,72 | 1,98E-02 | Not-filtered |
| 151 | ENSCAFT00000020750 | EPHB3 | 0,68 | 1,98E-02 | Not-filtered |
| 152 | ENSCAFT00000014810 | SERPINB6 | 0,67 | 1,98E-02 | Not-filtered |
| 153 | ENSCAFT00000019411 | COL6A3 | -1,31 | 1,98E-02 | Not-filtered |
| 154 | ENSCAFT00000000277 | MC5R | -0,94 | 1,98E-02 | Not-filtered |
| 155 | ENSCAFT00000011022 | SLC38A9 | -0,58 | 1,98E-02 | Not-filtered |
| 156 | ENSCAFT00000021964 | IL16 | 0,63 | 2,02E-02 | Not-filtered |
| 157 | ENSCAFT00000011311 | RARA | 0,59 | 2,02E-02 | Not-filtered |
| 158 | ENSCAFT00000045379 | TTC34 | 0,69 | 2,03E-02 | Not-filtered |
| **159** | **ENSCAFT00000026939** | **PPP1R9B** | **0,76** | **2,03E-02** | **99^th^** |
| 160 | ENSCAFT00000045878 |  | 0,73 | 2,05E-02 | Not-filtered |
| 161 | ENSCAFT00000026036 | TNFRSF1B | 0,84 | 2,06E-02 | Not-filtered |
| 162 | ENSCAFT00000036157 | C3 | 0,66 | 2,16E-02 | Not-filtered |
| 163 | ENSCAFT00000013878 | COQ9 | 0,34 | 2,21E-02 | Not-filtered |
| 164 | ENSCAFT00000048568 | CXHXorf36 | 0,38 | 2,29E-02 | Not-filtered |
| 165 | ENSCAFT00000016750 | OASL | -0,54 | 2,33E-02 | Not-filtered |
| 166 | ENSCAFT00000020372 | IL10RA | 0,58 | 2,36E-02 | Not-filtered |
| 167 | ENSCAFT00000022337 | PEX5 | 0,40 | 2,36E-02 | Not-filtered |
| 168 | ENSCAFT00000017458 | SLC37A2 | 0,53 | 2,37E-02 | Not-filtered |
| 169 | ENSCAFT00000013687 |  | 0,48 | 2,37E-02 | Not-filtered |
| **170** | **ENSCAFT00000027951** | **FAXDC2** | **0,50** | **2,39E-02** | **94^th^** |
| 171 | ENSCAFT00000013858 | ADRA1A | 0,46 | 2,39E-02 | Not-filtered |
| 172 | ENSCAFT00000011115 | PRR13 | 0,41 | 2,41E-02 | Not-filtered |
| 173 | ENSCAFT00000024439 | RCSD1 | 0,41 | 2,47E-02 | Not-filtered |
| **174** | **ENSCAFT00000045771** |  | **1,31** | **2,48E-02** | **96^th^** |
| 175 | ENSCAFT00000046339 | IER5 | 0,68 | 2,48E-02 | Not-filtered |
| 176 | ENSCAFT00000045328 | FBXW2 | 0,21 | 2,48E-02 | Not-filtered |
| 177 | ENSCAFT00000019420 | COL6A3 | -1,32 | 2,48E-02 | Not-filtered |
| **178** | **ENSCAFT00000031805** | **IER5L** | **0,83** | **2,48E-02** | **94^th^** |
| 179 | ENSCAFT00000036487 | REEP6 | 0,78 | 2,48E-02 | Not-filtered |
| 180 | ENSCAFT00000028543 | MPI | 0,43 | 2,48E-02 | Not-filtered |
| 181 | ENSCAFT00000024605 | ZWINT | 1,46 | 2,52E-02 | Not-filtered |
| 182 | ENSCAFT00000026954 | TMEM219 | 0,41 | 2,52E-02 | Not-filtered |
| **183** | **ENSCAFT00000027559** | **SLC39A1** | **0,66** | **2,52E-02** | **90^th^** |
| 184 | ENSCAFT00000048192 | LTBR | 0,78 | 2,53E-02 | Not-filtered |
| **185** | **ENSCAFT00000029765** |  | **0,71** | **2,53E-02** | **99^th^** |
| 186 | ENSCAFT00000022090 | TOR3A | 0,42 | 2,53E-02 | Not-filtered |
| 187 | ENSCAFT00000023727 | RPL22L1 | 0,40 | 2,53E-02 | Not-filtered |
| 188 | ENSCAFT00000003268 | CD72 | 0,47 | 2,54E-02 | Not-filtered |
| 189 | ENSCAFT00000032002 | ENG | 0,71 | 2,58E-02 | Not-filtered |
| 190 | ENSCAFT00000042960 |  | 1,29 | 2,59E-02 | Not-filtered |
| **191** | **ENSCAFT00000014198** | **RAB11FIP5** | **0,72** | **2,59E-02** | **90^th^** |
| **192** | **ENSCAFT00000026249** |  | **0,84** | **2,61E-02** | **91^st^** |
| 193 | ENSCAFT00000007977 | MYD88 | 0,58 | 2,61E-02 | Not-filtered |
| 194 | ENSCAFT00000020742 | FUCA1 | 0,23 | 2,61E-02 | Not-filtered |
| 195 | ENSCAFT00000019426 | COL6A3 | -1,31 | 2,61E-02 | Not-filtered |
| 196 | ENSCAFT00000008181 | CSRNP1 | 0,84 | 2,62E-02 | Not-filtered |
| 197 | ENSCAFT00000029568 | C3 | 0,63 | 2,62E-02 | Not-filtered |
| 198 | ENSCAFT00000004941 | PABPC4 | 0,85 | 2,64E-02 | Not-filtered |
| **199** | **ENSCAFT00000000260** | **NAB2** | **0,72** | **2,65E-02** | **92^nd^** |
| 200 | ENSCAFT00000009541 | MYADML2 | 0,92 | 2,67E-02 | Not-filtered |
| 201 | ENSCAFT00000017456 | RGMA | 0,75 | 2,67E-02 | Not-filtered |
| 202 | ENSCAFT00000046370 |  | 0,65 | 2,67E-02 | Not-filtered |
| 203 | ENSCAFT00000031626 | CTU2 | 0,65 | 2,67E-02 | Not-filtered |
| **204** | **ENSCAFT00000028433** | **SEMA7A** | **0,75** | **2,69E-02** | **97^th^** |
| 205 | ENSCAFT00000020170 | MRGBP | 0,66 | 2,72E-02 | Not-filtered |
| 206 | ENSCAFT00000011842 | PEPD | 0,49 | 2,72E-02 | Not-filtered |
| **207** | **ENSCAFT00000001475** | **CYB5R3** | **0,77** | **2,74E-02** | **96^th^** |
| 208 | ENSCAFT00000047908 | CCDC116 | 0,77 | 2,76E-02 | Not-filtered |
| 209 | ENSCAFT00000048129 | SASH3 | 0,62 | 2,76E-02 | Not-filtered |
| 210 | ENSCAFT00000044057 |  | 0,59 | 2,76E-02 | Not-filtered |
| 211 | ENSCAFT00000045210 | TAS1R2 | 0,75 | 2,79E-02 | Not-filtered |
| 212 | ENSCAFT00000045709 | WNT1 | 0,68 | 2,79E-02 | Not-filtered |
| 213 | ENSCAFT00000018498 | MECR | 0,51 | 2,79E-02 | Not-filtered |
| 214 | ENSCAFT00000018820 | COL18A1 | 0,75 | 2,82E-02 | Not-filtered |
| 215 | ENSCAFT00000029463 | TCTEX1D1 | -0,68 | 2,82E-02 | Not-filtered |
| 216 | ENSCAFT00000031121 | HES2 | 0,96 | 2,83E-02 | Not-filtered |
| 217 | ENSCAFT00000047285 | TRIM17 | -1,02 | 2,83E-02 | Not-filtered |
| 218 | ENSCAFT00000038993 | TTC6 | -1,01 | 2,83E-02 | Not-filtered |
| 219 | ENSCAFT00000011362 |  | 1,22 | 2,85E-02 | Not-filtered |
| **220** | **ENSCAFT00000021138** | **APOA1** | **0,76** | **2,85E-02** | **99^th^** |
| 221 | ENSCAFT00000032517 | CMTM3 | 0,73 | 2,85E-02 | Not-filtered |
| 222 | ENSCAFT00000011777 | KCTD15 | 0,69 | 2,85E-02 | Not-filtered |
| 223 | ENSCAFT00000037777 | SRD5A1 | 0,55 | 2,85E-02 | Not-filtered |
| **224** | **ENSCAFT00000023243** | **TMBIM1** | **0,54** | **2,85E-02** | **93^rd^** |
| 225 | ENSCAFT00000030969 | TMEM203 | 0,52 | 2,85E-02 | Not-filtered |
| 226 | ENSCAFT00000009183 | SPATA24 | 0,51 | 2,85E-02 | Not-filtered |
| 227 | ENSCAFT00000049973 | ACO1 | 0,32 | 2,85E-02 | Not-filtered |
| 228 | ENSCAFT00000008481 | SPRY2 | 0,25 | 2,85E-02 | Not-filtered |
| **229** | **ENSCAFT00000028730** |  | **0,23** | **2,85E-02** | **95^th^** |
| 230 | ENSCAFT00000000209 | FECH | 0,20 | 2,85E-02 | Not-filtered |
| 231 | ENSCAFT00000006192 | MGAM | -1,74 | 2,85E-02 | Not-filtered |
| 232 | ENSCAFT00000032444 |  | -1,46 | 2,85E-02 | Not-filtered |
| 233 | ENSCAFT00000044050 | UOX | -0,65 | 2,85E-02 | Not-filtered |
| 234 | ENSCAFT00000011576 | HTR7 | -0,41 | 2,85E-02 | Not-filtered |
| **235** | **ENSCAFT00000038047** | **OXLD1** | **0,65** | **2,85E-02** | **92^nd^** |
| 236 | ENSCAFT00000017780 | RBM20 | -0,53 | 2,85E-02 | Not-filtered |
| 237 | ENSCAFT00000018791 | RHOD | 0,44 | 2,85E-02 | Not-filtered |
| 238 | ENSCAFT00000047799 | SHD | 0,86 | 2,85E-02 | Not-filtered |
| 239 | ENSCAFT00000038326 | SQLE | 0,29 | 2,85E-02 | Not-filtered |
| 240 | ENSCAFT00000000090 | ITGA7 | 0,47 | 2,88E-02 | Not-filtered |
| 241 | ENSCAFT00000021287 | MAP3K11 | 0,72 | 2,91E-02 | Not-filtered |
| 242 | ENSCAFT00000048304 |  | 1,04 | 2,91E-02 | Not-filtered |
| **243** | **ENSCAFT00000030085** | **PLIN3** | **0,99** | **2,91E-02** | **92^nd^** |
| 244 | ENSCAFT00000023505 | TSSK1B | 0,82 | 2,91E-02 | Not-filtered |
| 245 | ENSCAFT00000022217 | ATP2A3 | 0,78 | 2,91E-02 | Not-filtered |
| 246 | ENSCAFT00000010557 | ARID5A | 0,75 | 2,91E-02 | Not-filtered |
| 247 | ENSCAFT00000030728 | PKMYT1 | 0,71 | 2,91E-02 | Not-filtered |
| 248 | ENSCAFT00000027675 | NSMCE1 | 0,39 | 2,91E-02 | Not-filtered |
| **249** | **ENSCAFT00000006939** |  | **0,60** | **2,93E-02** | **91^st^** |
| 250 | ENSCAFT00000046137 | SPIDR | 0,30 | 2,98E-02 | Not-filtered |
| 251 | ENSCAFT00000011244 | DEPP1 | 0,91 | 3,02E-02 | Not-filtered |
| 252 | ENSCAFT00000000389 | MAP7 | -0,74 | 3,02E-02 | Not-filtered |
| **253** | **ENSCAFT00000032429** | **ZDHHC1** | **0,67** | **3,03E-02** | **93^rd^** |
| 254 | ENSCAFT00000049248 | OTOS | 1,67 | 3,05E-02 | Not-filtered |
| 255 | ENSCAFT00000002252 | FGD2 | 0,52 | 3,11E-02 | Not-filtered |
| 256 | ENSCAFT00000039695 | SCLY | 0,72 | 3,11E-02 | Not-filtered |
| 257 | ENSCAFT00000031098 | TMEM141 | 0,60 | 3,18E-02 | Not-filtered |
| **258** | **ENSCAFT00000027334** |  | **0,70** | **3,26E-02** | **94^th^** |
| 259 | ENSCAFT00000017511 | HEPACAM | 0,57 | 3,26E-02 | Not-filtered |
| 260 | ENSCAFT00000046690 | SRA1 | 0,31 | 3,27E-02 | Not-filtered |
| 261 | ENSCAFT00000043352 | PROKR1 | -1,08 | 3,27E-02 | Not-filtered |
| 262 | ENSCAFT00000038598 | DXO | 0,53 | 3,28E-02 | Not-filtered |
| 263 | ENSCAFT00000031678 | PLPP7 | 0,69 | 3,28E-02 | Not-filtered |
| 264 | ENSCAFT00000026637 | SH3PXD2B | 0,54 | 3,35E-02 | Not-filtered |
| 265 | ENSCAFT00000004116 | TNNI3 | 0,72 | 3,36E-02 | Not-filtered |
| 266 | ENSCAFT00000009197 | TMEM173 | 0,54 | 3,37E-02 | Not-filtered |
| **267** | **ENSCAFT00000003950** | **SLC2A1** | **0,52** | **3,37E-02** | **92^nd^** |
| 268 | ENSCAFT00000023813 | LGALS3 | 0,43 | 3,37E-02 | Not-filtered |
| 269 | ENSCAFT00000011570 |  | 0,39 | 3,37E-02 | Not-filtered |
| 270 | ENSCAFT00000043190 | FBLN7 | 0,38 | 3,37E-02 | Not-filtered |
| 271 | ENSCAFT00000024118 |  | 0,36 | 3,37E-02 | Not-filtered |
| 272 | ENSCAFT00000017160 | TNNI1 | 1,10 | 3,39E-02 | Not-filtered |
| **273** | **ENSCAFT00000048841** | **CD99L2** | **0,70** | **3,39E-02** | **97th** |
| 274 | ENSCAFT00000046384 | ENG | 0,66 | 3,39E-02 | Not-filtered |
| 275 | ENSCAFT00000014976 | CHAC1 | 0,63 | 3,39E-02 | Not-filtered |
| 276 | ENSCAFT00000026288 | SMOC1 | 0,57 | 3,39E-02 | Not-filtered |
| **277** | **ENSCAFT00000047941** |  | **0,42** | **3,39E-02** | **93^rd^** |
| 278 | ENSCAFT00000007430 | AGBL5 | 0,37 | 3,39E-02 | Not-filtered |
| 279 | ENSCAFT00000015117 | ABI3BP | -1,40 | 3,39E-02 | Not-filtered |
| 280 | ENSCAFT00000014836 | CCDC148 | -0,81 | 3,39E-02 | Not-filtered |
| 281 | ENSCAFT00000009407 | ZIC5 | -0,57 | 3,39E-02 | Not-filtered |
| 282 | ENSCAFT00000015504 | PDZD7 | -0,47 | 3,39E-02 | Not-filtered |
| 283 | ENSCAFT00000028352 | ISLR | 0,81 | 3,47E-02 | Not-filtered |
| 284 | ENSCAFT00000024881 | ACTA2 | 0,71 | 3,49E-02 | Not-filtered |
| 285 | ENSCAFT00000002392 | TSPAN33 | 0,43 | 3,51E-02 | Not-filtered |
| 286 | ENSCAFT00000043744 | MESP2 | 0,66 | 3,53E-02 | Not-filtered |
| **287** | **ENSCAFT00000046278** | **CYB5R3** | **0,73** | **3,54E-02** | **94^th^** |
| 288 | ENSCAFT00000007733 | TSEN54 | 0,57 | 3,54E-02 | Not-filtered |
| 289 | ENSCAFT00000015914 | CBLN1 | 0,52 | 3,54E-02 | Not-filtered |
| **290** | **ENSCAFT00000020662** | **CYB561** | **0,60** | **3,55E-02** | **93^rd^** |
| 291 | ENSCAFT00000022385 | BCOR | 0,38 | 3,57E-02 | Not-filtered |
| **292** | **ENSCAFT00000048276** |  | **0,71** | **3,63E-02** | **90^th^** |
| 293 | ENSCAFT00000015425 | TSHB | -2,83 | 3,63E-02 | Not-filtered |
| 294 | ENSCAFT00000015161 | KCNE2 | -0,67 | 3,63E-02 | Not-filtered |
| **295** | **ENSCAFT00000031049** | **MIDN** | **0,76** | **3,64E-02** | **95^th^** |
| 296 | ENSCAFT00000042966 | CCDC116 | 0,73 | 3,64E-02 | Not-filtered |
| 297 | ENSCAFT00000048895 | EPS15L1 | 0,27 | 3,64E-02 | Not-filtered |
| 298 | ENSCAFT00000039155 | INSC | 1,04 | 3,65E-02 | Not-filtered |
| 299 | ENSCAFT00000025901 | CNIH3 | 0,81 | 3,65E-02 | Not-filtered |
| 300 | ENSCAFT00000036972 | BCAS4 | 0,80 | 3,65E-02 | Not-filtered |
| **301** | **ENSCAFT00000047634** | **RAB11FIP5** | **0,69** | **3,65E-02** | **92^nd^** |
| 302 | ENSCAFT00000006437 | TBXAS1 | 0,54 | 3,65E-02 | Not-filtered |
| 303 | ENSCAFT00000003334 | FHL2 | 0,40 | 3,65E-02 | Not-filtered |
| 304 | ENSCAFT00000000329 | CDO1 | 0,31 | 3,65E-02 | Not-filtered |
| 305 | ENSCAFT00000045689 | TBC1D7 | 0,25 | 3,65E-02 | Not-filtered |
| 306 | ENSCAFT00000028047 | SERPINA3 | -1,03 | 3,65E-02 | Not-filtered |
| 307 | ENSCAFT00000039388 |  | -0,44 | 3,65E-02 | Not-filtered |
| 308 | ENSCAFT00000016952 | HYAL2 | 0,68 | 3,66E-02 | Not-filtered |
| **309** | **ENSCAFT00000009467** | **RFTN1** | **0,61** | **3,67E-02** | **90^th^** |
| **310** | **ENSCAFT00000009662** |  | **0,47** | **3,68E-02** | **90^th^** |
| **311** | **ENSCAFT00000030879** | **PRXL2B** | **0,80** | **3,73E-02** | **91^st^** |
| 312 | ENSCAFT00000036196 | CTSH | 0,53 | 3,77E-02 | Not-filtered |
| 313 | ENSCAFT00000016466 | CYB5R1 | 0,47 | 3,77E-02 | Not-filtered |
| 314 | ENSCAFT00000008331 | MYO7A | 0,39 | 3,77E-02 | Not-filtered |
| 315 | ENSCAFT00000043486 | RBM38 | 0,66 | 3,79E-02 | Not-filtered |
| 316 | ENSCAFT00000005213 | C11H9orf43 | -1,43 | 3,80E-02 | Not-filtered |
| 317 | ENSCAFT00000027507 | ADAM19 | 0,42 | 3,80E-02 | Not-filtered |
| **318** | **ENSCAFT00000001996** | **LY6H** | **0,83** | **3,84E-02** | **90^th^** |
| **319** | **ENSCAFT00000026443** | **TBKBP1** | **0,71** | **3,84E-02** | **91^st^** |
| 320 | ENSCAFT00000037002 | TGFB1 | 0,60 | 3,84E-02 | Not-filtered |
| 321 | ENSCAFT00000046716 | SELENOS | 0,46 | 3,84E-02 | Not-filtered |
| **322** | **ENSCAFT00000000224** |  | **0,21** | **3,84E-02** | **99^th^** |
| 323 | ENSCAFT00000046176 |  | -1,03 | 3,84E-02 | Not-filtered |
| 324 | ENSCAFT00000010739 | ADGRD1 | -0,31 | 3,84E-02 | Not-filtered |
| 325 | ENSCAFT00000049636 | LYSMD4 | -0,22 | 3,84E-02 | Not-filtered |
| 326 | ENSCAFT00000008617 |  | 0,83 | 3,85E-02 | Not-filtered |
| **327** | **ENSCAFT00000046781** | **COTL1** | **0,69** | **3,86E-02** | **92^nd^** |
| 328 | ENSCAFT00000023821 | SLC2A1 | 0,50 | 3,86E-02 | Not-filtered |
| 329 | ENSCAFT00000035862 | ADAMTS14 | 0,75 | 3,88E-02 | Not-filtered |
| **330** | **ENSCAFT00000003651** | **IGFBP7** | **0,50** | **3,88E-02** | **95^th^** |
| 331 | ENSCAFT00000022824 |  | 0,46 | 3,88E-02 | Not-filtered |
| 332 | ENSCAFT00000049259 | EGR4 | 0,71 | 3,89E-02 | Not-filtered |
| 333 | ENSCAFT00000029080 | STXBP2 | 0,58 | 3,89E-02 | Not-filtered |
| 334 | ENSCAFT00000030441 | MYO1C | 0,45 | 3,89E-02 | Not-filtered |
| 335 | ENSCAFT00000001361 | PARVG | 0,67 | 3,90E-02 | Not-filtered |
| 336 | ENSCAFT00000044118 |  | 0,68 | 3,92E-02 | Not-filtered |
| 337 | ENSCAFT00000022956 | ZNF804A | 0,62 | 3,92E-02 | Not-filtered |
| 338 | ENSCAFT00000019651 | SLC37A4 | 0,54 | 3,92E-02 | Not-filtered |
| 339 | ENSCAFT00000017668 | ACACB | 0,39 | 3,92E-02 | Not-filtered |
| **340** | **ENSCAFT00000023142** | **ARL4D** | **0,55** | **3,93E-02** | **94^th^** |
| **341** | **ENSCAFT00000017502** | **SDHA** | **0,15** | **3,94E-02** | **96^th^** |
| 342 | ENSCAFT00000004509 | SPACA6 | -0,54 | 3,94E-02 | Not-filtered |
| 343 | ENSCAFT00000017565 | ITGB2 | 0,71 | 3,99E-02 | Not-filtered |
| 344 | ENSCAFT00000043567 | SOHLH1 | 1,26 | 4,04E-02 | Not-filtered |
| 345 | ENSCAFT00000029692 | SASH3 | 0,68 | 4,04E-02 | Not-filtered |
| 346 | ENSCAFT00000007662 | SLC25A19 | 0,67 | 4,04E-02 | Not-filtered |
| 347 | ENSCAFT00000048099 | MYCBPAP | 0,65 | 4,04E-02 | Not-filtered |
| 348 | ENSCAFT00000030431 | DAPK3 | 0,61 | 4,04E-02 | Not-filtered |
| 349 | ENSCAFT00000010812 | AMOTL2 | 0,45 | 4,04E-02 | Not-filtered |
| 350 | ENSCAFT00000013897 | BOLA3 | 0,38 | 4,04E-02 | Not-filtered |
| 351 | ENSCAFT00000036547 | AMOTL2 | 0,36 | 4,04E-02 | Not-filtered |
| 352 | ENSCAFT00000023491 | GULP1 | -0,52 | 4,04E-02 | Not-filtered |
| **353** | **ENSCAFT00000046003** | **CAPNS1** | **0,40** | **4,06E-02** | **92^nd^** |
| 354 | ENSCAFT00000049527 |  | 0,64 | 4,07E-02 | Not-filtered |
| 355 | ENSCAFT00000006254 | PLEKHA4 | 0,72 | 4,07E-02 | Not-filtered |
| **356** | **ENSCAFT00000006525** | **PODXL2** | **0,66** | **4,07E-02** | **94^th^** |
| **357** | **ENSCAFT00000031428** | **REXO4** | **0,60** | **4,07E-02** | **95^th^** |
| **358** | **ENSCAFT00000048442** | **PODXL2** | **0,67** | **4,07E-02** | **94^th^** |
| 359 | ENSCAFT00000024558 | TAS1R2 | 0,66 | 4,07E-02 | Not-filtered |
| **360** | **ENSCAFT00000017006** | **ASL** | **0,55** | **4,07E-02** | **94^th^** |
| 361 | ENSCAFT00000037539 | FAM161A | -0,57 | 4,07E-02 | Not-filtered |
| 362 | ENSCAFT00000046651 | ABI3 | 0,72 | 4,08E-02 | Not-filtered |
| 363 | ENSCAFT00000043135 | LDB3 | 0,85 | 4,13E-02 | Not-filtered |
| 364 | ENSCAFT00000047389 |  | 0,84 | 4,13E-02 | Not-filtered |
| 365 | ENSCAFT00000025530 | MMRN2 | 0,82 | 4,13E-02 | Not-filtered |
| 366 | ENSCAFT00000015021 | CD82 | 0,78 | 4,13E-02 | Not-filtered |
| 367 | ENSCAFT00000014092 | TGM2 | 0,71 | 4,13E-02 | Not-filtered |
| **368** | **ENSCAFT00000009494** | **PCYT2** | **0,69** | **4,13E-02** | **91^st^** |
| 369 | ENSCAFT00000047372 | MAFF | 0,58 | 4,13E-02 | Not-filtered |
| 370 | ENSCAFT00000046369 | ROM1 | 0,57 | 4,13E-02 | Not-filtered |
| 371 | ENSCAFT00000035631 | MCHR1 | 0,43 | 4,13E-02 | Not-filtered |
| 372 | ENSCAFT00000019881 | PFKFB4 | 0,42 | 4,13E-02 | Not-filtered |
| **373** | **ENSCAFT00000018410** | **MSMO1** | **0,24** | **4,13E-02** | **91^st^** |
| 374 | ENSCAFT00000043610 | SPACA6 | -0,45 | 4,13E-02 | Not-filtered |
| 375 | ENSCAFT00000023304 | LAG3 | 0,63 | 4,16E-02 | Not-filtered |
| 376 | ENSCAFT00000025466 | ITPKB | 0,69 | 4,17E-02 | Not-filtered |
| 377 | ENSCAFT00000048272 | METTL7A | 0,44 | 4,18E-02 | Not-filtered |
| **378** | **ENSCAFT00000046422** | **ERF** | **0,74** | **4,21E-02** | **90^th^** |
| 379 | ENSCAFT00000023093 | SH3TC1 | 0,69 | 4,21E-02 | Not-filtered |
| 380 | ENSCAFT00000018146 | MAPKAPK2 | 0,58 | 4,21E-02 | Not-filtered |
| 381 | ENSCAFT00000032430 | TPPP3 | 0,58 | 4,21E-02 | Not-filtered |
| 382 | ENSCAFT00000003309 | RCL1 | 0,28 | 4,21E-02 | Not-filtered |
| 383 | ENSCAFT00000038030 |  | -1,30 | 4,21E-02 | Not-filtered |
| 384 | ENSCAFT00000011043 | FAHD2A | 0,52 | 4,22E-02 | Not-filtered |
| 385 | ENSCAFT00000000248 |  | 0,82 | 4,22E-02 | Not-filtered |
| 386 | ENSCAFT00000029863 |  | 0,48 | 4,22E-02 | Not-filtered |
| 387 | ENSCAFT00000013320 | AGA | -0,22 | 4,22E-02 | Not-filtered |
| **388** | **ENSCAFT00000029100** | **EVI5L** | **0,70** | **4,23E-02** | **94^th^** |
| 389 | ENSCAFT00000044344 | GSTZ1 | 0,45 | 4,23E-02 | Not-filtered |
| 390 | ENSCAFT00000047011 | LMAN2 | 0,32 | 4,23E-02 | Not-filtered |
| 391 | ENSCAFT00000011246 | GAPDHS | 0,86 | 4,24E-02 | Not-filtered |
| 392 | ENSCAFT00000015739 | POLL | 0,45 | 4,24E-02 | Not-filtered |
| 393 | ENSCAFT00000046504 |  | 1,10 | 4,25E-02 | Not-filtered |
| **394** | **ENSCAFT00000017753** | **NRGN** | **0,62** | **4,25E-02** | **99^th^** |
| **395** | **ENSCAFT00000026846** | **FDPS** | **0,38** | **4,25E-02** | **91^st^** |
| 396 | ENSCAFT00000042959 | ALDH8A1 | -0,86 | 4,25E-02 | Not-filtered |
| 397 | ENSCAFT00000009996 | CDC25B | -0,40 | 4,25E-02 | Not-filtered |
| **398** | **ENSCAFT00000028891** |  | **0,59** | **4,26E-02** | **95^th^** |
| **399** | **ENSCAFT00000007432** | **APOE** | **0,81** | **4,26E-02** | **99^th^** |
| 400 | ENSCAFT00000001503 | SERHL2 | 0,35 | 4,26E-02 | Not-filtered |
| 401 | ENSCAFT00000015110 | HSD17B13 | -0,74 | 4,26E-02 | Not-filtered |
| 402 | ENSCAFT00000044769 |  | -0,67 | 4,26E-02 | Not-filtered |
| **403** | **ENSCAFT00000043093** | **C1QB** | **0,73** | **4,28E-02** | **91^st^** |
| **404** | **ENSCAFT00000026481** | **BCAN** | **0,61** | **4,28E-02** | **97^th^** |
| **405** | **ENSCAFT00000037889** |  | **0,50** | **4,28E-02** | **94^th^** |
| 406 | ENSCAFT00000023625 | BCS1L | 0,43 | 4,28E-02 | Not-filtered |
| 407 | ENSCAFT00000000089 | ITGA7 | 0,43 | 4,30E-02 | Not-filtered |
| 408 | ENSCAFT00000019060 | PRLHR | 1,06 | 4,31E-02 | Not-filtered |
| 409 | ENSCAFT00000002101 | PDGFB | 0,55 | 4,31E-02 | Not-filtered |
| **410** | **ENSCAFT00000043363** | **OAZ2** | **0,53** | **4,31E-02** | **96^th^** |
| 411 | ENSCAFT00000016881 | ZMYND10 | 0,46 | 4,31E-02 | Not-filtered |
| **412** | **ENSCAFT00000026732** | **ATP5MC1** | **0,39** | **4,34E-02** | **94^th^** |
| 413 | ENSCAFT00000008171 | INSIG1 | 0,24 | 4,34E-02 | Not-filtered |
| 414 | ENSCAFT00000015805 | STMND1 | -0,54 | 4,34E-02 | Not-filtered |
| **415** | **ENSCAFT00000023619** | **MXD4** | **0,71** | **4,35E-02** | **91^st^** |
| 416 | ENSCAFT00000012296 | ABCB9 | 0,50 | 4,35E-02 | Not-filtered |
| 417 | ENSCAFT00000007368 | GEMIN7 | 0,47 | 4,35E-02 | Not-filtered |
| 418 | ENSCAFT00000029317 | RNF135 | 0,45 | 4,35E-02 | Not-filtered |
| 419 | ENSCAFT00000006898 | CCDC8 | 0,42 | 4,35E-02 | Not-filtered |
| 420 | ENSCAFT00000025069 | DPYSL3 | 0,40 | 4,35E-02 | Not-filtered |
| 421 | ENSCAFT00000020133 | SELENBP1 | 0,33 | 4,35E-02 | Not-filtered |
| 422 | ENSCAFT00000046911 | DCDC1 | -0,98 | 4,35E-02 | Not-filtered |
| **423** | **ENSCAFT00000004280** |  | **0,54** | **4,37E-02** | **98^th^** |
| **424** | **ENSCAFT00000028305** | **STOML1** | **0,68** | **4,40E-02** | **94^th^** |
| 425 | ENSCAFT00000003023 | MTERF1 | -0,78 | 4,40E-02 | Not-filtered |
| 426 | ENSCAFT00000043816 | TMC7 | -0,48 | 4,40E-02 | Not-filtered |
| 427 | ENSCAFT00000046420 |  | 1,94 | 4,42E-02 | Not-filtered |
| 428 | ENSCAFT00000048762 |  | 1,09 | 4,42E-02 | Not-filtered |
| 429 | ENSCAFT00000043336 | LY6H | 0,78 | 4,42E-02 | Not-filtered |
| 430 | ENSCAFT00000022144 | FAH | 0,77 | 4,42E-02 | Not-filtered |
| **431** | **ENSCAFT00000048751** | **C1QA** | **0,69** | **4,42E-02** | **93^rd^** |
| 432 | ENSCAFT00000025974 | ADGRE5 | 0,69 | 4,42E-02 | Not-filtered |
| **433** | **ENSCAFT00000049194** | **CST3** | **0,65** | **4,42E-02** | **98^th^** |
| 434 | ENSCAFT00000015524 | NEURL2 | 0,64 | 4,42E-02 | Not-filtered |
| 435 | ENSCAFT00000046320 | RCN3 | 0,63 | 4,42E-02 | Not-filtered |
| 436 | ENSCAFT00000009007 | GAA | 0,60 | 4,42E-02 | Not-filtered |
| 437 | ENSCAFT00000045724 | CTU2 | 0,58 | 4,42E-02 | Not-filtered |
| 438 | ENSCAFT00000049447 | CCDC102A | 0,56 | 4,42E-02 | Not-filtered |
| **439** | **ENSCAFT00000029401** | **COPRS** | **0,48** | **4,42E-02** | **91^st^** |
| 440 | ENSCAFT00000001756 | MCHR1 | 0,47 | 4,42E-02 | Not-filtered |
| 441 | ENSCAFT00000008744 | SLCO2B1 | 0,39 | 4,42E-02 | Not-filtered |
| 442 | ENSCAFT00000007647 | MRPS7 | 0,38 | 4,42E-02 | Not-filtered |
| 443 | ENSCAFT00000019165 |  | 0,22 | 4,42E-02 | Not-filtered |
| 444 | ENSCAFT00000001433 | TTLL1 | 0,16 | 4,42E-02 | Not-filtered |
| 445 | ENSCAFT00000021053 | NRP2 | -0,26 | 4,46E-02 | Not-filtered |
| **446** | **ENSCAFT00000028942** | **CSF1R** | **0,63** | **4,48E-02** | **91^st^** |
| **447** | **ENSCAFT00000025423** | **COQ8A** | **0,62** | **4,48E-02** | **91^st^** |
| 448 | ENSCAFT00000047462 | MRC2 | 0,58 | 4,48E-02 | Not-filtered |
| **449** | **ENSCAFT00000004761** | **EMC10** | **0,48** | **4,48E-02** | **94^th^** |
| **450** | **ENSCAFT00000026830** | **LTBP2** | **2,58** | **4,49E-02** | **91^st^** |
| 451 | ENSCAFT00000002835 | SHISA3 | 1,62 | 4,49E-02 | Not-filtered |
| **452** | **ENSCAFT00000021217** | **APOD** | **1,08** | **4,49E-02** | **99^th^** |
| 453 | ENSCAFT00000023419 | CHST7 | 0,83 | 4,49E-02 | Not-filtered |
| 454 | ENSCAFT00000047446 | SELPLG | 0,77 | 4,49E-02 | Not-filtered |
| 455 | ENSCAFT00000039763 | TRIB3 | 0,75 | 4,49E-02 | Not-filtered |
| **456** | **ENSCAFT00000008918** | **LGALS3BP** | **0,70** | **4,49E-02** | **95^th^** |
| 457 | ENSCAFT00000022913 |  | 0,63 | 4,49E-02 | Not-filtered |
| 458 | ENSCAFT00000023281 | AOC3 | 0,61 | 4,49E-02 | Not-filtered |
| 459 | ENSCAFT00000000764 | TCF19 | 0,61 | 4,49E-02 | Not-filtered |
| 460 | ENSCAFT00000025475 | EPHA2 | 0,59 | 4,49E-02 | Not-filtered |
| **461** | **ENSCAFT00000019526** |  | **0,57** | **4,49E-02** | **91^st^** |
| 462 | ENSCAFT00000044681 | ARFGEF3 | 0,55 | 4,49E-02 | Not-filtered |
| **463** | **ENSCAFT00000007975** | **ACAA1** | **0,51** | **4,49E-02** | **93^rd^** |
| 464 | ENSCAFT00000019042 | DIRC2 | 0,24 | 4,49E-02 | Not-filtered |
| 465 | ENSCAFT00000028044 | SERPINA3 | -1,12 | 4,49E-02 | Not-filtered |
| 466 | ENSCAFT00000011466 | ZC3H8 | -0,40 | 4,49E-02 | Not-filtered |
| 467 | ENSCAFT00000030284 | HSPB11 | -0,34 | 4,49E-02 | Not-filtered |
| 468 | ENSCAFT00000016183 | GUSB | 0,50 | 4,49E-02 | Not-filtered |
| **469** | **ENSCAFT00000014487** | **MT2A** | **0,49** | **4,49E-02** | **98^th^** |
| 470 | ENSCAFT00000038426 |  | 0,80 | 4,50E-02 | Not-filtered |
| 471 | ENSCAFT00000017630 | CD99 | 0,47 | 4,50E-02 | Not-filtered |
| 472 | ENSCAFT00000045007 |  | 0,36 | 4,50E-02 | Not-filtered |
| **473** | **ENSCAFT00000034820** | **MT-ND1** | **-0,42** | **4,50E-02** | **98^th^** |
| 474 | ENSCAFT00000049525 | PRAG1 | 0,65 | 4,51E-02 | Not-filtered |
| 475 | ENSCAFT00000038192 | GSDMD | 0,78 | 4,53E-02 | Not-filtered |
| 476 | ENSCAFT00000045553 | NPEPL1 | 0,67 | 4,53E-02 | Not-filtered |
| **477** | **ENSCAFT00000014635** | **CHRM4** | **0,63** | **4,53E-02** | **91^st^** |
| **478** | **ENSCAFT00000030343** | **SNRPB** | **0,56** | **4,53E-02** | **93^rd^** |
| 479 | ENSCAFT00000046897 | EFCAB2 | 0,43 | 4,53E-02 | Not-filtered |
| 480 | ENSCAFT00000050202 | TRIM62 | 0,40 | 4,53E-02 | Not-filtered |
| 481 | ENSCAFT00000014157 | TLL1 | -1,10 | 4,53E-02 | Not-filtered |
| 482 | ENSCAFT00000010537 | ZNF382 | -0,88 | 4,53E-02 | Not-filtered |
| 483 | ENSCAFT00000020156 | ARHGAP30 | 0,54 | 4,53E-02 | Not-filtered |
| 484 | ENSCAFT00000028142 | TBX2 | 0,67 | 4,54E-02 | Not-filtered |
| **485** | **ENSCAFT00000009073** | **SIRT2** | **0,61** | **4,54E-02** | **98^th^** |
| 486 | ENSCAFT00000027087 | PLEKHO2 | 0,61 | 4,54E-02 | Not-filtered |
| 487 | ENSCAFT00000048821 | ITIH3 | 1,34 | 4,54E-02 | Not-filtered |
| **488** | **ENSCAFT00000030037** | **DEXI** | **0,73** | **4,54E-02** | **98^th^** |
| 489 | ENSCAFT00000013571 | MMP15 | 0,63 | 4,54E-02 | Not-filtered |
| **490** | **ENSCAFT00000026233** | **GALNT16** | **0,57** | **4,54E-02** | **95^th^** |
| **491** | **ENSCAFT00000019198** | **MCAM** | **0,53** | **4,54E-02** | **94^th^** |
| 492 | ENSCAFT00000001476 | WDR46 | 0,47 | 4,54E-02 | Not-filtered |
| **493** | **ENSCAFT00000024053** | **CAPZB** | **0,47** | **4,54E-02** | **96^th^** |
| **494** | **ENSCAFT00000027758** | **SUPT4H1** | **0,38** | **4,54E-02** | **93^rd^** |
| 495 | ENSCAFT00000008258 | LCT | -0,55 | 4,54E-02 | Not-filtered |
| 496 | ENSCAFT00000000403 | IL20RA | -0,37 | 4,54E-02 | Not-filtered |
| 497 | ENSCAFT00000046931 | HDHD2 | -0,35 | 4,54E-02 | Not-filtered |
| 498 | ENSCAFT00000011649 | KRT71 | 1,02 | 4,54E-02 | Not-filtered |
| 499 | ENSCAFT00000013918 | WNT1 | 0,67 | 4,54E-02 | Not-filtered |
| **500** | **ENSCAFT00000030389** | **CD99L2** | **0,64** | **4,54E-02** | **96th** |
| 501 | ENSCAFT00000025515 | GPR173 | 0,49 | 4,54E-02 | Not-filtered |
| 502 | ENSCAFT00000039444 |  | 0,38 | 4,54E-02 | Not-filtered |
| 503 | ENSCAFT00000046451 | SULF1 | -0,73 | 4,54E-02 | Not-filtered |
| 504 | ENSCAFT00000016563 | DHCR7 | 0,60 | 4,54E-02 | Not-filtered |
| **505** | **ENSCAFT00000027989** | **ATP5F1A** | **0,26** | **4,54E-02** | **98^th^** |
| 506 | ENSCAFT00000050049 | DNASE2 | 0,59 | 4,57E-02 | Not-filtered |
| 507 | ENSCAFT00000002073 | SIL1 | 0,46 | 4,61E-02 | Not-filtered |
| **508** | **ENSCAFT00000026928** |  | **0,54** | **4,61E-02** | **98^th^** |
| 509 | ENSCAFT00000044103 | PKDCC | 0,45 | 4,61E-02 | Not-filtered |
| 510 | ENSCAFT00000008885 | EOMES | 1,19 | 4,61E-02 | Not-filtered |
| 511 | ENSCAFT00000023001 | ETV4 | 0,69 | 4,61E-02 | Not-filtered |
| 512 | ENSCAFT00000021197 | MRC2 | 0,61 | 4,61E-02 | Not-filtered |
| **513** | **ENSCAFT00000045076** | **CEND1** | **0,59** | **4,61E-02** | **99^th^** |
| 514 | ENSCAFT00000013448 | MOGS | 0,55 | 4,61E-02 | Not-filtered |
| **515** | **ENSCAFT00000013822** | **ADGRG1** | **0,54** | **4,61E-02** | **93^rd^** |
| 516 | ENSCAFT00000043384 |  | 0,53 | 4,61E-02 | Not-filtered |
| 517 | ENSCAFT00000044371 | LRCH4 | 0,52 | 4,61E-02 | Not-filtered |
| **518** | **ENSCAFT00000013340** | **SCARA3** | **0,51** | **4,61E-02** | **90^th^** |
| 519 | ENSCAFT00000004561 | ETFB | 0,44 | 4,61E-02 | Not-filtered |
| 520 | ENSCAFT00000024087 | CD9 | 0,42 | 4,61E-02 | Not-filtered |
| 521 | ENSCAFT00000027168 | CITED1 | 0,39 | 4,61E-02 | Not-filtered |
| 522 | ENSCAFT00000025961 | TCAP | 0,36 | 4,61E-02 | Not-filtered |
| 523 | ENSCAFT00000029024 | SLFN14 | -0,64 | 4,61E-02 | Not-filtered |
| 524 | ENSCAFT00000020162 | NPAS4 | 0,95 | 4,62E-02 | Not-filtered |
| 525 | ENSCAFT00000018997 | TMEM136 | -0,48 | 4,63E-02 | Not-filtered |
| 526 | ENSCAFT00000009489 | PCYT2 | 0,69 | 4,63E-02 | Not-filtered |
| 527 | ENSCAFT00000045894 |  | 1,16 | 4,63E-02 | Not-filtered |
| 528 | ENSCAFT00000006386 | SULT2B1 | 1,02 | 4,63E-02 | Not-filtered |
| 529 | ENSCAFT00000037740 |  | 0,83 | 4,63E-02 | Not-filtered |
| 530 | ENSCAFT00000009576 | DCXR | 0,75 | 4,63E-02 | Not-filtered |
| **531** | **ENSCAFT00000024555** | **WSCD1** | **0,71** | **4,63E-02** | **90^th^** |
| **532** | **ENSCAFT00000035428** | **IER2** | **0,70** | **4,63E-02** | **93^rd^** |
| **533** | **ENSCAFT00000026529** | **CPLX1** | **0,67** | **4,63E-02** | **99^th^** |
| 534 | ENSCAFT00000049456 | SLC39A3 | 0,67 | 4,63E-02 | Not-filtered |
| 535 | ENSCAFT00000018316 | HAPLN3 | 0,67 | 4,63E-02 | Not-filtered |
| 536 | ENSCAFT00000006533 |  | 0,67 | 4,63E-02 | Not-filtered |
| 537 | ENSCAFT00000026538 | SLC49A3 | 0,61 | 4,63E-02 | Not-filtered |
| **538** | **ENSCAFT00000027930** | **YIPF2** | **0,60** | **4,63E-02** | **92^nd^** |
| 539 | ENSCAFT00000012188 | C1H19orf12 | 0,59 | 4,63E-02 | Not-filtered |
| 540 | ENSCAFT00000026821 | FAM117A | 0,59 | 4,63E-02 | Not-filtered |
| 541 | ENSCAFT00000007121 | VASP | 0,57 | 4,63E-02 | Not-filtered |
| 542 | ENSCAFT00000011352 |  | 0,56 | 4,63E-02 | Not-filtered |
| 543 | ENSCAFT00000015230 | CHID1 | 0,55 | 4,63E-02 | Not-filtered |
| 544 | ENSCAFT00000044230 | RRP1 | 0,54 | 4,63E-02 | Not-filtered |
| 545 | ENSCAFT00000035983 | TTC32 | 0,45 | 4,63E-02 | Not-filtered |
| 546 | ENSCAFT00000000217 | RBMS2 | 0,40 | 4,63E-02 | Not-filtered |
| 547 | ENSCAFT00000021389 | UCK2 | 0,38 | 4,63E-02 | Not-filtered |
| **548** | **ENSCAFT00000020625** | **SDHC** | **0,26** | **4,63E-02** | **90^th^** |
| **549** | **ENSCAFT00000016300** |  | **0,21** | **4,63E-02** | **92^nd^** |
| 550 | ENSCAFT00000000234 | CCDC68 | -1,06 | 4,63E-02 | Not-filtered |
| 551 | ENSCAFT00000047464 | CYP1B1 | -0,94 | 4,63E-02 | Not-filtered |
| 552 | ENSCAFT00000003375 |  | -0,87 | 4,63E-02 | Not-filtered |
| 553 | ENSCAFT00000037896 |  | -0,85 | 4,63E-02 | Not-filtered |
| 554 | ENSCAFT00000010394 | HPX | -0,55 | 4,63E-02 | Not-filtered |
| 555 | ENSCAFT00000002701 | BICRAL | -0,44 | 4,63E-02 | Not-filtered |
| 556 | ENSCAFT00000018299 | TARBP1 | -0,41 | 4,63E-02 | Not-filtered |
| **557** | **ENSCAFT00000039816** | **OAT** | **-0,29** | **4,63E-02** | **95^th^** |
| 558 | ENSCAFT00000029092 | RAD51D | 0,54 | 4,63E-02 | Not-filtered |
| **559** | **ENSCAFT00000007134** | **ME3** | **0,51** | **4,63E-02** | **96^th^** |
| 560 | ENSCAFT00000046733 | SRI | -0,23 | 4,63E-02 | Not-filtered |
| **561** | **ENSCAFT00000031352** | **BSG** | **0,65** | **4,64E-02** | **99^th^** |
| 562 | ENSCAFT00000023294 | SLC25A1 | 0,60 | 4,64E-02 | Not-filtered |
| 563 | ENSCAFT00000029653 | SLC46A1 | 0,48 | 4,64E-02 | Not-filtered |
| 564 | ENSCAFT00000026293 | ITGAL | 0,71 | 4,67E-02 | Not-filtered |
| **565** | **ENSCAFT00000009418** | **ASPSCR1** | **0,65** | **4,67E-02** | **96^th^** |
| 566 | ENSCAFT00000029224 |  | 1,11 | 4,67E-02 | Not-filtered |
| 567 | ENSCAFT00000044447 |  | 0,61 | 4,67E-02 | Not-filtered |
| 568 | ENSCAFT00000030888 | MOB3A | 0,64 | 4,67E-02 | Not-filtered |
| 569 | ENSCAFT00000030881 | PNPLA7 | 0,56 | 4,67E-02 | Not-filtered |
| 570 | ENSCAFT00000021177 | MRC2 | 0,61 | 4,72E-02 | Not-filtered |
| 571 | ENSCAFT00000048982 | CD101 | -0,71 | 4,72E-02 | Not-filtered |
| 572 | ENSCAFT00000019844 | CASTOR1 | 0,50 | 4,74E-02 | Not-filtered |
| 573 | ENSCAFT00000019653 | SCLY | 0,63 | 4,76E-02 | Not-filtered |
| 574 | ENSCAFT00000016804 | SELENOS | 0,45 | 4,76E-02 | Not-filtered |
| 575 | ENSCAFT00000020341 | TCN2 | 0,55 | 4,78E-02 | Not-filtered |
| 576 | ENSCAFT00000005428 | STK40 | 0,55 | 4,78E-02 | Not-filtered |
| 577 | ENSCAFT00000024761 | NWD1 | 0,36 | 4,78E-02 | Not-filtered |
| **578** | **ENSCAFT00000032066** | **SLC2A8** | **0,67** | **4,78E-02** | **92^nd^** |
| **579** | **ENSCAFT00000008608** | **SERTAD1** | **0,63** | **4,79E-02** | **92^nd^** |
| **580** | **ENSCAFT00000015047** | **GOT1** | **0,35** | **4,79E-02** | **98^th^** |
| 581 | ENSCAFT00000049200 | TRIB2 | 0,18 | 4,79E-02 | Not-filtered |
| 582 | ENSCAFT00000031228 |  | 0,78 | 4,80E-02 | Not-filtered |
| 583 | ENSCAFT00000045026 | COL17A1 | 0,62 | 4,80E-02 | Not-filtered |
| 584 | ENSCAFT00000046184 | CRTAC1 | 0,48 | 4,80E-02 | Not-filtered |
| 585 | ENSCAFT00000014206 | LYZF2 | 1,27 | 4,82E-02 | Not-filtered |
| 586 | ENSCAFT00000043197 |  | 0,53 | 4,82E-02 | Not-filtered |
| 587 | ENSCAFT00000000131 | DGKA | -0,19 | 4,83E-02 | Not-filtered |
| 588 | ENSCAFT00000044444 | LMO3 | -0,58 | 4,83E-02 | Not-filtered |
| **589** | **ENSCAFT00000022063** | **VGF** | **0,68** | **4,86E-02** | **98^th^** |
| **590** | **ENSCAFT00000046233** | **LTBP2** | **2,55** | **4,87E-02** | **91^st^** |
| 591 | ENSCAFT00000042954 | SOX17 | 0,81 | 4,87E-02 | Not-filtered |
| **592** | **ENSCAFT00000036976** | **EVI5L** | **0,67** | **4,87E-02** | **93^rd^** |
| **593** | **ENSCAFT00000045135** | **PPP1CA** | **0,61** | **4,87E-02** | **93^rd^** |
| 594 | ENSCAFT00000002463 | RECQL4 | 0,55 | 4,87E-02 | Not-filtered |
| **595** | **ENSCAFT00000023617** | **CHCHD1** | **0,51** | **4,87E-02** | **91^st^** |
| 596 | ENSCAFT00000015184 | DNMBP | -0,40 | 4,87E-02 | Not-filtered |
| **597** | **ENSCAFT00000021221** | **APOD** | **1,03** | **4,89E-02** | **99^th^** |
| **598** | **ENSCAFT00000017804** | **LRP10** | **0,75** | **4,89E-02** | **90^th^** |
| **599** | **ENSCAFT00000029276** | **SREBF1** | **0,68** | **4,89E-02** | **90^th^** |
| 600 | ENSCAFT00000044529 | BOP1 | 0,68 | 4,89E-02 | Not-filtered |
| 601 | ENSCAFT00000048512 | KIF26B | 0,67 | 4,89E-02 | Not-filtered |
| 602 | ENSCAFT00000024534 | TSPAN9 | 0,64 | 4,89E-02 | Not-filtered |
| **603** | **ENSCAFT00000048052** | **PODXL2** | **0,64** | **4,89E-02** | **94^th^** |
| **604** | **ENSCAFT00000004100** | **HSPBP1** | **0,64** | **4,89E-02** | **97^th^** |
| **605** | **ENSCAFT00000031601** | **PPP1R26** | **0,62** | **4,89E-02** | **96^th^** |
| **606** | **ENSCAFT00000022529** | **ACSBG1** | **0,61** | **4,89E-02** | **92^nd^** |
| **607** | **ENSCAFT00000043153** | **CLEC2L** | **0,59** | **4,89E-02** | **94^th^** |
| **608** | **ENSCAFT00000026430** | **CLTB** | **0,59** | **4,89E-02** | **98^th^** |
| **609** | **ENSCAFT00000004187** | **TTYH1** | **0,58** | **4,89E-02** | **98^th^** |
| 610 | ENSCAFT00000023140 | DNAJC4 | 0,56 | 4,89E-02 | Not-filtered |
| 611 | ENSCAFT00000020594 | NARF | 0,54 | 4,89E-02 | Not-filtered |
| 612 | ENSCAFT00000007824 | PROSER2 | 0,51 | 4,89E-02 | Not-filtered |
| **613** | **ENSCAFT00000019330** | **HYOU1** | **0,48** | **4,89E-02** | **95^th^** |
| 614 | ENSCAFT00000017479 | SLC37A2 | 0,47 | 4,89E-02 | Not-filtered |
| 615 | ENSCAFT00000008964 | LMCD1 | 0,43 | 4,89E-02 | Not-filtered |
| 616 | ENSCAFT00000031424 | DENND2D | 0,42 | 4,89E-02 | Not-filtered |
| 617 | ENSCAFT00000031667 | KLHDC4 | 0,37 | 4,89E-02 | Not-filtered |
| 618 | ENSCAFT00000016873 | HSF2BP | 0,24 | 4,89E-02 | Not-filtered |
| **619** | **ENSCAFT00000015156** |  | **0,23** | **4,89E-02** | **93^rd^** |
| 620 | ENSCAFT00000032485 | SLC44A5 | -0,69 | 4,89E-02 | Not-filtered |
| 621 | ENSCAFT00000010536 | LMNTD2 | -0,68 | 4,89E-02 | Not-filtered |
| 622 | ENSCAFT00000047719 | HTR7 | -0,44 | 4,89E-02 | Not-filtered |
| 623 | ENSCAFT00000010002 | CDC25B | -0,40 | 4,89E-02 | Not-filtered |
| 624 | ENSCAFT00000026258 | SUSD6 | 0,44 | 4,89E-02 | Not-filtered |
| 625 | ENSCAFT00000035327 | NR2E1 | -0,48 | 4,90E-02 | Not-filtered |
| 626 | ENSCAFT00000023033 | GNB1L | 0,59 | 4,90E-02 | Not-filtered |
| **627** | **ENSCAFT00000014776** | **CRTAC1** | **0,56** | **4,91E-02** | **90^th^** |
| **628** | **ENSCAFT00000023116** | **TPI1** | **0,35** | **4,94E-02** | **99^th^** |
| 629 | ENSCAFT00000046346 | PIGN | 0,28 | 4,94E-02 | Not-filtered |
| 630 | ENSCAFT00000048429 |  | 0,53 | 4,95E-02 | Not-filtered |
| 631 | ENSCAFT00000046568 | CEACAM23 | 0,83 | 4,95E-02 | Not-filtered |
| 632 | ENSCAFT00000002737 | CHRNA5 | 0,39 | 4,97E-02 | Not-filtered |
| 633 | ENSCAFT00000023141 | DNAJC4 | 0,50 | 4,98E-02 | Not-filtered |
| 634 | ENSCAFT00000016048 | HMGCS2 | 1,64 | 4,98E-02 | Not-filtered |
| 635 | ENSCAFT00000031156 | SSTR5 | 0,93 | 4,98E-02 | Not-filtered |
| 636 | ENSCAFT00000008126 |  | 0,72 | 4,98E-02 | Not-filtered |
| **637** | **ENSCAFT00000013911** | **DDN** | **0,67** | **4,98E-02** | **98^th^** |
| 638 | ENSCAFT00000011216 | AKIP1 | 0,61 | 4,98E-02 | Not-filtered |
| 639 | ENSCAFT00000046490 | FN3K | 0,48 | 4,98E-02 | Not-filtered |
| 640 | ENSCAFT00000029547 | TRIP10 | 0,46 | 4,98E-02 | Not-filtered |
| 641 | ENSCAFT00000011194 | ST5 | 0,39 | 4,98E-02 | Not-filtered |
| **642** | **ENSCAFT00000026747** | **TRAPPC1** | **0,34** | **4,98E-02** | **93^rd^** |
| 643 | ENSCAFT00000017922 | ZNF639 | -0,27 | 4,98E-02 | Not-filtered |
| 644 | ENSCAFT00000023373 | MACROD1 | 0,69 | 4,99E-02 | Not-filtered |
| 645 | ENSCAFT00000017596 | ROBO4 | 0,64 | 4,99E-02 | Not-filtered |
| 646 | ENSCAFT00000020445 | FCER1G | 0,54 | 4,99E-02 | Not-filtered |
| 647 | ENSCAFT00000002605 | IMPDH1 | 0,50 | 4,99E-02 | Not-filtered |
| 648 | ENSCAFT00000001193 | PPT2 | 0,45 | 4,99E-02 | Not-filtered |
| 649 | ENSCAFT00000043269 |  | 0,28 | 4,99E-02 | Not-filtered |
| **650** | **ENSCAFT00000002307** | **LGALS1** | **0,74** | **5,00E-02** | **92^nd^** |
| 651 | ENSCAFT00000015776 | ADCY7 | 0,37 | 5,00E-02 | Not-filtered |
|  | ENSCAFT00000005173 | SPRED2 |  |  | **Added 97^th^, 95^th^,90^th^** |
|  | ENSCAFT00000023941 | MED28 |  |  | **Added 97^th^, 95^th^,90^th^** |
|  | ENSCAFT00000046536 |  |  |  | **Added 97^th^, 95^th^,90^th^** |
|  | ENSCAFT00000019347 |  |  |  | **Added 97^th^, 95^th^,90^th^** |
|  | ENSCAFT00000035659 | TINAGL1 |  |  | **Added 97^th^, 95^th^,90^th^** |
|  | ENSCAFT00000020451 | TMEM50A |  |  | **Added 97^th^, 95^th^** |
|  | ENSCAFT00000026776 | PHOSPHO1 |  |  | **Added 97^th^, 95^th^,90^th^** |
|  | ENSCAFT00000046387 | FBXO17 |  |  | **Added 97^th^, 95^th^,90^th^** |
|  | ENSCAFT00000022306 | ACHE |  |  | **Added 97^th^, 95^th^** |
|  | ENSCAFT00000025894 | PKN1 |  |  | **Added 97^th^, 95^th^** |
|  | ENSCAFT00000046295 | CBX2 |  |  | **Added 97^th^, 95^th^,90^th^** |
|  | ENSCAFT00000028935 | LHFPL1 |  |  | **Added 97^th^, 95^th^,90^th^** |
|  | ENSCAFT00000005896 | PREP |  |  | **Added 97^th^, 95^th^,90^th^** |
|  | ENSCAFT00000012109 | ANKRD37 |  |  | **Added 97^th^, 95^th^,90^th^** |
|  | ENSCAFT00000019616 | HMBS |  |  | **Added 97^th^, 95^th^,90^th^** |
|  | ENSCAFT00000000818 | SNCAIP |  |  | **Added 97^th^, 95^th^,90^th^** |
|  | ENSCAFT00000048703 | DENND10 |  |  | **Added 97^th^, 95^th^,90^th^** |
|  | ENSCAFT00000017817 | UNC93B1 |  |  | **Added 97^th^, 95^th^,90^th^** |
|  | ENSCAFT00000023184 | TNS1 |  |  | **Added 97^th^, 95^th^,90^th^** |
|  | ENSCAFT00000002969 | CKAP4 |  |  | **Added 97^th^, 95^th^,90^th^** |
|  | ENSCAFT00000001416 | TSPO |  |  | **Added 97^th^, 95^th^,90^th^** |
|  | ENSCAFT00000001830 | TRAPPC9 |  |  | **Added 97^th^, 95^th^** |
|  | ENSCAFT00000030449 | PNMA6A |  |  | **Added 97^th^, 95^th^** |
|  | ENSCAFT00000032043 | NIBAN2 |  |  | **Added 97^th^, 95^th^,90^th^** |
|  | ENSCAFT00000009379 | SRA1 |  |  | **Added 97^th^, 95^th^,90^th^** |
|  | ENSCAFT00000047650 | ACOT8 |  |  | **Added 97^th^, 95^th^,90^th^** |
|  | ENSCAFT00000027145 | TOB1 |  |  | **Added 97^th^, 95^th^,90^th^** |
|  | ENSCAFT00000045343 | TARBP1 |  |  | **Added 97^th^, 95^th^,90^th^** |
|  | ENSCAFT00000018419 | PPP1CA |  |  | **Added 97^th^, 95^th^** |
|  | ENSCAFT00000003571 | SUSD3 |  |  | **Added 97^th^, 95^th^,90^th^** |
|  | ENSCAFT00000016049 | TEX264 |  |  | **Added 97^th^, 95^th^** |
|  | ENSCAFT00000038320 | RNH1 |  |  | **Added 97^th^** |
|  | ENSCAFT00000001412 | TTLL12 |  |  | **Added 97^th^, 95^th^** |
|  | ENSCAFT00000014664 | PI4K2A |  |  | **Added 97^th^, 95^th^,90^th^** |
|  | ENSCAFT00000016906 | CSTB |  |  | **Added 97^th^, 95^th^** |
|  | ENSCAFT00000009602 | ZNF268 |  |  | **Added 97^th^, 95^th^,90^th^** |
|  | ENSCAFT00000044467 | GIMD1 |  |  | **Added 97^th^, 95^th^,90^th^** |
|  | ENSCAFT00000023290 | CTDSP1 |  |  | **Added 97^th^, 95^th^,90^th^** |
|  | ENSCAFT00000021383 | DGKI |  |  | **Added 97^th^, 95^th^,90^th^** |
|  | ENSCAFT00000047216 |  |  |  | **Added 97^th^, 95^th^,90^th^** |
|  | ENSCAFT00000002360 | KCTD17 |  |  | **Added 97^th^** |
|  | ENSCAFT00000030093 | TICAM1 |  |  | **Added 97^th^, 95^th^,90^th^** |
|  | ENSCAFT00000031657 | NTNG2 |  |  | **Added 97^th^, 95^th^,90^th^** |
|  | ENSCAFT00000009186 | CHMP6 |  |  | **Added 97^th^** |
|  | ENSCAFT00000028660 | SNX33 |  |  | **Added 97^th^, 95^th^,90^th^** |
|  | ENSCAFT00000002363 | TST |  |  | **Added 97th,95th** |
|  | ENSCAFT00000007160 | FOSB |  |  | **Added 97^th^, 95^th^,90^th^** |
|  | ENSCAFT00000008360 | CAPN5 |  |  | **Added 97th,95th** |
|  | ENSCAFT00000014711 | GOLGA7B |  |  | **Added 97^th^** |
|  | ENSCAFT00000023163 | CPZ |  |  | **Added 97^th^, 95^th^,90^th^** |
|  | ENSCAFT00000023453 | HSPG2 |  |  | **Added 97^th^, 95^th^,90^th^** |
|  | ENSCAFT00000026936 | BORCS6 |  |  | **Added 97^th^, 95^th^,90^th^** |
|  | ENSCAFT00000029761 | CAPS |  |  | **Added 97th,95th** |
|  | ENSCAFT00000029769 | NRTN |  |  | **Added 97^th^, 95^th^,90^th^** |
|  | ENSCAFT00000035590 | NRTN |  |  | **Added 97^th^, 95^th^,90^th^** |
|  | ENSCAFT00000044415 | ADGRA2 |  |  | **Added 97^th^, 95^th^,90^th^** |
|  | ENSCAFT00000008800 | MTMR14 |  |  | **Added 97^th^, 95^th^,90^th^** |
|  | ENSCAFT00000013256 | DOK1 |  |  | **Added 97^th^, 95^th^,90^th^** |
|  | ENSCAFT00000014170 | PRADC1 |  |  | **Added 97^th^, 95^th^,90^th^** |
|  | ENSCAFT00000014432 | PLBD2 |  |  | **Added 97^th^** |
|  | ENSCAFT00000024336 | SLC38A5 |  |  | **Added 97^th^, 95^th^,90^th^** |
|  | ENSCAFT00000028320 | MRM1 |  |  | **Added 97^th^, 95^th^,90^th^** |
|  | ENSCAFT00000028651 | SIN3A |  |  | **Added 97^th^, 95^th^,90^th^** |
|  | ENSCAFT00000036757 | TSPAN13 |  |  | **Added 97th,95th** |
|  | ENSCAFT00000043367 | TRAPPC6A |  |  | **Added 97^th^, 95^th^,90^th^** |
|  | ENSCAFT00000043331 | NCKAP1L |  |  | **Added 97^th^, 95^th^,90^th^** |
|  | ENSCAFT00000031387 |  |  |  | **Added 97^th^, 95^th^,90^th^** |
|  | ENSCAFT00000019530 | CERS2 |  |  | **Added 97th,95th** |
|  | ENSCAFT00000019209 | CIB1 |  |  | **Added 97th,95th** |
|  | ENSCAFT00000030547 | BCAP31 |  |  | **Added 97^th^** |
|  | ENSCAFT00000025240 | RGR |  |  | **Added 97^th^, 95^th^,90^th^** |
|  | ENSCAFT00000000313 | SHMT2 |  |  | **Added 97^th^, 95^th^,90^th^** |
|  | ENSCAFT00000011116 | ETV2 |  |  | **Added 97th,95th** |
|  | ENSCAFT00000043906 | H2BC18 |  |  | **Added 97^th^, 95^th^,90^th^** |
|  | ENSCAFT00000044751 | TRIOBP |  |  | **Added 97^th^, 95^th^,90^th^** |
|  | ENSCAFT00000002342 | MFNG |  |  | **Added 97^th^, 95^th^,90^th^** |
|  | ENSCAFT00000005111 | COL17A1 |  |  | **Added 97^th^, 95^th^,90^th^** |
|  | ENSCAFT00000000694 | PPP1R18 |  |  | **Added 97^th^, 95^th^,90^th^** |
|  | ENSCAFT00000013323 | NCKAP5L |  |  | **Added 97^th^, 95^th^,90^th^** |
|  | ENSCAFT00000011300 | ID1 |  |  | **Added 97^th^, 95^th^,90^th^** |
|  | ENSCAFT00000030472 | SLC43A2 |  |  | **Added 97^th^, 95^th^,90^th^** |
|  | ENSCAFT00000002700 | TOM1 |  |  | **Added 97^th^, 95^th^,90^th^** |
|  | ENSCAFT00000010865 | PRAG1 |  |  | **Added 97^th^, 95^th^,90^th^** |
|  | ENSCAFT00000023210 | VAT1 |  |  | **Added 97^th^, 95^th^** |
|  | ENSCAFT00000043059 | SREBF1 |  |  | **Added 97^th^, 95^th^,90^th^** |
|  | ENSCAFT00000005467 | PDIA4 |  |  | **Added 97^th^, 95^th^** |
|  | ENSCAFT00000014352 | DDX54 |  |  | **Added 97^th^, 95^th^,90^th^** |
|  | ENSCAFT00000019028 | KIF7 |  |  | **Added 97^th^, 95^th^,90^th^** |
|  | ENSCAFT00000027222 | GSTZ1 |  |  | **Added 97^th^, 95^th^,90^th^** |
|  | ENSCAFT00000031817 | S1PR1 |  |  | **Added 97^th^, 95^th^,90^th^** |
|  | ENSCAFT00000025531 | MMRN2 |  |  | **Added 97^th^, 95^th^,90^th^** |
|  | ENSCAFT00000027110 | DNASE2 |  |  | **Added 97^th^, 95^th^,90^th^** |
|  | ENSCAFT00000019200 | DENND10 |  |  | **Added 97^th^, 95^th^,90^th^** |
|  | ENSCAFT00000014281 | HERPUD1 |  |  | **Added 97^th^, 95^th^,90^th^** |
|  | ENSCAFT00000002231 | SOX10 |  |  | **Added 95^th^** |
|  | ENSCAFT00000048807 | KIF19 |  |  | **Added 97^th^, 95^th^,90^th^** |
|  | ENSCAFT00000005648 | ASTN2 |  |  | **Added 97^th^, 95^th^,90^th^** |
|  | ENSCAFT00000019797 | RPS6KA1 |  |  | **Added 97^th^, 95^th^,90^th^** |
|  | ENSCAFT00000026182 | PLOD1 |  |  | **Added 97^th^, 95^th^,90^th^** |
|  | ENSCAFT00000029231 | NUDT14 |  |  | **Added 95^th^** |
|  | ENSCAFT00000023969 | ITIH3 |  |  | **Added 97^th^, 95^th^,90^th^** |
|  | ENSCAFT00000030863 | ARHGAP4 |  |  | **Added 97^th^, 95^th^,90^th^** |
|  | ENSCAFT00000045788 | LMNA |  |  | **Added 97^th^, 95^th^,90^th^** |
|  | ENSCAFT00000026478 | EFNB3 |  |  | **Added 97^th^, 95^th^,90^th^** |
|  | ENSCAFT00000030174 | CARHSP1 |  |  | **Added 97^th^, 95^th^,90^th^** |
|  | ENSCAFT00000046454 | SCAMP4 |  |  | **Added 97^th^, 95^th^,90^th^** |
|  | ENSCAFT00000006555 | EHD2 |  |  | **Added 97^th^, 95^th^,90^th^** |
|  | ENSCAFT00000049535 | TARBP2 |  |  | **Added 97^th^, 95^th^,90^th^** |
|  | ENSCAFT00000002299 | TRIOBP |  |  | **Added 97^th^, 95^th^,90^th^** |
|  | ENSCAFT00000007452 | NECTIN2 |  |  | **Added 97^th^, 95^th^,90^th^** |
|  | ENSCAFT00000039596 | TSPAN11 |  |  | **Added 97^th^, 95^th^,90^th^** |
|  | ENSCAFT00000038921 | JPH2 |  |  | **Added 97^th^, 95^th^,90^th^** |
|  | ENSCAFT00000030963 | ARHGEF16 |  |  | **Added 97^th^, 95^th^,90^th^** |
|  | ENSCAFT00000023556 | MYOZ1 |  |  | **Added 97^th^, 95^th^,90^th^** |
|  | ENSCAFT00000050110 |  |  |  | **Added 97^th^, 95^th^,90^th^** |
|  | ENSCAFT00000020548 | OGFOD3 |  |  | **Added 97^th^, 95^th^,90^th^** |
|  | ENSCAFT00000019412 | TRIM26 |  |  | **Added 97^th^, 95^th^,90^th^** |
|  | ENSCAFT00000025645 | EPHX1 |  |  | **Added 97^th^, 95^th^,90^th^** |
|  | ENSCAFT00000025232 | CCDC22 |  |  | **Added 97^th^, 95^th^,90^th^** |
|  | ENSCAFT00000019049 | KREMEN1 |  |  | **Added 97^th^, 95^th^,90^th^** |
|  | ENSCAFT00000017710 | INPP5A |  |  | **Added 97^th^, 95^th^,90^th^** |
|  | ENSCAFT00000006272 | BCAT2 |  |  | **Added 97^th^, 95^th^,90^th^** |
|  | ENSCAFT00000047869 | PER2 |  |  | **Added 97^th^, 95^th^,90^th^** |
|  | ENSCAFT00000022273 | LARS2 |  |  | **Added 97^th^, 95^th^,90^th^** |
|  | ENSCAFT00000047037 | TPH1 |  |  | **Added 95^th^** |
|  | ENSCAFT00000003181 | SPATA18 |  |  | **Added 95^th^** |
|  | ENSCAFT00000028109 | EEF2K |  |  | **Added 95^th^** |
|  | ENSCAFT00000017991 | SELPLG |  |  | **Added 97^th^,95^th^,90^th^** |
|  | ENSCAFT00000011130 | AACS |  |  | **Added 95^th^** |
|  | ENSCAFT00000049304 |  |  |  | **Added 95^th^** |
|  | ENSCAFT00000014535 | CHST14 |  |  | **Added 97^th^, 95^th^,90^th^** |
|  | ENSCAFT00000045470 | HSPG2 |  |  | **Added 97^th^, 95^th^,90^th^** |
|  | ENSCAFT00000024856 | STK11IP |  |  | **Added 97^th^, 95^th^,90^th^** |
|  | ENSCAFT00000024922 | EVC2 |  |  | **Added 97^th^, 95^th^,90^th^** |
|  | ENSCAFT00000001166 | TNXB |  |  | **Added 97^th^, 95^th^,90^th^** |
|  | ENSCAFT00000046665 | TNXB |  |  | **Added 97^th^, 95^th^,90^th^** |
|  | ENSCAFT00000048328 | C17orf113 |  |  | **Added 97^th^, 95^th^,90^th^** |
|  | ENSCAFT00000030678 |  |  |  | **Added 95^th^** |
|  | ENSCAFT00000029702 | UNC119 |  |  | **Added 97^th^, 95^th^,90^th^** |
|  | ENSCAFT00000026151 | IL27RA |  |  | **Added 95^th^** |
|  | ENSCAFT00000005227 |  |  |  | **Added 95^th^** |
|  | ENSCAFT00000037329 | HCK |  |  | **Added 97^th^, 95^th^,90^th^** |
|  | ENSCAFT00000022140 | TM7SF2 |  |  | **Added 97^th^, 95^th^,90^th^** |
|  | ENSCAFT00000002028 | DEF6 |  |  | **Added 97^th^, 95^th^,90^th^** |
|  | ENSCAFT00000011738 | ATG101 |  |  | **Added 95^th^** |
|  | ENSCAFT00000007181 | NDUFB6 |  |  | **Added 95^th^** |
|  | ENSCAFT00000032147 | TAT |  |  | **Added 95^th^** |
|  | ENSCAFT00000008578 | GALNT14 |  |  | **Added 90^th^** |
|  | ENSCAFT00000022979 | PTPN6 |  |  | **Added 90^th^** |
|  | ENSCAFT00000019676 | CDC42SE1 |  |  | **Added 90^th^** |
|  | ENSCAFT00000022238 | LIMD1 |  |  | **Added 90^th^** |
