## Supplemental Table 5 for "On taming the effect of transcript level intra-condition count variation during differential expression analysis: a story of dogs, foxes and wolves"

|  | **Wolves and dogs** | | | **Aggressive and tame foxes** | | | |
| --- | --- | --- | --- | --- | --- | --- | --- |
| Percentile | Outliers | r^2^ | RMSE | Outliers | r^2^ |  | RMSE |
| NF | 281 | 0.74 | 0.81 | 20 | 0.49 |  | 1.12 |
| 99 | 262 | 0.76 | 0.79 | 20 | 0.5 |  | 1.11 |
| 98 | 249 | 0.77 | 0.77 | 19 | 0.51 |  | 1.11 |
| 97 | 233 | 0.78 | 0.76 | 18 | 0.52 |  | 1.1 |
| 96 | 219 | 0.78 | 0.76 | 18 | 0.53 |  | 1.1 |
| 95 | 218 | 0.79 | 0.75 | 17 | 0.54 |  | 1.09 |
| 94 | 205 | 0.8 | 0.74 | 16 | 0.55 |  | 1.09 |
| 93 | 198 | 0.8 | 0.73 | 15 | 0.55 |  | 1.09 |
| 92 | 186 | 0.81 | 0.73 | 15 | 0.56 |  | 1.08 |
| 91 | 179 | 0.81 | 0.72 | 14 | 0.57 |  | 1.08 |
| 90 | 172 | 0.82 | 0.72 | 14 | 0.58 |  | 1.07 |
| 85 | 141 | 0.83 | 0.7 | 13 | 0.61 |  | 1.05 |
| 80 | 123 | 0.85 | 0.68 | 13 | 0.65 |  | 1.03 |
| 75 | 118 | 0.86 | 0.67 | 14 | 0.68 |  | 1 |
| 70 | 102 | 0.87 | 0.65 | 15 | 0.72 |  | 0.97 |
