## Supplemental Table 6 for "On taming the effect of transcript level intra-condition count variation during differential expression analysis: a story of dogs, foxes and wolves"

| **Gene Family** | **Group** | **N genes** | **Gene name and log2FC value** |
| --- | --- | --- | --- |
| **Up regulated genes** | | | |
| Cholinergic receptor nicotinic alpha | **Shared** | 1 | **CHRNA5** (1.11 in dogs, 0.39 in tame foxes) |
| Squalene epoxidase | **Shared** | 1 | **SQLE** (0.55 in dogs, 0.29 in tame foxes) |
| Rho GTPase activating protein | **Shared** | 1 | **ARHGAP25** (0.86 in dogs, 0.70 in tame foxes) |
|  | Tame fox | 1 | ARHGAP30 (0.54) |
| Integrin alpha subunits | Dog | 3 | ITGA6 (1.26. 1.25); ITGA8 (1.14. 0.91); ITGAX (0.97) |
|  | Tame fox | 1 | ITGAL (0.71) |
|  | **Shared** | 1 | **ITGA7** (0.76 in dogs, 0.47 and 0.43 in tame foxes) |
| Myosin | Dog | 1 | MYO3A (1.12) |
|  | Tame fox | 2 | MYO1F (0.90); MYO1C (0.45) |
|  | **Shared** | 1 | **MYO7A** (0.82 in dogs, 0.39 in tame foxes) |
| Tribbles pseudokinase | Tame fox | 2 | TRIB1 (0.91); TRIB3 (0.75) |
|  | **Shared** | 1 | **TRIB2** (0.62 in dogs, 0.18 in tame foxes) |
| EF hand calcium binding | Dog | 1 | EFCAB1 (2.6) |
|  | Tame fox | 1 | EFCAB2 (0.43) |
| Transcription factor | Dog | 1 | TCF23 (2.05) |
|  | Tame fox | 1 | TCF19 (0.61) |
| Adhesion G protein-coupled receptors | Dog | 1 | ADGRG6 (1.46) |
|  | Tame fox | 1 | ADGRG1 (0.54) |
| Patatin Like Phospholipase Domain | Dog | 1 | PNPLA4 (1.42) |
|  | Tame fox | 1 | PNPLA7 (0.56) |
| SRY-box | Dog | 1 | SOX6 (1.27) |
|  | Tame fox | 1 | SOX17(0.81) |
| Hyaluronan and proteoglycan link protein | Dog | 1 | HAPLN1 (1.16) |
|  | Tame fox | 1 | HAPLN3 (0.67) |
| Serine/threonine kinase | Dog | 2 | STK17A (1.16. 1.14); STK32A (1.11) |
|  | Tame fox | 1 | STK40 (0.55) |
| Potassium channels | Dog | 1 | KCTD16 (0.98) |
|  | Tame fox | 1 | KCTD15 (0.69) |
| Podocalyxin like | Dog | 1 | PODXL (0.96. 0.85) |
|  | Tame fox | 1 | PODXL2 (0.77. 0.66. 0.64) |
| ATP binding cassette subfamily B | Dog | 1 | ABCB1 (0.93) |
|  | Tame fox | 1 | ABCB9 (0.50) |
| Zinc finger DHHC-type | Dog | 1 | ZDHHC15 (0.76) |
|  | Tame fox | 1 | ZDHHC1 (0.67) |
| Sushi domain | Dog | 1 | SUSD1 (0.69) |
|  | Tame fox | 1 | SUSD6 (0.44) |
| TBC1 domain family | Dog | 1 | TBC1D5 (0.55) |
|  | Tame fox | 1 | TBC1D7 (0.25) |
| Mitogen-activated protein kinase kinase kinases | Dog | 1 | MAP3K5 (0.51) |
|  | Tame fox | 1 | MAP3K11 (0.72) |
| Spermatogenesis Associated | Dog | 1 | SPATA5 (0.88) |
|  | Tame fox | 1 | SPATA24 (0.51) |
| **Down regulated genes** | | | |
| Stathmin domain | **Shared** | 1 | **STMND1** (-1.18 in dogs, -0.54 in tame foxes) |
| Oligoadenylate synthetase like | **Shared** | 1 | **OASL** (-0.40 in dogs, -0.54 in tame foxes) |
| Heat shock protein family B | Dog | 1 | HSPB8 (-0.69) |
|  | Tame fox | 1 | HSPB11 (-0.34) |
